# An atlas of white matter anatomy, its variability, and reproducibility based on Constrained Spherical Deconvolution of diffusion MRI

**DOI:** 10.1101/2021.10.13.464139

**Authors:** Ahmed Radwan, Stefan Sunaert, Kurt Schilling, Maxime Descoteaux, Bennett A. Landman, Mathieu Vandenbulcke, Tom Theys, Patrick Dupont, Louise Emsell

**Affiliations:** KU Leuven, Department of Imaging and pathology, Translational MRI, Leuven, Belgium; KU Leuven, Leuven Brain Institute, Department of Neurosciences, Leuven, Belgium; UZ Leuven, Department of Radiology, Leuven, Belgium; Department of Radiology and Radiological Sciences, Vanderbilt University Medical Center, Nashville, TN, USA; SCIL, Université de Sherbrooke, Quebec, Canada; Department of Electrical Engineering and Computer Science, Vanderbilt University, Nashville, TN, USA; KU Leuven, Department of Neurosciences, Neuropsychiatry, Leuven, Belgium; University Psychiatric Center (UPC) – Leuven, Geriatric Psychiatry, Leuven, Belgium; KU Leuven, Department of Neurosciences, Research Group Experimental Neurosurgery and Neuroanatomy, Leuven, Belgium; Department of Neurosurgery, University Hospitals Leuven, Leuven, Belgium; KU Leuven, Laboratory for Cognitive Neurology, Department of Neurosciences, Leuven, Belgium

## Abstract

Virtual dissection of white matter (WM) using diffusion MRI tractography is confounded by its poor reproducibility. Despite the increased adoption of advanced reconstruction models, early region-of-interest driven protocols based on diffusion tensor imaging (DTI) remain the dominant reference for virtual dissection protocols. Here we bridge this gap by providing a comprehensive description of typical WM anatomy reconstructed using a reproducible automated subject-specific parcellation-based approach based on probabilistic constrained-spherical deconvolution (CSD) tractography. We complement this with a WM template in MNI space comprising 68 bundles, including all associated anatomical tract selection labels and associated automated workflows. Additionally, we demonstrate bundle inter- and intra-subject variability using 40 (20 test-retest) datasets from the human connectome project (HCP) and 5 sessions with varying b-values and number of b-shells from the single-subject Multiple Acquisitions for Standardization of Structural Imaging Validation and Evaluation (MASSIVE) dataset. The most reliably reconstructed bundles were the whole pyramidal tracts, primary corticospinal tracts, whole superior longitudinal fasciculi, frontal, parietal and occipital segments of the corpus callosum and middle cerebellar peduncles. More variability was found in less dense bundles, e.g., the first segment of the superior longitudinal fasciculus, fornix, dentato-rubro-thalamic tract (DRTT), and premotor pyramidal tract. Using the DRTT as an example, we show that this variability can be reduced by using a higher number of seeding attempts. Overall inter-session similarity was high for HCP test-retest data (median weighted-dice = 0.963, stdev = 0.201 and IQR = 0.099). Compared to the HCP-template bundles there was a high level of agreement for the HCP test-retest data (median weighted-dice = 0.747, stdev = 0.220 and IQR = 0.277) and for the MASSIVE data (median weighted-dice = 0.767, stdev = 0.255 and IQR = 0.338). In summary, this WM atlas provides an overview of the capabilities and limitations of automated subject-specific probabilistic CSD tractography for mapping white matter fasciculi in healthy adults. It will be most useful in applications requiring a highly reproducible parcellation-based dissection protocol, as well as being an educational resource for applied neuroimaging and clinical professionals.

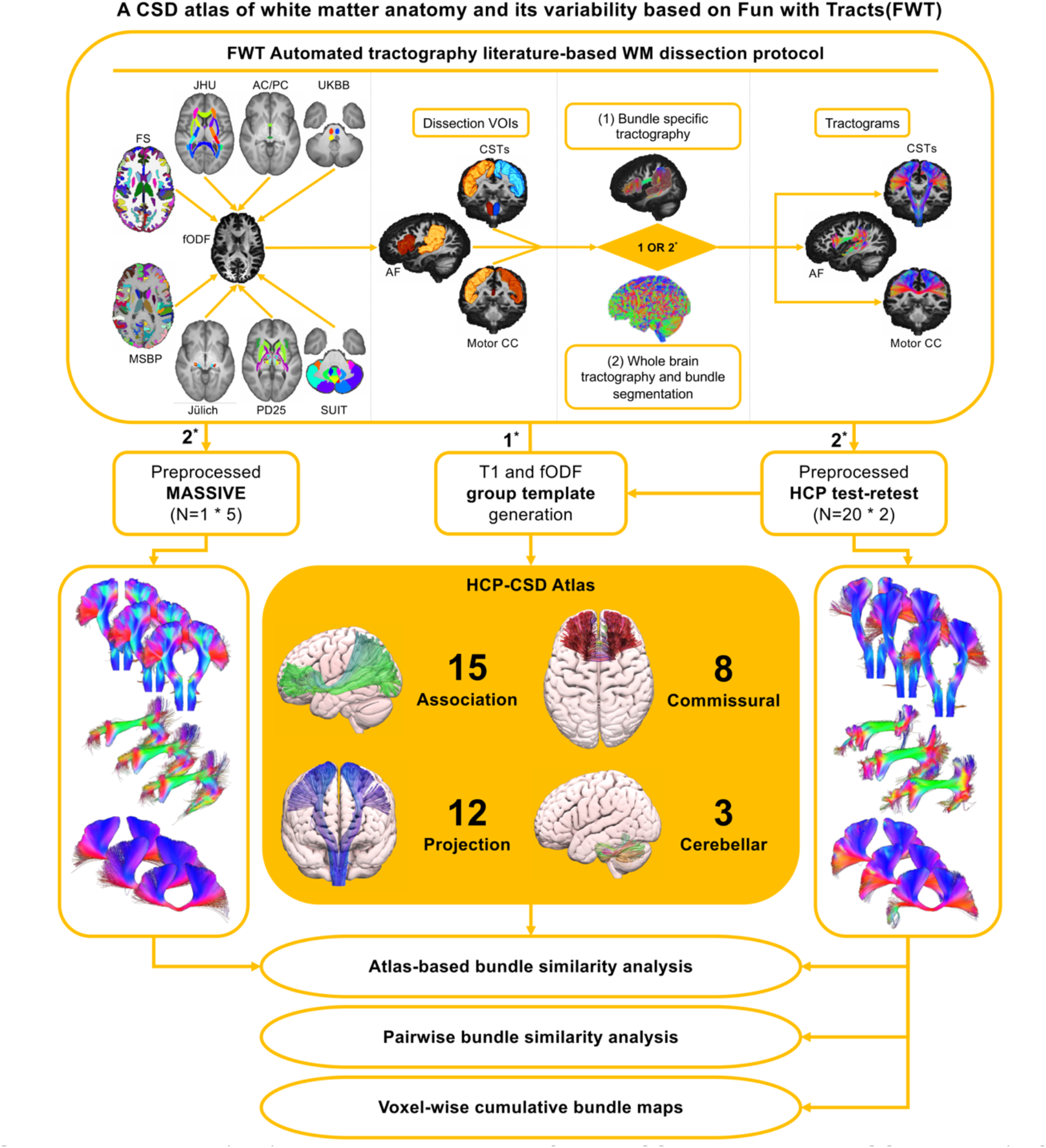
Graphical abstract. (Top) shows the FWT pipeline for both CSTs, AF, and motor CC bundles. (Left to right) show the required input structural parcellation maps and a priori atlases for FWT and the resulting virtual dissection include/exclude VOIs. FWT provides two approaches to virtual dissection: (1) is a bundle-specific approach where streamlines are only seeded for the bundle of interest, (2) is a whole brain tractography followed by streamlines segmentation, (top right) shows output tractograms. (Middle) Group-averaged T1 and fODF images are generated from the HCP test-retest data, and FWT is applied to generate the HCP-atlas using the bundle-specific approach (1*). FWT’s whole brain tracking and segmentation approach (2*) was applied to the HCP and MASSIVE dataset (right and left) and conducted model-based, and pair-wise similarity analyses and generated voxel-wise cumulative maps per bundle. FWT= Fun With Tracts, FS= FreeSurfer, MSBP= MultiScaleBrainParcellator, PD25= NIST Parkinson’s histological, JHU= John’s Hopkins university, Juelich= Juelich university histological atlas, AC/PC= anterior commissure/posterior commissure) UKBB= UK Biobank, SUIT (spatially unbiased cerebellar atlas template), dMRI= diffusion magnetic resonance imaging, CSD= constrained spherical deconvolution, fODF= fiber orientation distribution function, CST= corticospinal tract, AF= arcuate fasciculus, CC= corpus callosum, HCP= human connectome project, MASSIVE= Multiple acquisitions for standardization of structural imaging validation and evaluation.

## 1 Introduction

Characterising the macroscopic structural organisation of brain connectivity in vivo is central to understanding the human nervous system in health and disease. The advent of diffusion magnetic resonance imaging (dMRI) fiber tractography (FT) more than twenty years ago enabled significant progress in mapping major white matter (WM) fiber bundles described in anatomical and surgical literature. Initial work based on Diffusion Tensor Imaging (DTI) (Basser et al., 1994a, 1994b; Mori et al., 1999) drove the development of white matter dissection protocols (Catani, 2006; Mori et al., 2009, 2008; Wakana et al., 2007), which are still widely used today. This is because DTI is a simple and effective approach for reconstructing the core of large fasciculi (Catani et al., 2002; Stieltjes et al., 2001), data acquisition for DTI modelling requires a relatively short scan time (5-10 minutes) and standard pulse sequences are widely available owing to their regulatory approval for clinical practice. However, it is widely accepted that DTI suffers limitations that make it suboptimal for many tractography applications, particularly in a clinical setting (Farquharson et al., 2013; Mori and Tournier, 2014; Tournier et al., 2011). This has led to increased interest in more accurate and reliable approaches using high angular resolution imaging (HARDI) data (Bayrak et al., 2020; Bloy et al., 2012; Wasserthal et al., 2018; Yeh et al., 2018). At the same time, technological advances to accelerate data acquisition and reconstruction, such as multiband (Bouyagoub et al., 2020; Duan et al., 2015; Larkman et al., 2001; Moeller et al., 2010) and compressed sensing (Lustig et al., 2007b, 2007a) in combination with improved computational efficiency, will increase the adoption of advanced reconstruction models in applications traditionally reserved for DTI. This means that there is a need for updated reference material based on more representative virtual dissections using HARDI data acquired on 3T scanners.

Automated approaches relying on anatomical and orientational priors have been shown to considerably improve the accuracy of tract representations (Rheault et al., 2019) and address the poor reproducibility, and operator dependency that confound the manual virtual dissection process (Kreilkamp et al., 2019; Maffei et al., 2021; Soares et al., 2013). There are currently only a limited number of approaches that automate the virtual dissection process from start to finish (Warrington et al., 2020; Wasserthal et al., 2018; Yendiki et al., 2011). However, often the underlying dissection protocol is not explicitly detailed or the workflow is based on model bundles defined a priori.

This work addresses both the currently unmet need for an updated HARDI human white matter atlas that is relevant for clinical research studies, and a standardized, reproducible virtual dissection approach based on anatomical definitions. Using CSD due to its versatility and potential application to clinical data (Calamuneri et al., 2018; Toselli et al., 2017; Wilkins et al., 2015) we extend earlier virtual dissection protocols based on DTI by providing a descriptive summary of the normal WM anatomy of 68 fiber bundles reconstructed using probabilistic tractography. We complement our anatomical descriptions with an open-source group atlas in MNI space and automated subject-specific virtual dissection software, which we call “Fun with Tracts” (FWT) that incorporates all the anatomical inclusion and exclusion labels per bundle. As probabilistic tractography produces inherently variable results, we also demonstrate how this may vary within and between individuals using open-source test-retest datasets from the human connectome project (HCP) (Van Essen et al., 2012) and the Multiple acquisitions for standardization of structural imaging validation and evaluation (MASSIVE) dataset (Froeling et al., 2017). The result is a detailed and accessible reference for the virtual dissection of normal white matter anatomy using CSD tractography.

## 2 Material and methods

In summary, we generated a group average template from the HCP test-retest data, which was used to define the atlas of bundles based on a bundle-specific seeding and tracking approach. Whole-brain tractography followed by streamlines dissection was applied to the individual scans of the HCP test-retest and MASSIVE datasets. Both tracking approaches used the same virtual dissection protocol. Individual HCP test-retest results were used to demonstrate reproducibility in a group of subjects having two scans with the same acquisition parameters. Results from the MASSIVE datasets were used to demonstrate reproducibility in the same subject scanned multiple times with different acquisition parameters. Further details are provided below.

### 2.1 Imaging data

#### 2.1.1 HCP test-retest data

The test-retest HCP dataset (Van Essen et al., 2012) (https://db.humanconnectome.org/data/projects/HCP_Retest) consists of 2 scans acquired at 1 - 11 months apart using the same scanning protocol on the same 3-Tesla Siemens Skyra scanner (Siemens Healthineers, Erlangen, Germany) using a 32-channel phased array receive head coil. The HCP T1-weighted images were acquired using a 3D Magnetization Prepared Rapid Acquisition Gradient Echo (MPRAGE) pulse sequence with 0.7 mm isotropic voxels. The diffusion data was acquired with 90 directions per shell with b-values (1000, 2000, & 3000 s/mm^2^), and 1.25 mm isotropic voxels. We used the preprocessed (Andersson et al., 2003a; Andersson and Sotiropoulos, 2016, 2015; Glasser et al., 2013; Jenkinson et al., 2002) imaging data of 20 random subjects (10 male / 10 female) resulting in 40 scans (see supplementary table 1 for details).

#### 2.1.2 MASSIVE data

The MASSIVE dataset (Froeling et al., 2017) (http://www.massive-data.org), comprises multiple scans of the same healthy individual (female, 25 years old) using various b-values and diffusion sampling schemes. Imaging data was acquired on a 3-Tesla Philips Achieva scanner (Philips Healthcare, Best, The Netherlands) with an 8-channel phased array receive head coil. MASSIVE 3D T1-weighted images were acquired with 1 mm isotropic voxels, while dMRI images were acquired with 2.5 mm isotropic voxels and multiple shells with (0 - 9000 s/mm^2^) b-values. We used the (0 – 4000 s/mm^2^) b-shell data with the following b-values in s/mm^2^ and number of diffusion-weighting gradient directions respectively (b500 - 125, b1000 - 250, b2000 - 250, b3000 - 250, b4000 – 300). Five different sessions were generated using the preprocessed dMRI data acquired with an anterior-posterior (AP) phase-encoding axis and a negative gradient polarity to create 5 different sessions. Two sessions had multi-shell and three had single shell data with varying b-values, numbers of diffusion-weighted volumes and interleaved b0 volumes. Additionally, we used corresponding reversed-phase encoded b0 images for Echo-Planar Imaging (EPI) distortion correction in FSL (Jenkinson et al., 2012). We used the MASSIVE data to investigate FWT reproducibility in the same subject with different b-values, number of diffusion directions and b-shells. Table 1 lists the b-values and total number of volumes of all dMRI data used.

**Table 1:** Diffusion weightings and total number of volumes.

| **Table 1: Diffusion weightings and total number of volumes** |  |  |  |  |
| --- | --- | --- | --- | --- |
| Data/dMRI parameters |  | Number of volumes per shell | Diffusion weighting shell (b-values in s/mm<sup>2</sup>) | Total number of volumes |
| HCP test-retest data |  | 18, 90, 90, 90 | 0, 1000, 2000, 3000 | 288 |
| MASSIVE datasets | 1 | 60, 126, 251, 126 | 0, 500, 1000, 2000 | 563 |
|  | 2 | 60, 125, 251, 251 | 0, 2000, 3000, 4000 | 687 |
|  | 3 | 30, 251 | 0, 1000 | 281 |
|  | 4 | 30, 251 | 0, 2000 | 281 |
|  | 5 | 30, 251 | 0, 3000 | 281 |

### 2.2 Imaging data analysis

#### 2.2.1 Image preprocessing

All data was arranged in a Brain Imaging Data Structure (Gorgolewski et al., 2016) (BIDS) convention so that MultiScale Brain Parcellator (MSBP) (Tourbier et al., 2019) and FWT could run automatically. The T1-weighted images from both the HCP test-retest and the MASSIVE datasets were parcellated using FreeSurfer (Fischl, 2012) and the MSBP (Tourbier et al., 2019). We used MRTrix3 (Tournier et al., 2019) and FSL (Jenkinson et al., 2012) for dMRI preprocessing. The HCP test-retest data did not undergo any additional preprocessing. The MASSIVE diffusion data were preprocessed to remove any residual noise (Cordero-Grande et al., 2019; Veraart et al., 2016), motion/eddy and EPI related artefacts and distortions (Andersson et al., 2003a, 2003b; Andersson and Sotiropoulos, 2016; Bastiani et al., 2019; Skare and Bammer, 2009; Smith et al., 2004). fODF maps were generated using Multi-shell Multi-tissue CSD (Jeurissen et al., 2014) (MSMT-CSD) for all data, with three tissue types for multi-shell data and two tissue types of single-shell data.

#### 2.2.2 HCP test-retest group template creation

We created group averaged T1-weighted images and WM fODF maps from each individual HCP dataset in MRTrix3 (Tournier et al., 2019). These will be referred to throughout this work as the HCP-template for simplicity. The resulting averaged T1-weighted image was parcellated using FreeSurfer (Fischl, 2012) and MSBP (Tourbier et al., 2019).

#### 2.2.3 Virtual white matter dissection protocol

We describe 68 WM fiber bundles using a fully automated subject-specific whole brain tractography dissection workflow called “Fun With Tracts” (FWT) based on anatomical and neuroimaging literature definitions: 15 bilateral association bundles: The arcuate fasciculus, cingulum, fornix, frontal aslant tract, Inferior fronto-occipital fasciculus, inferior and middle longitudinal fasciculi, whole superior longitudinal fasciculus and its subcomponents, uncinate fasciculus, vertical occipital fasciculus. 8 commissural bundles (anterior commissure, and corpus callosum (in 7 segments). 12 bilateral projection bundles: the medial lemniscus, optic pathway, whole pyramidal tract and its subcomponents, and thalamic radiations). 3 bilateral cerebellar bundles (Dentato-rubro-thalamic tract, inferior and middle cerebellar peduncles). Definitions for each bundle are described in the results section and the inclusion/exclusion VOIs are provided in the supplementary methods.

#### 2.2.4 FWT

Figure 1 provides a schematic description of the employed methods in FWT. A detailed technical description including tracking and optimization parameters is provided in supplementary methods. The automated bash workflows can be found at (https://github.com/KUL-Radneuron/KUL_FWT.git). FWT employs automated tractography using MRTrix3 (Tournier et al., 2019) v3.0.2, which is constrained by a selection of grey matter and WM 3D VOIs/parcels used to create “inclusion” and “exclusion” areas, based on the neuroanatomical literature. These VOIs/parcels are obtained from FreeSurfer (Fischl, 2012) v6 (FS), the MSBP (Tourbier et al., 2019) v1.1.1, and several a priori atlases (see supplementary materials for details), along with custom VOIs manually defined in template space (the anterior commissure midline, and posterior commissure VOIs), and other custom VOIs that are generated by label propagation, (e.g., sub-segmentation of the periventricular white matter, temporal stem, insula, and superior temporal gyrus subcortical white matter). Streamlines filtering is done using ScilPy (Bore et al., 2021; “Scilpy documentation,” 2021 (Bore et al., 2021; “Scilpy documentation,” 2021)) v.1.1.0 and DIPY (Garyfallidis et al., 2014) tools and v1.3.0 for all bundles except the optic radiations for which we used the fiber-to-Bundle coherence (FBC) (Meesters et al., 2017; Portegies et al., 2015) tool in DIPY (Garyfallidis et al., 2014) v1.3.0. FWT can be used for individual bundle tractography or whole brain tractography and streamlines segmentation.

**Figure 1:**
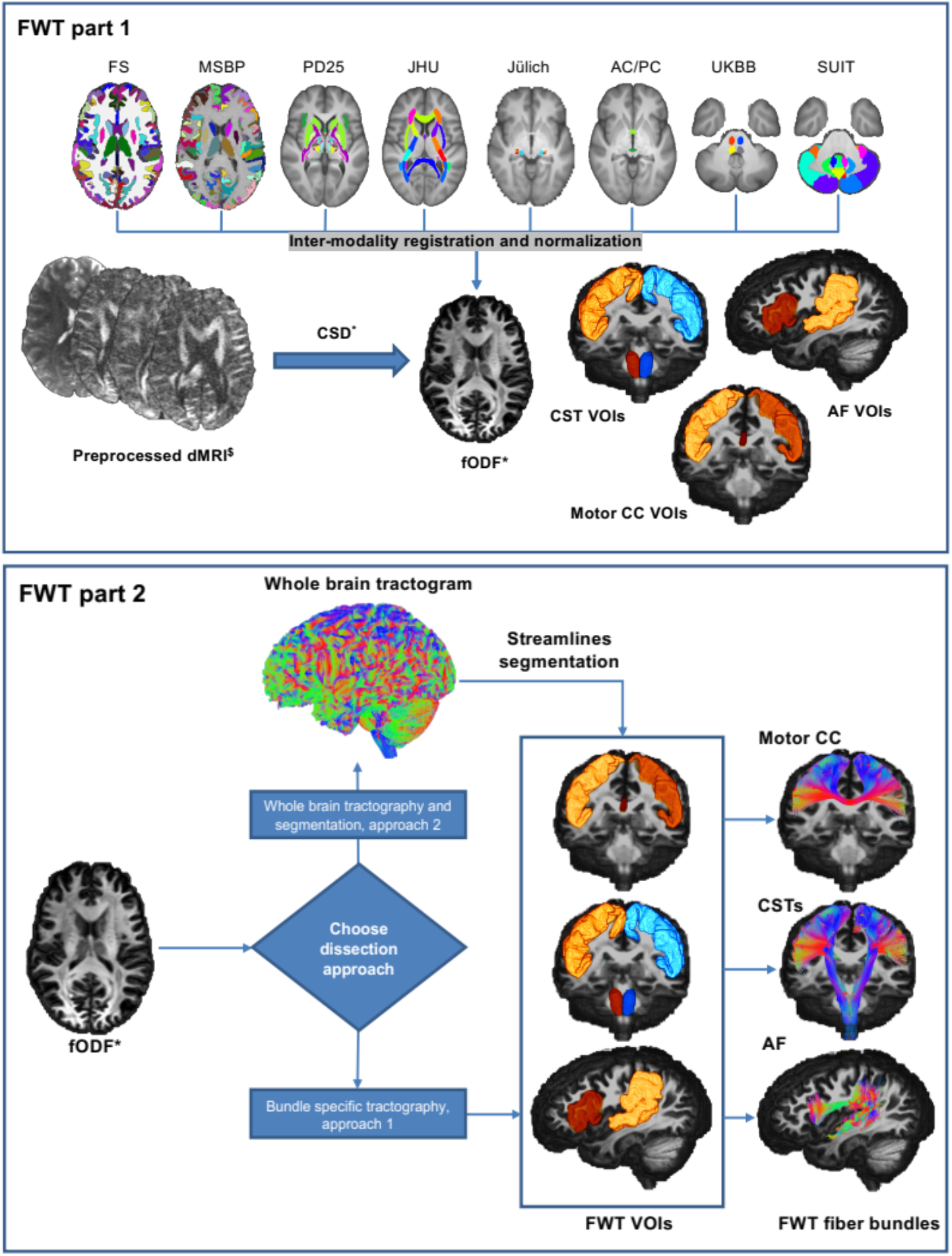
Graphical representation of the 2 parts of FWT for the corticospinal tracts (CSTs), arcuate fasciculi (AF), and motor corpus callosum (CC). FWT part 1 (top) requires as input preprocessed diffusion data, and FS recon-all output, and MSBP output. The script generates all VOIs used for virtual dissection by combining various anatomical VOIs from different parcellation maps and atlases. FWT part 2 (bottom) generates all tractograms from preprocessed diffusion data and VOIs created by FWT part 1, this script provides two approaches to virtual dissection; (1) Bundle specific and (2) Whole brain tractography followed by segmentation. FS = FreeSurfer, MSBP = MultiScale Brain Parcellator, PD25= NIST Parkinson’s histological, JHU= John’s Hopkins university, Juelich= Juelich university histological atlas, AC/PC= anterior commissure/posterior commissure, manually defined VOIsin template space, UKBB= UK Biobank, SUIT= spatially unbiased cerebellar atlas template, dMRI= diffusion magnetic resonance imaging, CSD= constrained spherical deconvolution, fODF = fiber orientation distribution function. $= preprocessed, should include correction for motion, Eddy currents, EPI distortion, imaging noise and bias. *= Other models e.g., DTI with FACT.

#### 2.2.5 Creation of inclusion and exclusion VOIs

We used the first part of the FWT workflow to automatically generate the inclusion and exclusion VOIs for all bundles in every dataset. This was applied to the two sessions of each subject in the HCP test-retest dataset individually, as well as to the different scan sessions of the preprocessed MASSIVE dataset. A similar workflow designed for group template T1 images was used on the HCP-template data. FWT automatically generates VOIs from the outputs of FS (Fischl, 2012) recon-all (FreeSurferWiki, 2020), MSBP (Tourbier et al., 2019), along with the UK BioBank (UKBB) volumetric atlas of fiber bundles (Miller et al., 2016), the spatially unbiased atlas template of the cerebellum and brainstem (Diedrichsen, 2006; Diedrichsen et al., 2011, 2009; Diedrichsen and Zotow, 2015) (SUIT), the Neuroimaging and surgical technologies (NIST) Parkinson’s disease histological atlas (Xiao et al., 2017, 2015, 2012) (PD25), and manually defined VOIs for the anterior and posterior commissures. Finally, all resulting anatomical VOIs are warped to diffusion space using ANTs (Avants et al., 2011), then combined to form bundle specific inclusion and exclusion VOIs. An equivalent workflow is available for group-averaged T1-weighted images.

#### 2.2.6 Individual tractography and creation of FWT-HCP template bundles

The second part of the FWT workflow provides the choice of two different whole brain WM virtual dissection approaches, both of which rely on the inclusion and exclusion VOIs created by the first part of FWT. The first approach employs bundle-specific seeding followed by streamline filtering and smoothing. The second generates a whole brain tractogram with 10 million streamlines by default followed by streamline dissection, then streamline filtering and smoothing. An equivalent workflow for fODF group averaged maps is also available generating whole brain tractograms with 20 million streamlines by default.

For this work we relied on bundle specific tractography (first approach) to generate the 68 HCP-template bundles, and whole brain tractography followed by bundle dissection (second approach) for all individual datasets. All template tractograms underwent visual quality assurance prior to further use.

#### 2.2.7 Tractogram reproducibility measures

Tractograms can be used to generate segmentation maps and as such can be evaluated using well-known similarity/dissimilarity measures such as Hausdorff distance, overlap measures, dice similarity coefficient (DSC), etc. While image similarity metrics tend to be largely reproducible, generally intuitive and exploit the pixel or voxel-wise composition of an image, the nature of streamline data presents a particular challenge for essentially binary-based similarity measures (Rheault et al., 2020). Streamline specific variants of these measures have recently been developed but have been shown to underestimate tractogram similarity (Rheault et al., 2020). We therefore used voxel-based weighted-dice similarity (wDSC) scores to simultaneously assess overlap and streamline density agreement per voxel in our inter-subject and intra-subject variability analyses (Cousineau et al., 2017). Additional similarity measures, e.g., DSC, density correlation (Rheault et al., 2020), volume overlap and overreach (Maier-Hein et al., 2017), and bundle adjacency (Garyfallidis et al., 2012) are provided in the supplementary material. In addition, we also generated voxel-wise cumulative maps for each bundle to evaluate inter-subject variability. The equations for the different measures used, can be found in supplementary materials.

### 2.3 Research questions and analyses

**(A)** First, we described and presented the atlas of bundles generated with our literature-based dissection protocol (FWT) for 68 WM fasciculi based on the HCP-template data using the bundle-specific seeding and tractography approach. Additionally, we highlight inter-session variability in the HCP test-retest data with respect to the template bundles by calculating wDSC scores. These were shown in radar plots and used to calculate single-rater agreement intra-class correlation (ICC) scores.

**(B)** To assess inter-session/intra-subject variability in the HCP individual datasets we applied the whole-brain tractography and bundle segmentation approach. We then calculated descriptive statistics and generated violin plots for the wDSC scores resulting from simple pairwise comparisons of the output from each session. This provided a measure of FWT reproducibility within and between subjects regardless of similarity to a model bundle. HCP inter-subject variability was evaluated by generating voxel-wise heat-maps of the summed binary masks of each bundle.

**(C)** FWT was applied to the MASSIVE data to test its performance on an independent dataset with different acquisition parameters and sampling scheme. Model-based wDSC bundle similarity scores, using the HCP-template bundles as references, were then calculated for every bundle from both the HCP and MASSIVE datasets. We computed wDSC descriptive statistics and generated violin plots for evaluation. This provided a framework for the comparison of the outputs from both datasets. Finally, we used single-rater agreement intra-class correlation scores to evaluate overall inter-session agreement.

## 3 Results

### 3.1 Qualitative bundle descriptions, template output and sample variability

Detailed qualitative descriptions of the protocol used to create the inclusion and exclusion VOIs for each fiber bundle along with the supporting literature are provided below. Demonstrative figures are also included for the tractography output of bundles reconstructed from the HCP-template. Bundles are grouped by type (Meynert, 1888), i.e., association, commissural, and projection bundles, with the cerebellar bundles grouped separately. To demonstrate inter-session variability in the source HCP test-retest data wDSC scores for each bundle compared to HCP-template bundles are shown on radar plots, and table 2 provides summary statistics for each bundle. We observe that some bundles are stable and reproducible, having a high median wDSC, a high minimum wDSC and a small Max.A.I.D. wDSC, while others are more variable and less stable, having lower median and minimum wDSC and larger Max.A.I.D. Overall single-score agreement ICC was 0.713 (upper-bound = 0.789, lower-bound = 0.637, P < 0.05).

**Table 2:** Summary weighted-dice statistics for all tracked bundles derived from comparing each bundle to the corresponding HCP-template bundle

| Bundles | N | Mean | Median | STDev | Min | Max | Max A.I.D. | IQR |
| --- | --- | --- | --- | --- | --- | --- | --- | --- |
| AF all L/R | 40/40 | 0.76/0.79 | 0.80/0.78 | 0.08/0.06 | 0.50/0.65 | 0.86/0.88 | 0.03/0.02 | 0.08/0.09 |
| CCing L/R | 40/40 | 0.86/0.74 | 0.86/0.71 | 0.04/0.08 | 0.77/0.59 | 0.95/0.89 | 0.03/0.02 | 0.05/0.11 |
| **TCing L/R** | 40/40 | 0.71/0.77 | 0.77/0.82 | 0.14/0.09 | 0.17/0.58 | 0.86/0.88 | 0.03/0.03 | 0.16/0.15 |
| **FAT L/R** | 40/40 | 0.73/0.74 | 0.80/0.79 | 0.16/0.12 | 0.25/0.52 | 0.89/0.9 | 0.11/0.13 | 0.19/0.21 |
| **Fx L/R** | 32/33 | 0.68/0.67 | 0.76/0.73 | 0.22/0.19 | 0.13/0.23 | 0.89/0.91 | 0.16/0.03 | 0.28/0.26 |
| **IFOF L/R** | 40/40 | 0.74/0.80 | 0.77/0.80 | 0.11/0.03 | 0.31/0.70 | 0.84/0.84 | 0.10/0.06 | 0.09/0.04 |
| **ILF L/R** | 40/40 | 0.70/0.71 | 0.7/0.71 | 0.03/0.04 | 0.64/0.61 | 0.76/0.78 | 0.05/0.04 | 0.04/0.04 |
| **MdLF L/R** | 40/40 | 0.73/0.53 | 0.79/0.56 | 0.17/0.15 | 0.16/0.05 | 0.89/0.79 | 0.52/0.35 | 0.14/0.18 |
| **SLF all L/R** | 40/40 | 0.74/0.8 | 0.78/0.81 | 0.11/0.06 | 0.35/0.67 | 0.89/0.88 | 0.28/0.18 | 0.09/0.10 |
| **SLF I L/R** | 40/40 | 0.61/0.14 | 0.67/0.11 | 0.19/0.14 | 0.10/0.00 | 0.83/0.58 | 0.10/0.05 | 0.16/0.14 |
| **SLF IId L/R** | 40/40 | 0.75/0.76 | 0.79/0.78 | 0.13/0.11 | 0.33/0.40 | 0.93/0.87 | 0.2/0.11 | 0.12/0.11 |
| **SLF IIv L/R** | 40/40 | 0.48/0.52 | 0.46/0.52 | 0.22/0.14 | 0.02/0.16 | 0.81/0.75 | 0.06/0.04 | 0.33/0.22 |
| **SLF III L/R** | 40/40 | 0.52/0.66 | 0.53/0.69 | 0.14/0.10 | 0.16/0.41 | 0.78/0.81 | 0.03/0.05 | 0.19/0.14 |
| **UF L/R** | 40/40 | 0.39/0.50 | 0.39/0.47 | 0.08/0.08 | 0.2/0.37 | 0.57/0.64 | 0.31/0.19 | 0.10/0.12 |
| **VOF L/R** | 40/40 | 0.62/0.28 | 0.7/0.28 | 0.19/0.09 | 0/0.08 | 0.80/0.44 | 0.27/0.36 | 0.20/0.15 |
| **Ant Comm** | 38 | 0.52 | 0.53 | 0.10 | 0.24 | 0.66 | 0.05 | 0.09 |
| **CC Motor** | 40 | 0.67 | 0.71 | 0.16 | 0.02 | 0.84 | 0.11 | 0.10 |
| **CC Occipital** | 40 | 0.80 | 0.82 | 0.06 | 0.64 | 0.89 | 0.49 | 0.05 |
| **CC Parietal** | 40 | 0.84 | 0.85 | 0.03 | 0.74 | 0.90 | 0.30 | 0.04 |
| **CC PMcandSMA** | 40 | 0.73 | 0.73 | 0.04 | 0.64 | 0.79 | 0.17 | 0.05 |
| **CC PreF** | 40 | 0.80 | 0.80 | 0.02 | 0.75 | 0.83 | 0.20 | 0.02 |
| **CC Sensory** | 36 | 0.72 | 0.79 | 0.19 | 0.05 | 0.90 | 0.14 | 0.20 |
| **CC Temporal** | 39 | 0.33 | 0.37 | 0.15 | 0.02 | 0.53 | 0.03 | 0.20 |
| **ML L/R** | 40/40 | 0.92/0.94 | 0.94/0.94 | 0.06/0.03 | 0.70/0.85 | 0.97/0.97 | 0.46/0.20 | 0.04/0.02 |
| **OR L/R** | 40/40 | 0.48/0.48 | 0.55/0.51 | 0.23/0.16 | 0.02/0.11 | 0.83/0.70 | 0.11/0.06 | 0.34/0.24 |
| **OR OL L/R** | 40/40 | 0.84/0.83 | 0.85/0.84 | 0.04/0.05 | 0.74/0.73 | 0.89/0.93 | 0.11/0.05 | 0.05/0.04 |
| **OT L/R** | 37/40 | 0.89/0.87 | 0.92/0.90 | 0.09/0.08 | 0.49/0.52 | 0.96/0.95 | 0.09/0.09 | 0.06/0.04 |
| **PyT all L/R** | 40/40 | 0.91/0.90 | 0.91/0.90 | 0.02/0.01 | 0.86/0.86 | 0.94/0.92 | 0.07/0.08 | 0.02/0.01 |
| **CST L/R** | 40/40 | 0.94/0.94 | 0.94/0.94 | 0.02/0.01 | 0.90/0.91 | 0.96/0.96 | 0.45/0.19 | 0.03/0.01 |
| **M1 CST L/R** | 40/40 | 0.94/0.94 | 0.94/0.95 | 0.02/0.01 | 0.87/0.91 | 0.96/0.96 | 0.59/0.28 | 0.03/0.02 |
| **PyT PMC L/R** | 39/35 | 0.22/0.20 | 0.22/0.21 | 0.04/0.05 | 0.11/0.11 | 0.31/0.33 | 0.03/0.06 | 0.05/0.06 |
| **PyT SMA L/R** | 40/40 | 0.88/0.9 | 0.90/0.91 | 0.06/0.03 | 0.62/0.79 | 0.93/0.95 | 0.40/0.25 | 0.05/0.04 |
| **ATR L/R** | 40/40 | 0.62/0.56 | 0.66/0.61 | 0.19/0.19 | 0.17/0.07 | 0.88/0.84 | 0.43/0.37 | 0.24/0.31 |
| **PaTR L/R** | 40/40 | 0.67/0.65 | 0.68/0.67 | 0.10/0.08 | 0.42/0.47 | 0.82/0.76 | 0.14/0.14 | 0.13/0.10 |
| **STR L/R** | 40/40 | 0.86/0.85 | 0.87/0.85 | 0.05/0.04 | 0.73/0.70 | 0.91/0.90 | 0.10/0.15 | 0.05/0.04 |
| **DRTT L/R** | 36/31 | 0.52/0.46 | 0.54/0.49 | 0.18/0.10 | 0.08/0.13 | 0.82/0.60 | 0.06/0.59 | 0.22/0.11 |
| **ICP L/R** | 40/40 | 0.53/0.48 | 0.57/0.52 | 0.18/0.16 | 0.04/0 | 0.76/0.73 | 0.22/0.40 | 0.15/0.16 |
| **MCP L/R** | 40/40 | 0.73/0.77 | 0.74/0.77 | 0.04/0.04 | 0.63/0.64 | 0.81/0.84 | 0.03/0.29 | 0.04/0.04 |

N = number of reconstructed bundles, STDev = standard deviation, Min = minimum, Max = maximum, Max. A.I.D. = maximum absolute intersession difference, IQR = interquartile range, L = left, R = right, AF all = Arcuate fasciculus (excluding the precentral gyrus), CCing = Cingulate cingulum, TCing = Temporal cingulum, FAT = frontal aslant tract, Fx = Fornix, IFOF = inferior fronto-occipital fasciculus, ILF = inferior longitudinal fasciculus, MdLF = middle longitudinal fasciculus, SLF all = superior longitudinal fasciculus (whole bundle), SLF I = 1<sup>st</sup> segment of SLF, SLF
IIId = dorsal subdivision of 2<sup>nd</sup> segment of SLF, SLF IIv = ventral subdivision of 2<sup>nd</sup> segment of SLF, SLF III = 3<sup>rd</sup> segment of SLF, UF = uncinate fasciculus, VOF = vertical occipital fasciculus, ML = medial lemniscus, OR = optic radiation, OR OL = optic radiation (using whole occipital lobe as cortical inclusion), OT = optic tract, PyT all = pyramidal tract (whole bundle), CST = corticospinal tract (primary motor and primary sensory as cortical inclusion), M1 CST = corticospinal tract (only primary motor as cortical inclusion), PyT PMC = Premotor cortex pyramidal tract, PyT SMA = supplementary motor area pyramidal tract, ATR = anterior thalamic radiation, PaTR = parietal thalamic radiation, STR = superior thalamic radiation, Ant Comm = anterior commissure, CC = corpus callosum, CC PMcandSMA = premotor and supplementary motor corpus callosum, CC PreF = prefrontal corpus callosum, DRT = dentato-rubro-thalamic tract, ICP = inferior cerebellar peduncle, MCP = middle cerebellar peduncle.

#### 3.1.1 Association fiber bundles *(in alphabetical order)*

##### 3.1.1.1 Arcuate Fasciculus (AF)

The AF is one of the main components of the dorsal language stream and constitutes a perisylvian fronto-temporal pathway consisting of a long ‘direct’ association fiber system connecting the ventral precentral and posterior portion of the inferior and middle frontal gyri with the middle and superior temporal gyri, and two shorter ‘in-direct’ bundles: (1) an anterior network connecting the supramarginal and superior temporal gyri with the precentral gyrus, and (2) a posterior network connecting the posterior middle temporal gyrus with the angular gyrus (Bain et al., 2019; Bernard et al., 2019; Catani, 2006; Chen et al., 2015; Fernández-Miranda et al., 2015; Wakana et al., 2007; Wang et al., 2016; Yeh et al., 2018), see figure 2. The AF in our data was generated for all datasets on both sides.

**Figure 2:**
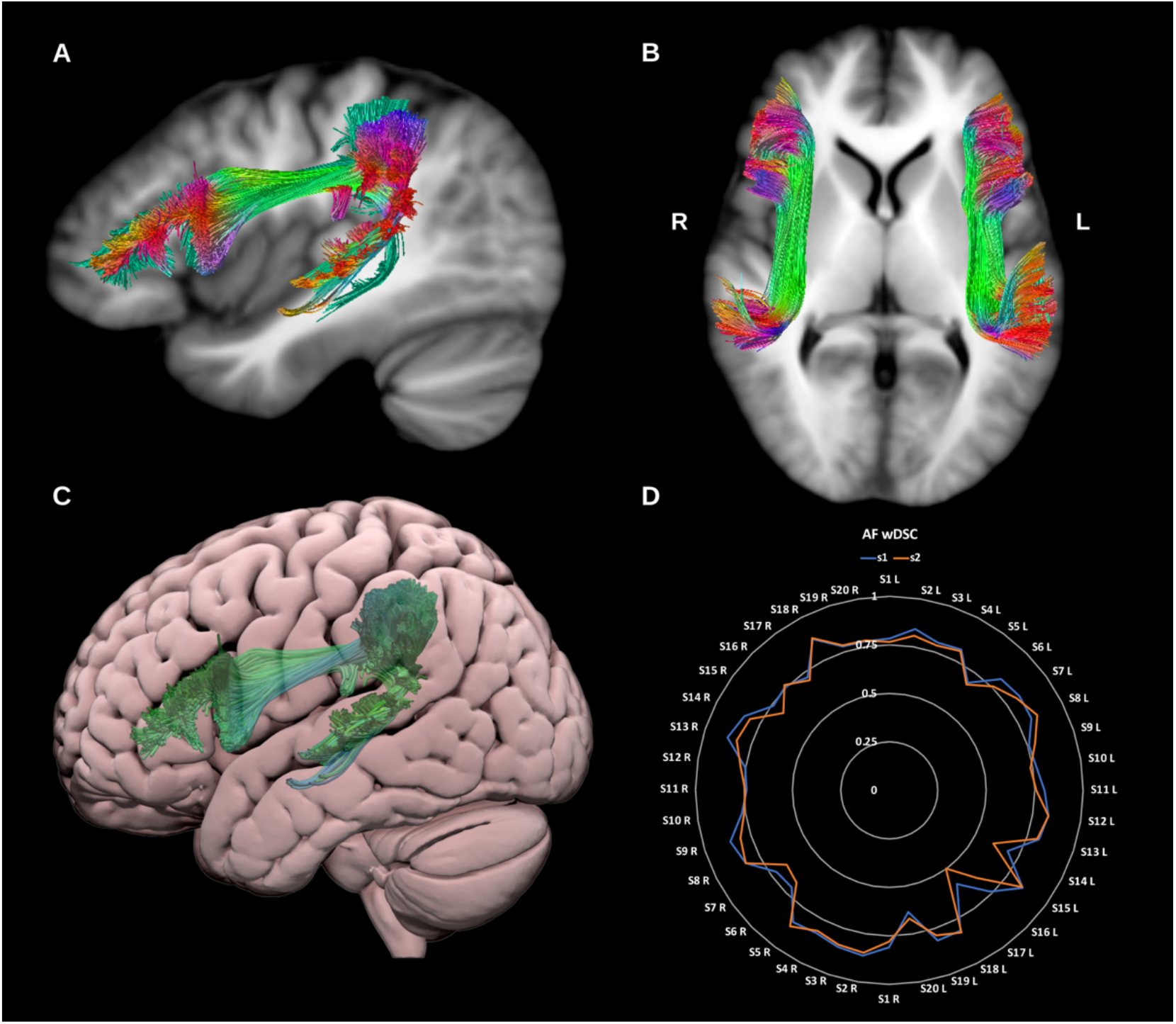
(A) and (B) show sagittal and axial slices of the T1-weighted group average image with the **arcuate fasciculi** overlaid in directional color coding. (C) shows a lateral projection of the semitransparent MNI pial surface with the left arcuate fasciculus in green. (D) shows a radar plot of the wDSC scores (vertical range) of both AFs using first session (blue) and second session (orange). Left (L) AF’s are on the first half of the circle and right (R) ones are on the second half. AF = arcuate fasciculus. MNI = Montreal Neurological Institute, wDSC = weighted dice similarity coefficient.

##### 3.1.1.2 Cingulum (CG)

The cingulum bundle (CG) is the principal WM tract of the cingulate gyrus and the limbic system and is involved in a diverse range of functions spanning emotional, behavioral and sensorimotor control, mnemonic processing, nociception and executive function. Broadly speaking, the cingulum is a bidirectional fiber system that encircles the corpus callosum lateral to the cingulate gyrus, extending from the frontal lobes to the WM of the ventral temporal lobe with numerous lateral projections joining and leaving the bundle along its path (Jones et al., 2013; Metzler-Baddeley et al., 2012; Pascalau et al., 2018; Wu et al., 2016b), see figure 3. The complexity of cingulum connections is inadequately characterized by dMRI, which tends to reconstruct either a single unilateral bundle extending from the subgenual cingulate to the temporal lobe, or two subdivisions encompassing the subgenual-dorsal, and dorsal-temporal segments. However, primate tracer studies suggest the CG could be subdivided into three or four regions based on differences in anatomical connectivity and neurotransmitter profile (Heilbronner and Haber, 2014). Here we reconstruct two sub-divisions in keeping with previous dMRI literature.

**Figure 3:**
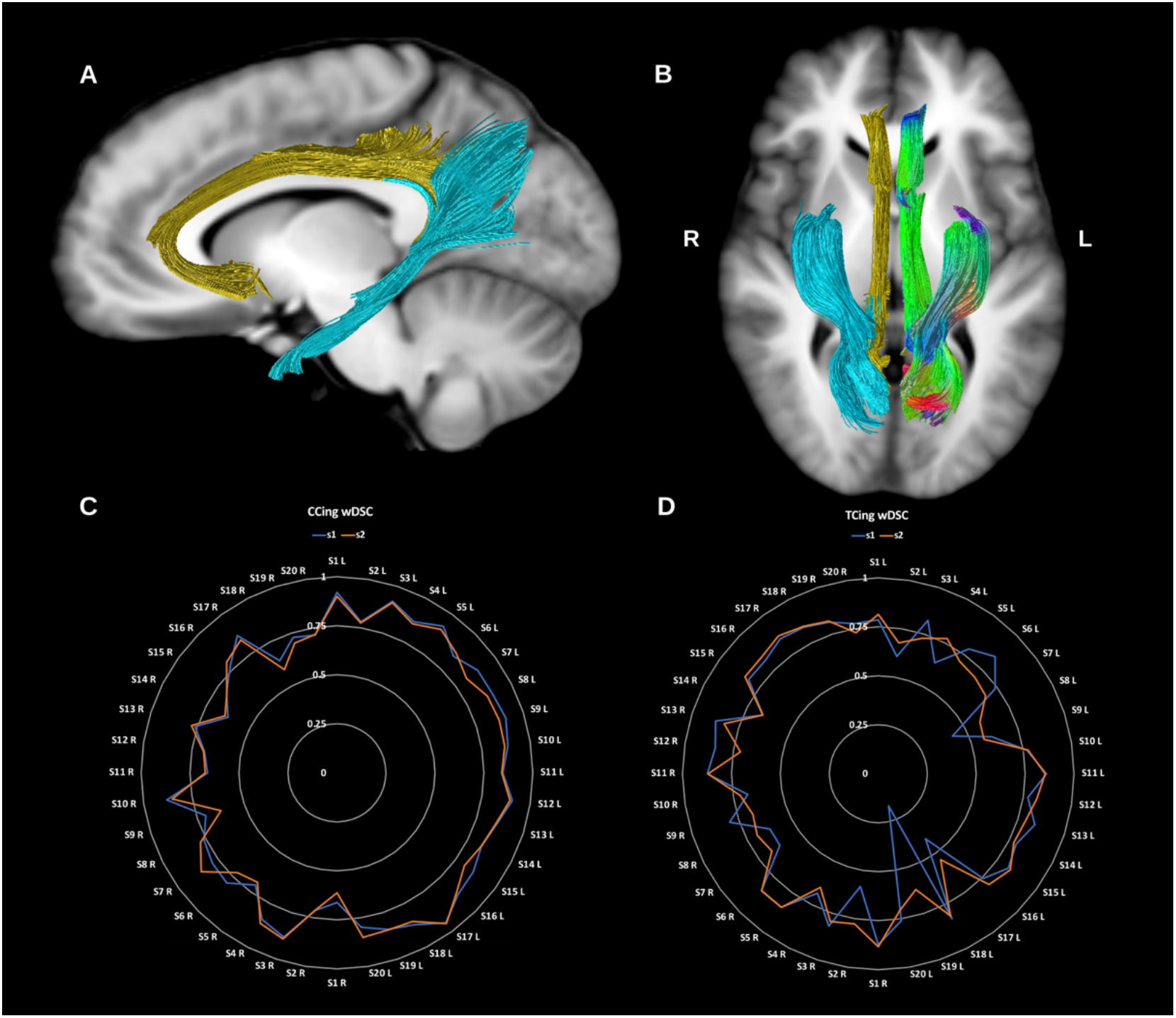
(A) and (B) show sagittal and axial slices of the T1-weighted group average image with the **cingulum bundles** overlaid in directional color coding for the left side and in gold for the cingulate portion (CCing) and light blue for the temporal portion (TCing). (C) and (D) show radar plots of the wDSC scores (vertical ranges) of both the CCings and TCings as reconstructed using first session (blue) and second session (orange). Left sided bundles (L) are on the first half of the circle and right ones (R) are on the second half. wDSC = weighted dice similarity coefficient

###### 3.1.1.2.1 Cingulate Cingulum

This represents the dorsal (cingulate) component of the cingulum and is reconstructed by tracking the streamlines between the rostral and caudal anterior cingulate cortex and the posterior cingulate, isthmic posterior cingulate cortex and the precuneus. It is immediately superior to the body of the corpus callosum. It has a low curvature C shape, tipped on its open end with anterior/ventral extension to the subgenual cortex and ends posteriorly behind the splenium of the corpus callosum. The cingulate cingulum is generated for all datasets on both sides.

###### 3.1.1.2.2 Temporal Cingulum

This represents the ventral component of the cingulum and is reconstructed by tracking the streamlines between the hippocampus and the isthmic posterior cingulate cortex and precuneus posteriorly. It is located in the medial temporal lobe, and has a slanted orientation, ascending from the anterior medial temporal lobe to the midline parietal region, behind the splenium of the corpus callosum. The temporal cingulum is generated for all datasets on both sides but was less reproducible than the cingulate cingulum.

##### 3.1.1.3 Fornix (FX)

The fornix is the major fiber pathway associated with the hippocampus, and comprises predominantly efferent fibers connecting the hippocampus with the prefrontal cortex, the anterior thalamic nuclei, the mammillary bodies, the ventral striatum, and the basal forebrain. Initially formed by the alveus and fimbria, WM of the bilateral hippocampus coalesce as the fornix crus and body before diverging again into the pre-commissural and post-commissural fornix columns which derive their name from their position relative to the anterior commissure. Recent research suggests fibers within the fornix are arranged topographically reflecting functional anterior-posterior gradient along the long axis of the hippocampus, with laterally located fibers arising from the anterior hippocampus and medially located fibers originating in the posterior hippocampus (Christiansen et al., 2017; Liacu et al., 2012; Pascalau et al., 2018; Strange et al., 2014), see figure 4. Here we reconstruct the fornix as a single lateralized bundle for each side. Reproducibility was poor and reconstruction failed in 15 datasets.

**Figure 4:**
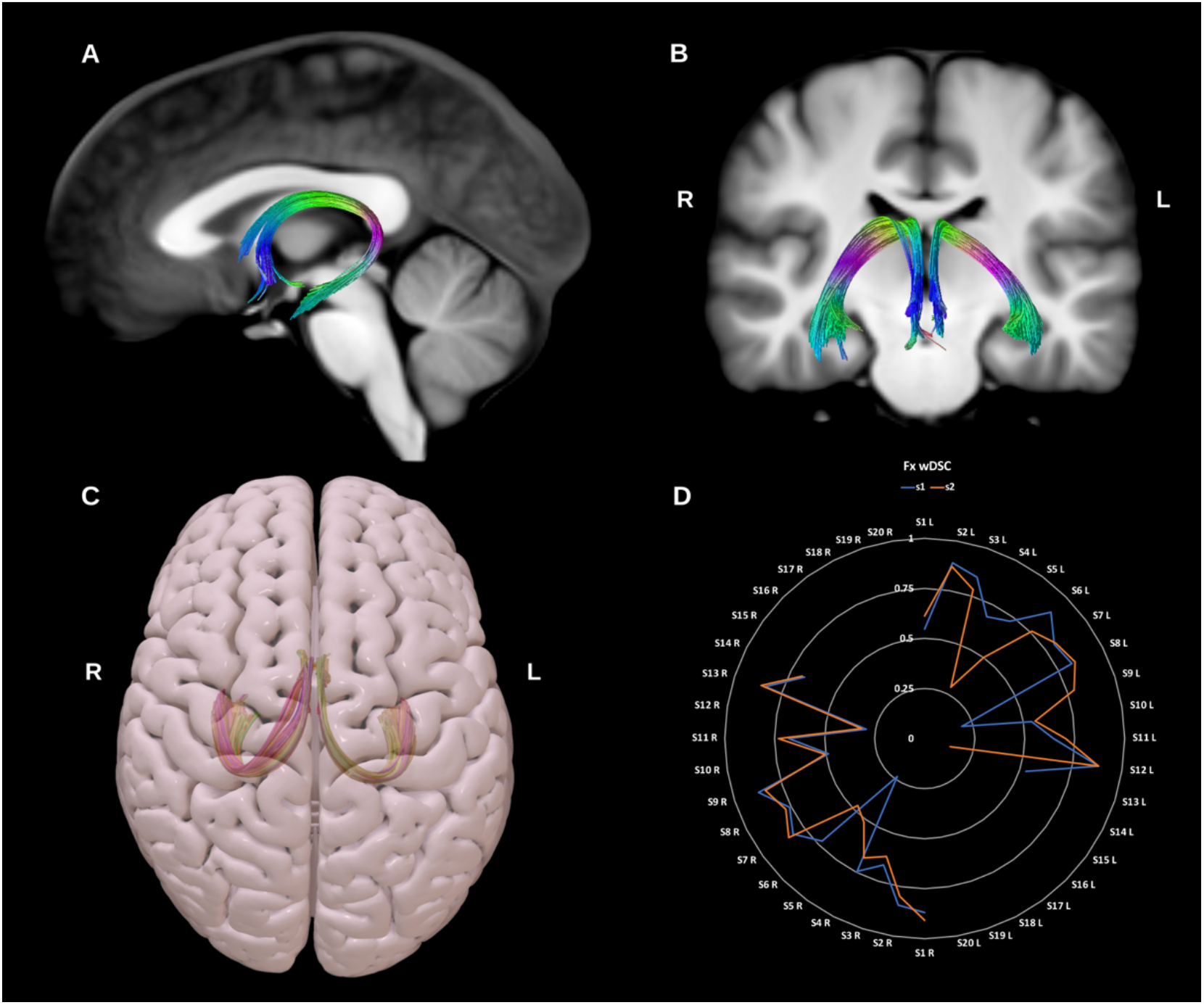
(A) and (B) show sagittal and coronal slices of the T1-weighted group average image with the **fornix (Fx)** overlaid in directional color coding. (C) shows a superior projection of the semitransparent MNI pial surface with both fornices shown in yellow and red. (D) shows a radar plot of the wDSC scores (vertical range) of both fornices as reconstructed using first session (blue) and second session (orange) data. A missing line indicates a missing bundle for that dataset. Left sided bundles (L) are on the first half of the circle and right ones (R) are on the second half. MNI = Montreal Neurological Institute, wDSC = weighted dice similarity coefficient.

##### 3.1.1.4 Frontal aslant tract (FAT)

The FAT is a recently described association bundle connecting the inferior and superior frontal lobe, and is involved in speech initiation, verbal fluency and executive function/inhibitory control. More specifically it connects the pars opercularis and pars triangularis in the inferior frontal gyrus (IFG) with the pre-supplementary motor area (pre-SMA), supplementary motor area (SMA), and the anterior cingulate cortex (Dick et al., 2019; La Corte et al., 2021; Pascual-Diaz et al., 2020), see figure 5. The FAT was successfully reconstructed for all datasets on both sides.

**Figure 5:**
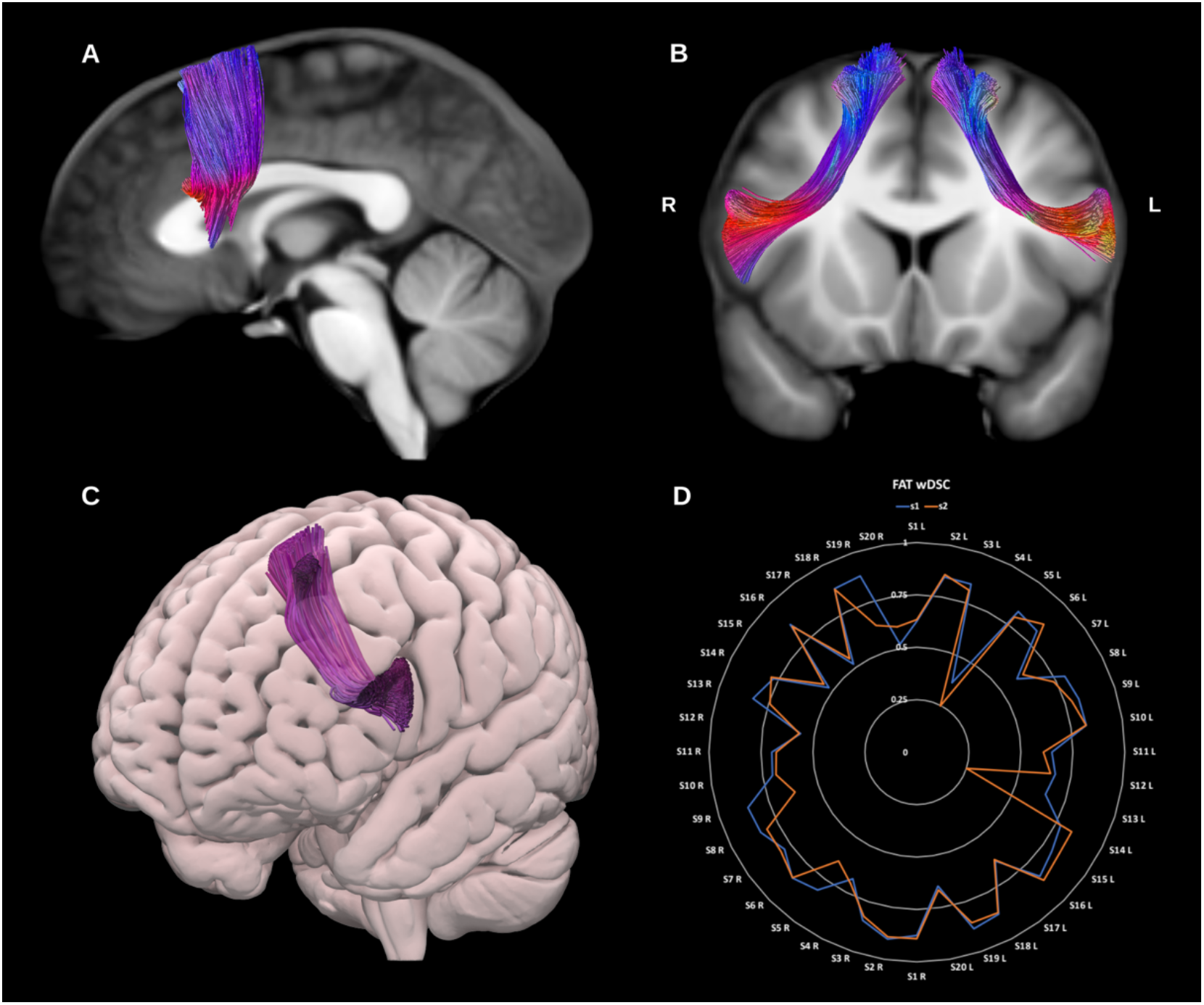
(A) and (B) show sagittal and coronal slices of the T1-weighted group average image with the **frontal aslant tracts (FAT)** overlaid in directional color coding. (C) shows an oblique anterior projection of the semitransparent MNI pial surface with left FAT in purple. (D) shows a radar plot of the wDSC scores (vertical range) of both FATs as reconstructed using first session (blue) and second session (orange) data. Left sided bundles (L) are on the first half of the circle and right ones (R) are on the second half. MNI = Montreal Neurological Institute, wDSC = weighted dice similarity coefficient.

##### 3.1.1.5 Inferior Fronto-occipital fasciculus (IFOF)

The IFOF is a large, long-range association fiber bundle connecting the occipital and temporal lobes to the frontal lobes, specifically the lingual, posterior fusiform, cuneus and polar occipital cortex, with the inferior frontal gyrus, medial fronto-orbital region and frontal pole, see figure 6. Notably, it narrows at the level of the extreme capsule. Though theoretically distinct from other temporal lobe association pathways, the IFOF runs in close proximity to the middle longitudinal fasciculus (MdLF), inferior longitudinal fasciculus (ILF) and uncinate fasciculus (UF), which may pose issues for some tracking algorithms (Forkel et al., 2014; Wu et al., 2016a). Considerable debate prevails regarding its exact functions however it is believed to serve the ventral visual and language streams along with the inferior longitudinal fasciculus (ILF) and uncinate fasciculus (UF) (Caverzasi et al., 2016). The right IFOF may be related to facial recognition functions and semantic visual stream processes (Herbet et al., 2018), while the left IFOF is related to semantic language functions (Almairac et al., 2015). The IFOF was generated for all datasets on both sides.

**Figure 6:**
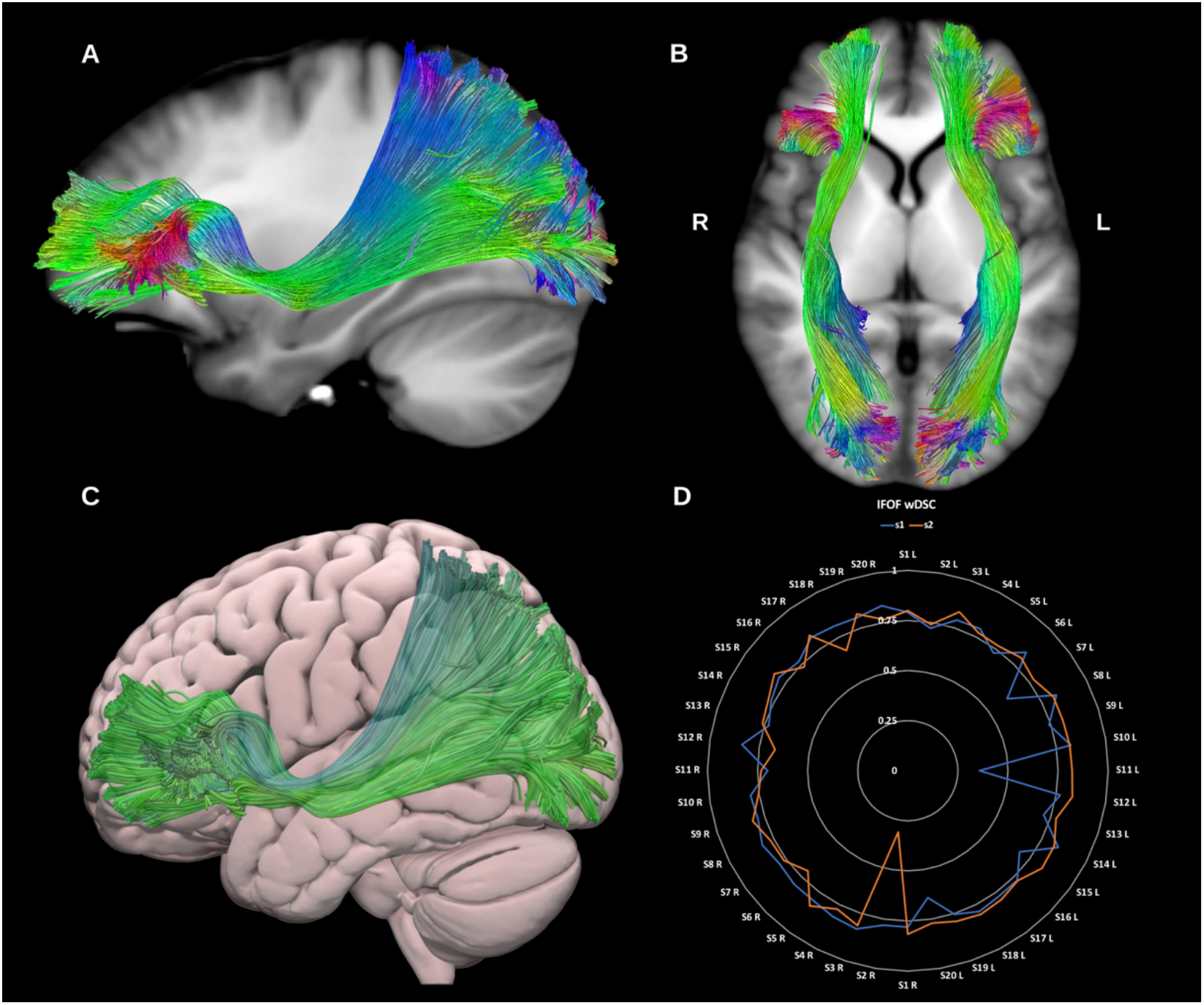
(A) and (B) show sagittal and axial slices of the T1-weighted group average image with the **inferior fronto-occipital fasciculi (IFOF)** overlaid in directional color coding. (C) shows a lateral projection of the semitransparent MNI pial surface with left IFOF in green. (D) shows a radar plot of the wDSC scores (vertical range) of both IFOFs as reconstructed using first session (blue) and second session (orange) data. Left sided bundles (L) are on the first half of the circle and right ones (R) are on the second half. MNI = Montreal Neurological Institute, wDSC = weighted dice similarity coefficient.

##### 3.1.1.6 Inferior longitudinal fasciculus (ILF)

The ILF is a large association tract connecting the occipital and temporal lobes, and may play an important functional role in visual memory and emotional processing. It lies in direct contact with or close proximity to several bundles, including the UF, IFOF, AF, optic radiations (ORs) and tapetal fibers of the corpus callosum (CC). Whilst commonly depicted as a single bundle, the ILF may have up to four morphological subdivisions which reflect its occipital termination points, and include lingual, cuneate, fusiform and dorso-lateral occipital subcomponents. However, given the lack of consensus about the existence and functional significance of these subdivisions, here we construct the ILF as a single bundle (Herbet et al., 2018; Latini et al., 2017; Panesar et al., 2018), see figure 7. The ILF was generated for all datasets on both sides.

**Figure 7:**
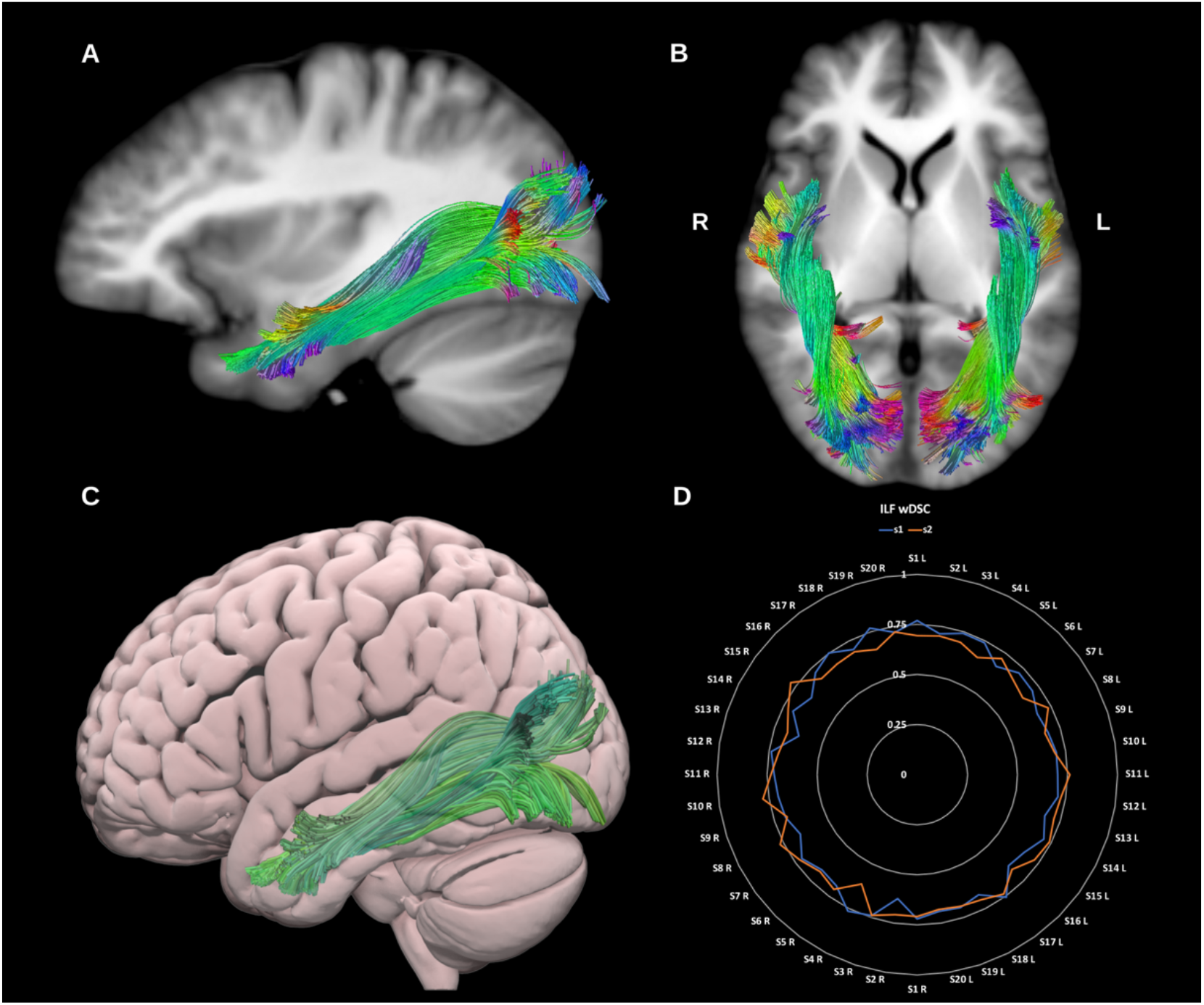
(A) and (B) show sagittal and axial slices of the T1-weighted group average image with the **inferior longitudinal fasciculi (ILF)** overlaid in directional color coding. (C) shows a lateral projection of the semitransparent MNI pial surface with the left ILF in green. (D) shows a radar plot of the wDSC scores (vertical range) of both ILFs as reconstructed using first session (blue) and second session (orange) data. Left sided bundles (L) are on the first half of the circle and right ones (R) are on the second half. MNI = Montreal Neurological Institute, wDSC = weighted dice similarity coefficient.

##### 3.1.1.7 Middle Longitudinal Fasciculus (MdLF)

The MdLF is a large association bundle which is hypothesized to play a role in language, visual and auditory processing (De Witt Hamer et al., 2010; Dick and Tremblay, 2012; Kalyvas et al., 2020). Broadly speaking it is thought to connect the temporal pole, superior temporal gyrus, angular gyrus, superior parietal lobule and precuneus forming a distinct association bundle that runs medial to the AF and lateral to the IFOF (Makris et al., 2017, 2009; Maldonado et al., 2013; Menjot de Champfleur et al., 2013; Seltzer and Pandya, 1984; Wang et al., 2013), see figure 7. However, there remains considerable debate about the anatomical description of the MdLF owing to its relatively recent discovery in humans, and the evolution in its connectivity profile as a result of differences in tracking algorithms. Indeed, an accurate reconstruction of the MdLF is hindered by its close proximity to the extreme capsule, ILF, IFOF and AF. The MdLF was generated for all datasets on both sides.

##### 3.1.1.8 Superior Longitudinal Fasciculus (SLF – I, II, and III)

The SLF is a parieto-occipital association fiber system, located in the dorso-lateral aspect of the cerebrum and generally believed to comprise three or four components (de Schotten et al., 2011; Wang et al., 2016). Broadly speaking it connects the frontal with the occipital, parietal and temporal lobes, see figure 9. The SLF is related to language (Madhavan et al., 2014) visuo-spatial (Hong et al., 2019), and meta-cognitive (Zheng et al., 2020) functions. When tracked as a whole the SLF is highly reproducible, being successfully generated in all sessions for all datasets on both sides.

**Figure 8:**
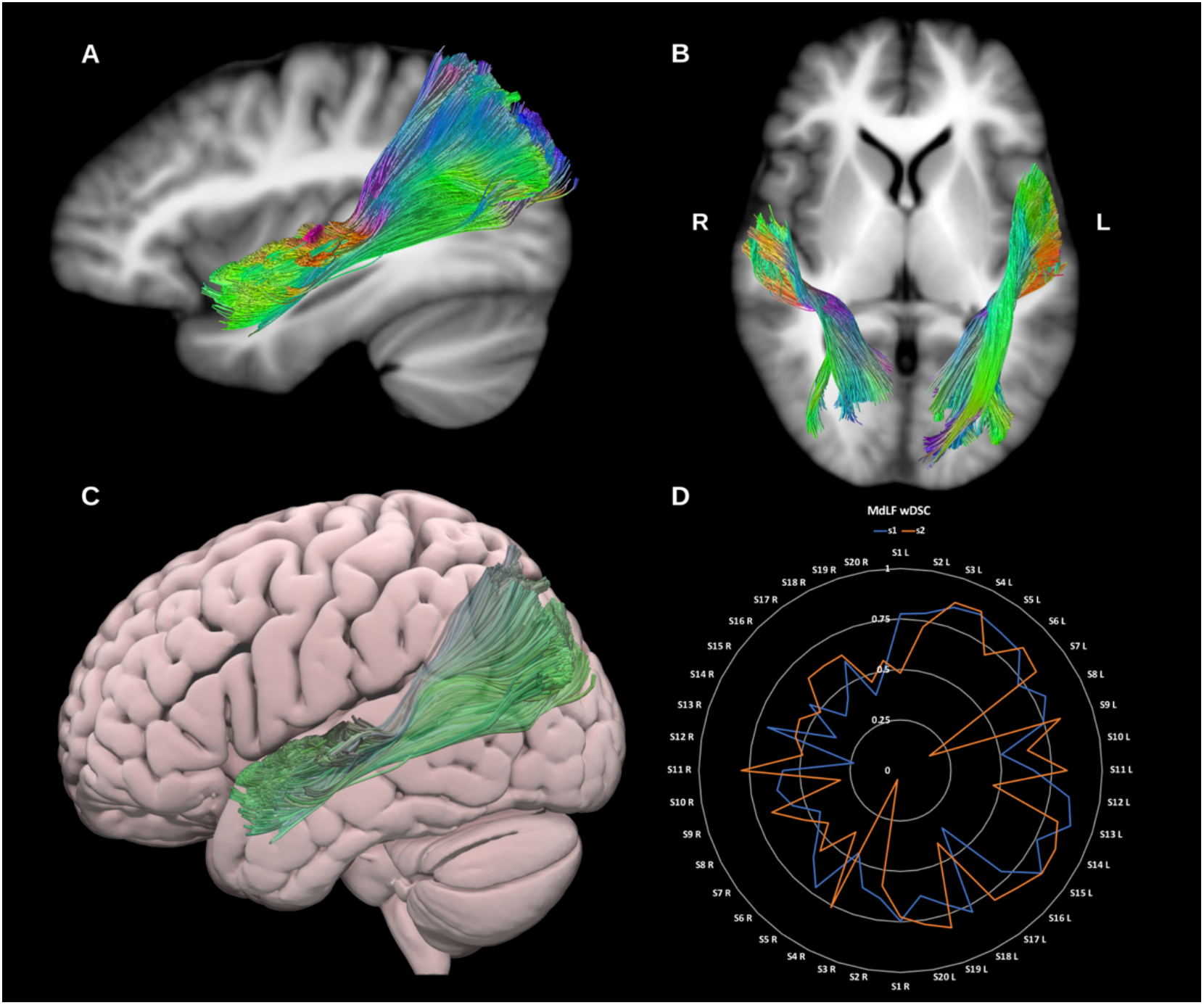
(A), and (B) show sagittal and axial slices of the T1-weighted group average image with the **Middle longitudinal fasciculi (MdLF)** overlaid in directional color coding. (C) shows a lateral projection of the semitransparent MNI pial surface with the left MdLF in green. (D) shows a radar plot of the wDSC scores (vertical range) of both MdLFs as reconstructed using first session (blue) and second session (orange) data. Left sided bundles (L) are on the first half of the circle and right ones (R) are on the second half. MNI = Montreal Neurological Institute, wDSC = weighted dice similarity coefficient.

**Figure 9:**
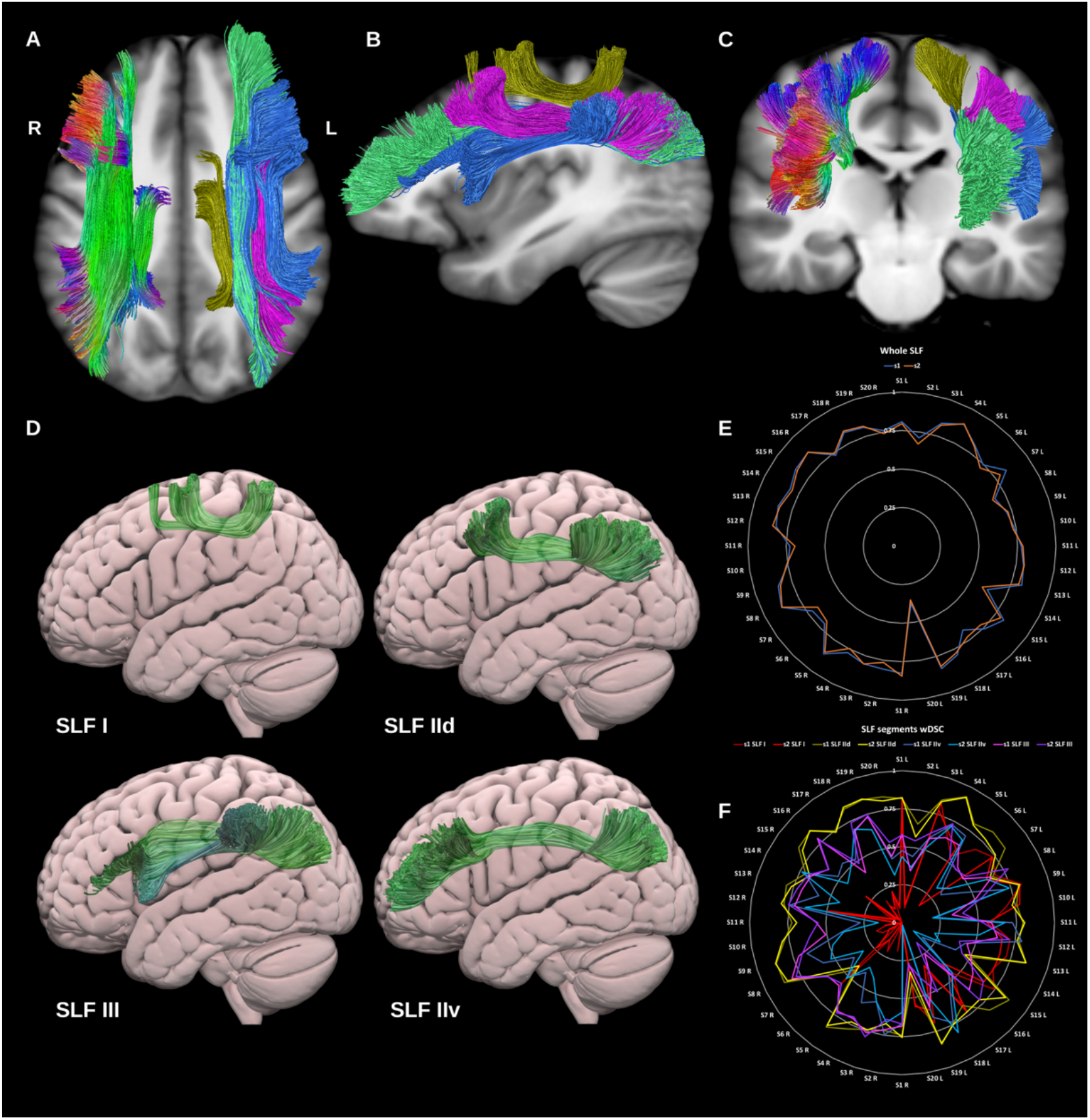
(A), (B), and (C) show sagittal and axial slices of the T1-weighted group average image with the **superior longitudinal fasciculi (SLF)** overlaid in directional color coding on the right side and show in gold (SLF-I), pink (SLF-IId), green (SLF-IIv) and blue (SLF-III). (D) shows a radar plot of the wDSC scores (vertical range) of both whole SLFs as reconstructed using first session (blue) and second session (orange) data. Left sided bundles (L) are on the first half of the circle and right ones (R) are on the second half. (E) shows lateral projections of the semitransparent MNI pial surface with the left SLF I, IId, IIv, and III in green. (F) shows a radar plot of the wDSC scores (vertical range) of SLF components for each side. MNI= Montreal Neurological Institute, wDSC= weighted dice similarity coefficient.

###### 3.1.1.8.1 SLF-I

The SLF-I connects the superior parietal lobule and precuneus with posterior superior frontal cortical areas. It is a rather short association bundle that is located above the level of the cingulate cortex (Wang et al., 2016). This was generated for all datasets on both sides but was the least reproducible of the SLF segments, most likely due to its small volume and low streamlines count.

###### 3.1.1.8.2 SLF-II

SLF-II is located more infero-lateral to SLF-I and originates in the anterior intraparietal sulcus and the angular gyrus, terminating in the posterior regions of the superior and middle frontal gyri. SLF-II can be subdivided into two distinct subcomponents (Barbeau et al., 2020), the first is the dorsal component (SLF-IId), which connects the supramarginal and inferior parietal cortices to the dorsal middle frontal gyrus. The second is the ventral component (SLF-IIv), which is longer, more ventral and connects the supramarginal and inferior parietal cortices to the rostral middle frontal gyrus. The SLF-IId and SLF-IIv were successfully reconstructed for all datasets on both sides.

###### 3.1.1.8.3 SLF-III

SLF-III connects the intraparietal sulcus and inferior parietal lobule to the inferior frontal gyrus (Barbeau et al., 2020). It is immediately superior and medial to the arcuate fasciculus partially overlapping with the horizontal fibers of the Arcuate fasciculus. The SLF-III was generated for all datasets on both sides.

##### 3.1.1.9 Uncinate Fasciculus (UF)

The UF is typically characterized as a hook shaped bidirectional association fiber bundle linking the ventral, medial and orbital frontal lobes and rostral temporal lobes. Whilst DTI typically reconstructs a short hook-shaped fasciculus (Kurki et al., 2013), higher order dMRI reconstructions have revealed a more extensive fiber system which expands into a fan shaped trajectory in the frontal lobe. Such higher order models (e.g., CSD) in combination with microdissection suggest that the UF may be further subdivided into 5 subcomponents (Hau et al., 2017). However, for the present reconstruction, we opted to reconstruct the UF as one bundle for practicality, see figure 10. This was generated for all datasets on both sides.

**Figure 10:**
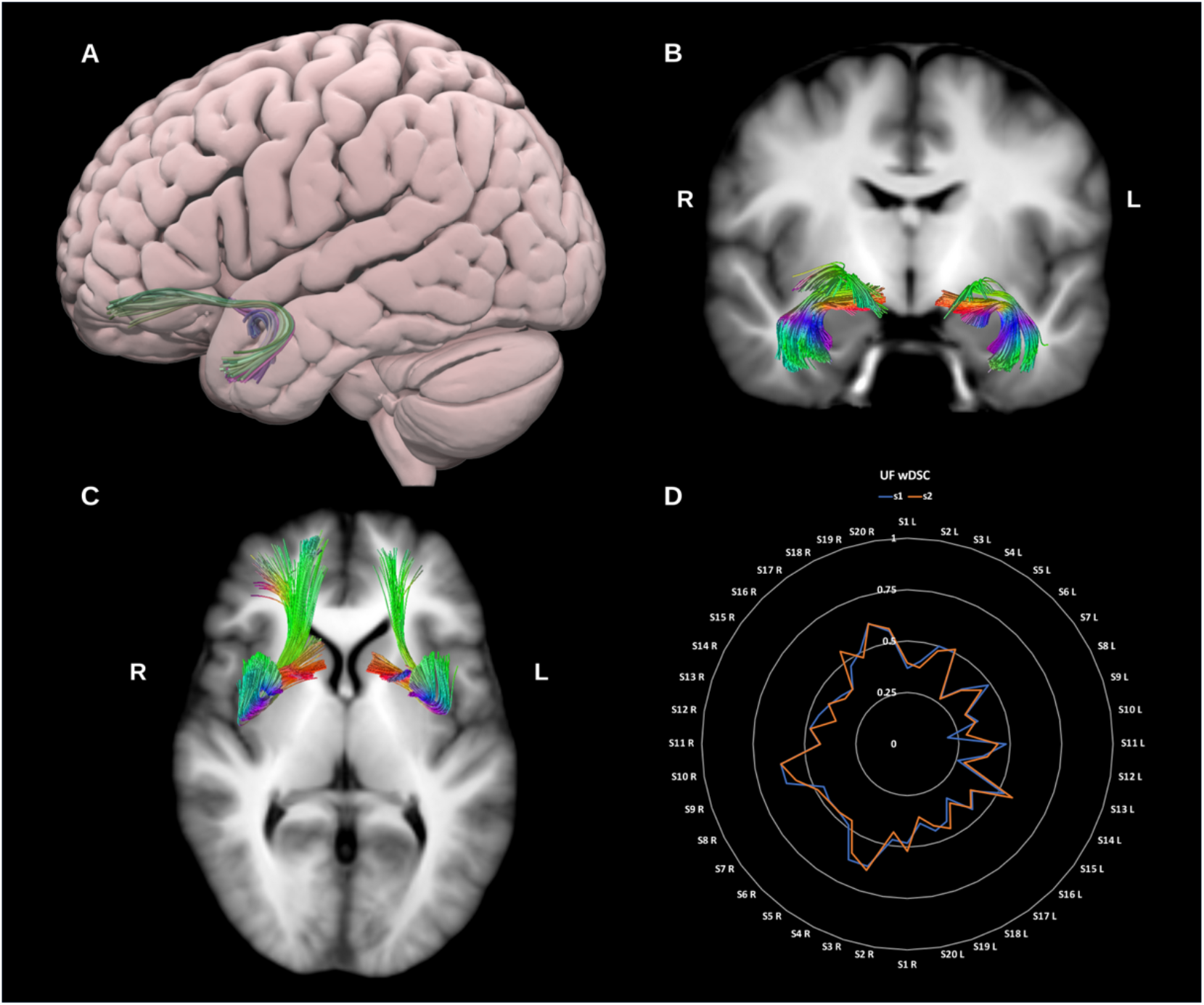
(A) shows a lateral projection of the semitransparent MNI pial surface with the left **uncinate fasciculus (UF)** in directional color coding, (B) and (C) show coronal and axial slices of the T1-weighted group average image with the UF overlaid in directional color coding. (D) shows a radar plot of the wDSC scores (vertical range) of both UFs as reconstructed using first session (blue) and second session (orange) data. Left sided bundles (L) are on the first half of the circle and right ones (R) are on the second half. MNI = Montreal Neurological Institute, wDSC = weighted dice similarity coefficient.

##### 3.1.1.10 Vertical Occipital Fasciculus (VOF)

The VOF is described as a short slanted vertical association bundle in the lateral aspect of the occipital lobe. It connects the superior part of the occipital lobe and adjacent cortex of the occipito-parietal sulcus to the inferior aspect of the occipital lobe and adjacent occipito-temporal cortical areas (Jitsuishi et al., 2020; Schurr et al., 2019; Yeatman et al., 2014), see figure 11. This was generated for all datasets on both sides.

**Figure 11:**
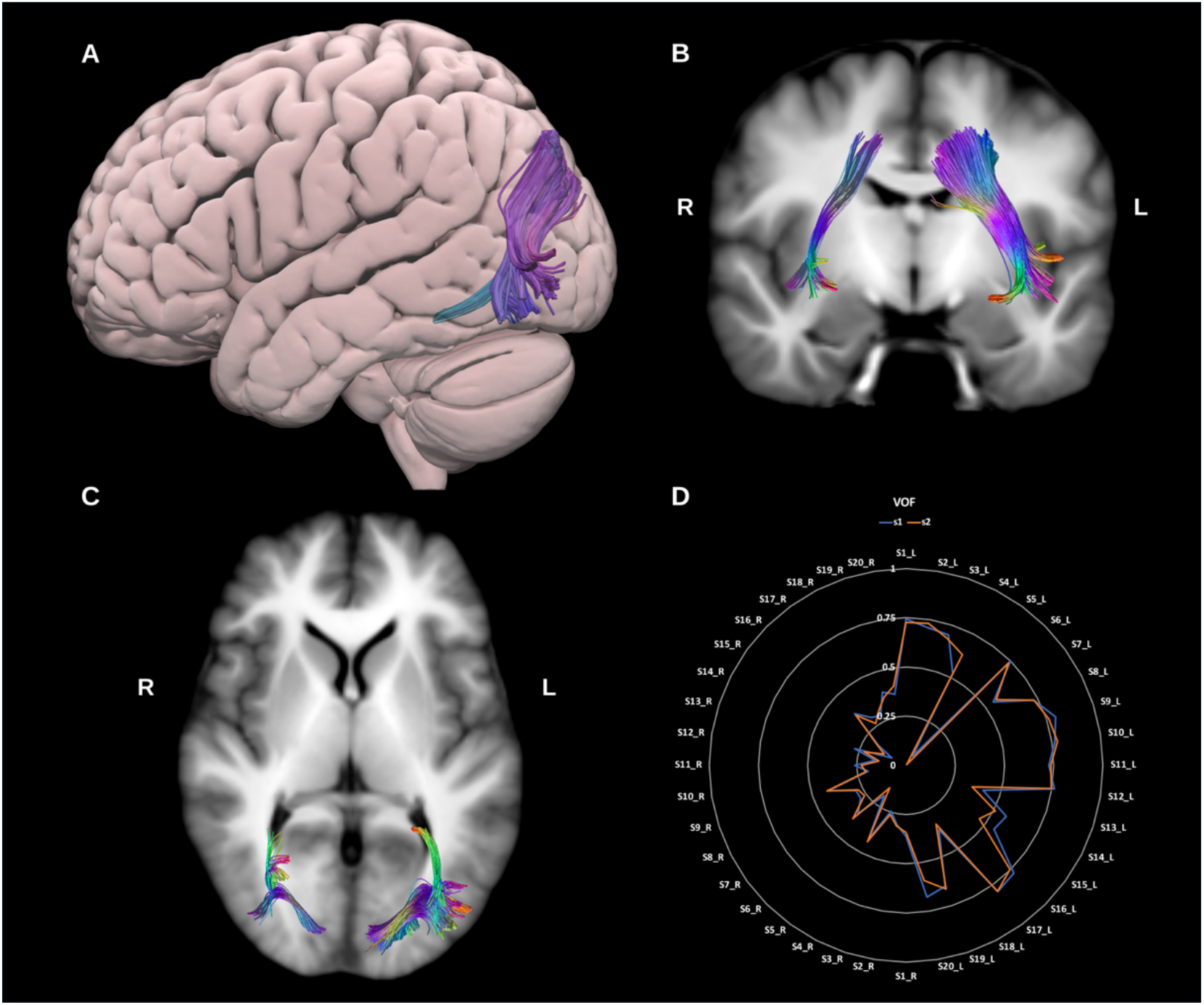
(A) shows a lateral projection of the semitransparent MNI pial surface with the left **vertical occipital fasciculus (VOF)** in directional color coding, (B) and (C) show coronal and axial slices of the T1-weighted group average image with the VOF overlaid in directional color coding. (D) shows a radar plot of the wDSC scores (vertical range) of both VOFs as reconstructed using first session (blue) and second session (orange) data. Left sided bundles (L) are on the first half of the circle and right ones (R) are on the second half. MNI = Montreal Neurological Institute, wDSC = weighted dice similarity coefficient.

#### 3.1.2 Commissural fiber bundles *(in alphabetical order)*

##### 3.1.2.1 Anterior Commissure (AC)

The anterior commissure crosses the midline anterior to the pre-commissural columns of the fornix, above the basal forebrain and below the medial and ventral portion of the anterior limb of the internal capsule. It has two main parts, the first, anterior division which includes the olfactory decussation and the second, largest division which connects the temporal lobes, occipital lobes, and predominantly the bilateral amygdalae (van Meer et al., 2016). During its course the AC intersects with the UF, ILF, sagittal stratum and optic radiations, which complicates tracking the true extent of the structure (Çavdar et al., 2021; Choi et al., 2011; Kikinis et al., 2015; Peltier et al., 2011; van Meer et al., 2016; Wilde et al., 2006), see figure 12. The AC was reconstructed successfully in both sessions for all but one subject.

**Figure 12:**
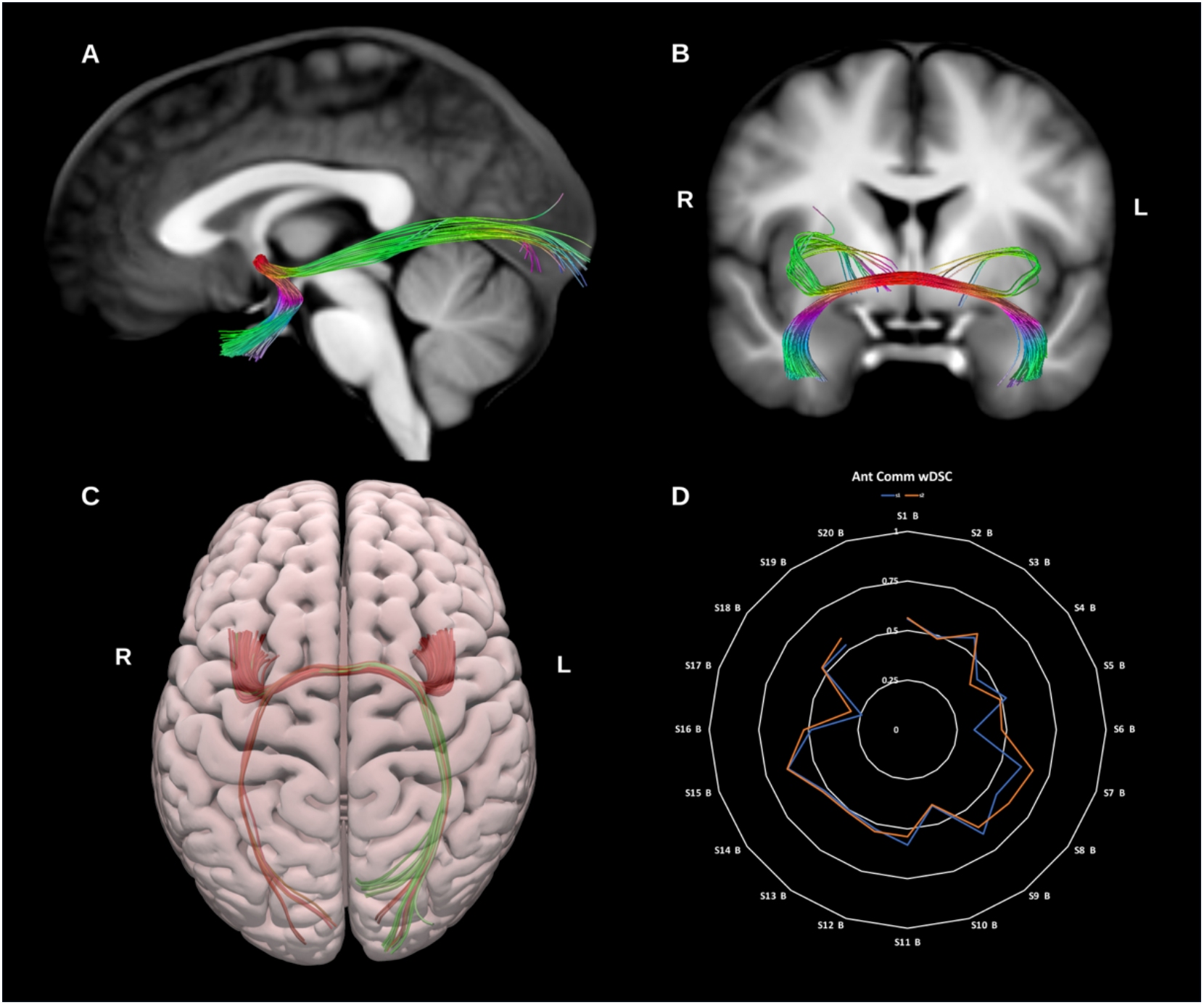
(A) and (B) show sagittal and coronal slices of the T1-weighted group average image with the **anterior commissure (Ant. Comm.)** overlaid in directional color coding, (C) shows a superior projection of the semitransparent MNI pial surface with the Ant. Comm. in directional color coding, and (D) shows a radar plot of the wDSC scores (vertical range) of the Ant. Comm. As reconstructed using first session (blue) and second session (orange) data. MNI = Montreal Neurological

##### 3.1.2.2 Corpus Callosum (CC)

The corpus callosum is the largest fiber bundle in the human brain and connects the right and left cerebral hemispheres, and hence its primary function is interhemispheric information transfer and integration. Traditionally the CC is divided into several subcomponents: the rostrum, genu, body, isthmus, splenium and tapetum. It is common in DTI based literature to find three subdivisions: the genu, which forms the forceps minor and connects left and right prefrontal and anterior cingulate cortices; the callosal body, and the splenium, which forms the forceps major and connects left and right posterior parietal, medial occipital and medial temporal cortices. Improvements in fiber-tracking algorithms and integration with functional data have led to alternative parcellation strategies (Catani, 2006; Fabri et al., 2014; Nazem-Zadeh et al., 2012; Ohoshi et al., 2019, 2019; Phillips et al., 2013). Here we have reconstructed the CC into 7 segments comprising the prefrontal, premotor, motor, sensory, parietal, occipital and temporal callosal fibers, see figures 13 and 14.

**Figure 13:**
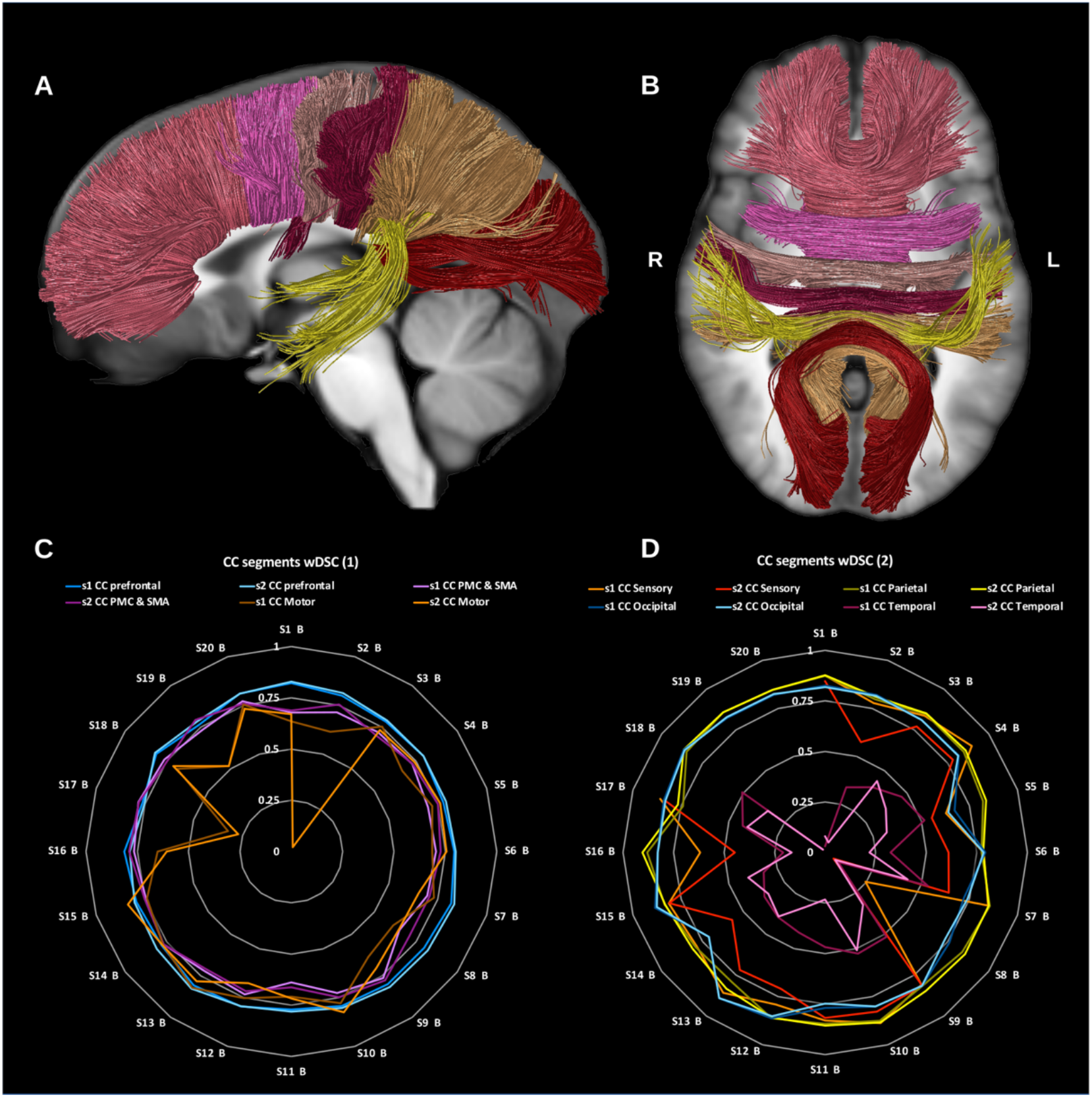
(A) and (B) show sagittal and axial slices of the T1-weighted group average image with the all segments of **corpus callosum (CC)**. Radar plots of wDSC (vertical ranges) resulting from comparison to HCP-template bundles, (C) Prefrontal CC (blue and light blue), premotor and supplementary motor CC (pink and fuschia), motor CC (dark and light orange), (D) sensory CC (orange and red), parietal CC (gold and yellow), occipital CC (blue and light blue), and temporal CC (fuschia and pink). PMC = premotor cortex, SMA = supplementary motor cortex.

**Figure 14:**
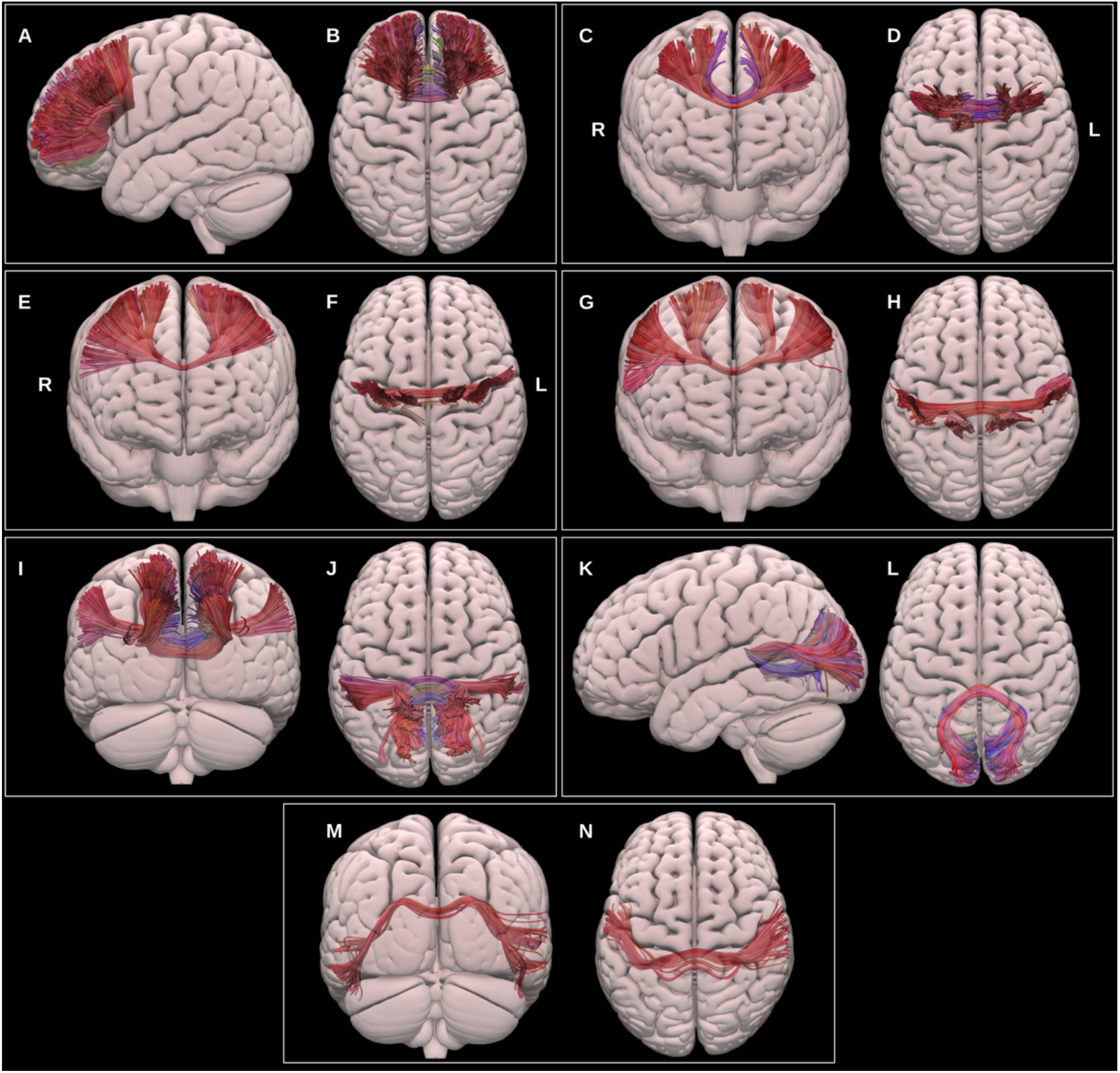
Multiple projections of the semitransparent MNI pial surface, with lateral and superior views showing the prefrontal CC (A & B), anterior and superior views showing the PMC and SMA CC (C & D), anterior and superior views showing the motor CC (E & F), anterior and superior views showing the sensory CC (G & H), posterior and superior views showing the parietal CC (I & J), lateral and superior views showing the occipital CC, and posterior and superior views showing the temporal CC (M & N).

###### 3.1.2.2.1 Prefrontal corpus callosum (CC PreF)

This was reconstructed for all datasets using the bilateral prefrontal cortices and the genu and anterior third of the body of the corpus callosum.

###### 3.1.2.2.2 Premotor and supplementary motor corpus callosum (CC PMC and SMA)

This was reconstructed for all datasets using the caudal middle frontal gyri, the supplementary motor areas bilaterally and the central body of the corpus callosum.

###### 3.1.2.2.3 Motor corpus callosum (CC motor)

This segment connects the primary motor cortices (precentral gyri) of both hemispheres via the posterior third of the body of the corpus callosum. The motor CC was generated for all datasets.

###### 3.1.2.2.4 Sensory corpus callosum (CC sensory)

This was reconstructed using the primary sensory cortices (postcentral gyri) on both sides via the posterior third of the body and splenium of the corpus callosum in the midline. The sensory CC was reconstructed for all except 4 datasets.

###### 3.1.2.2.5 Parietal corpus callosum (CC parietal)

This was reconstructed for all data using the whole parietal lobes bilaterally and the splenium of the corpus callosum.

###### 3.1.2.2.6 Occipital corpus callosum (CC occipital)

This was reconstructed for all data using the occipital lobes on both sides and the splenium of the corpus callosum.

###### 3.1.2.2.7 Temporal corpus callosum (CC temporal)

This was reconstructed using the lateral aspect of the temporal lobes bilaterally and the splenium of the corpus callosum in the midline. The temporal CC reconstruction failed in 1 dataset.

#### 3.1.3 Projection fiber bundles *(in alphabetical order)*

##### 3.1.3.1 Medial Lemniscus (ML)

The primary dorsal ascending tracts originate in the dorsal columns of the spinal cord, ascend through the brain stem medulla as the medial lemniscus to the thalamic ventral-posterior medial (VPM) and lateral (VPL) nuclei and via the internal capsule to the primary somatosensory cortex in the postcentral gyrus (Jang and Seo, 2015; Peng et al., 2019a), see figure 15. The ML was reconstructed using the UKBB derived posterior brainstem VOIs, the VPM and VPL thalamic nuclei, and primary sensory cortices. This was generated in all datasets on both sides.

**Figure 15:**
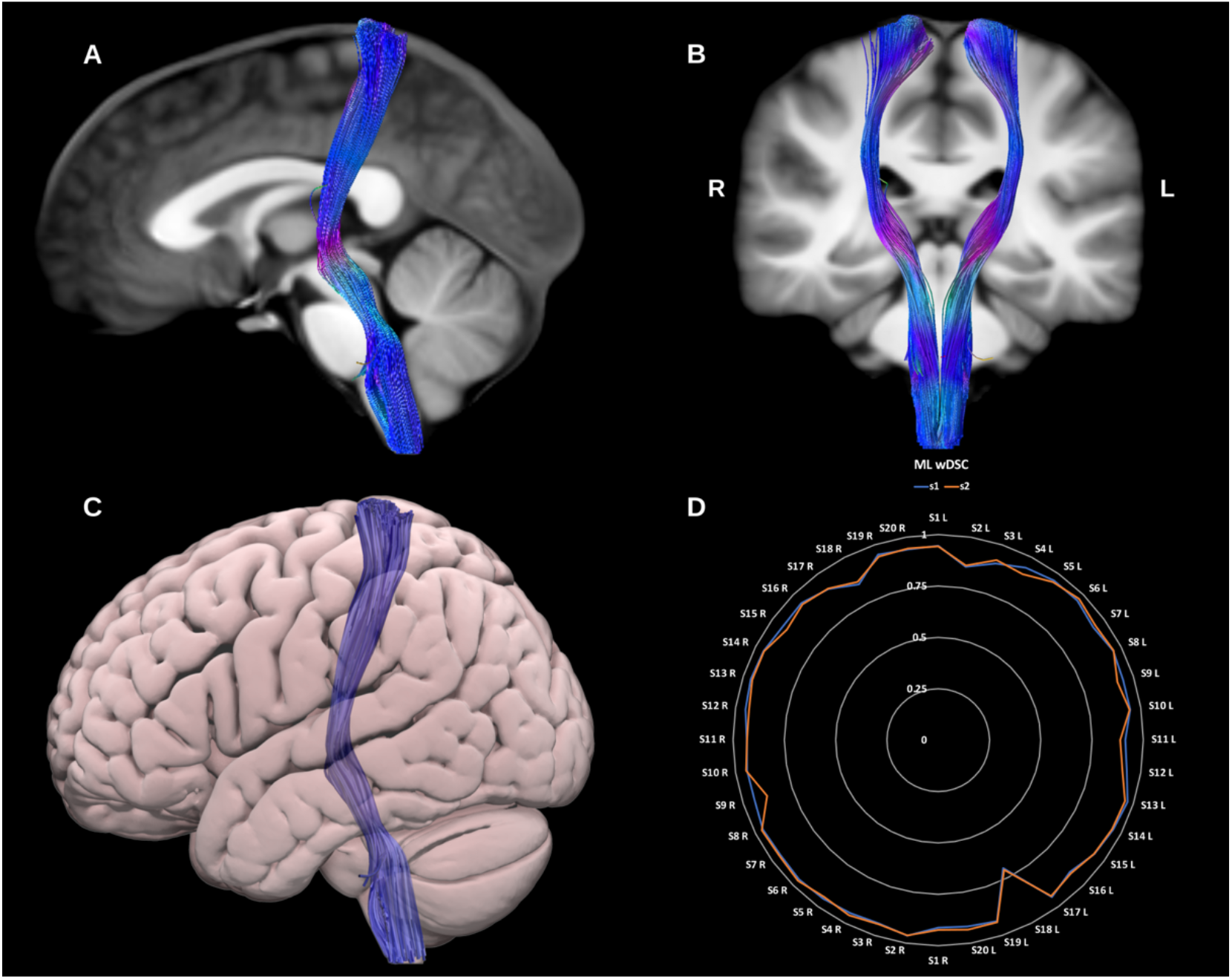
(A) and (B) show sagittal and coronal slices of the T1-weighted group average image with the **medial lemniscus (ML)** overlaid in directional color coding, (C) shows a lateral projection of the semitransparent MNI pial surface with the ML blue, and (D) shows a radar plot of the wDSC scores (vertical range) of the ML as reconstructed using first session (blue) and second session (orange) data. Left sided bundles (L) are on the first half of the circle and right ones (R) are on the second half. MNI = Montreal Neurological Institute, wDSC = weighted dice similarity coefficient.

##### 3.1.3.2 Optic Radiations (OR)

The ORs connect the lateral geniculate nucleus of the thalamus to the primary visual cortex within the occipital lobe. They are critical for visual processing and present important challenges during surgical procedures involving the temporal lobe. Notably, Meyer’s loop presents a particular challenge for tractography (Mehra and Moshirfar, 2021). This portion of the OR, which exhibits significant inter-individual variability, is characterized by a sharp anterior projection in the anterior temporal lobe that then bends posteriorly to join the sagittal stratum. Moreover, within the anterior loop, OR fibers may intersect with fibers of the anterior commissure, ILF and tapetum, increasing the likelihood of spurious streamlines (Chamberland et al., 2018; Goga and Türe, 2015; Lim et al., 2015; Martínez-Heras et al., 2015), see figure 16. We generated two representations of the optic radiations, first using a classical anatomical definition with only the pericalcarine cortex as a cortical inclusion VOI, and second using the entire occipital lobe as a cortical inclusion VOI. Both versions of the ORs were successfully reconstructed for all datasets on both sides.

**Figure 16:**
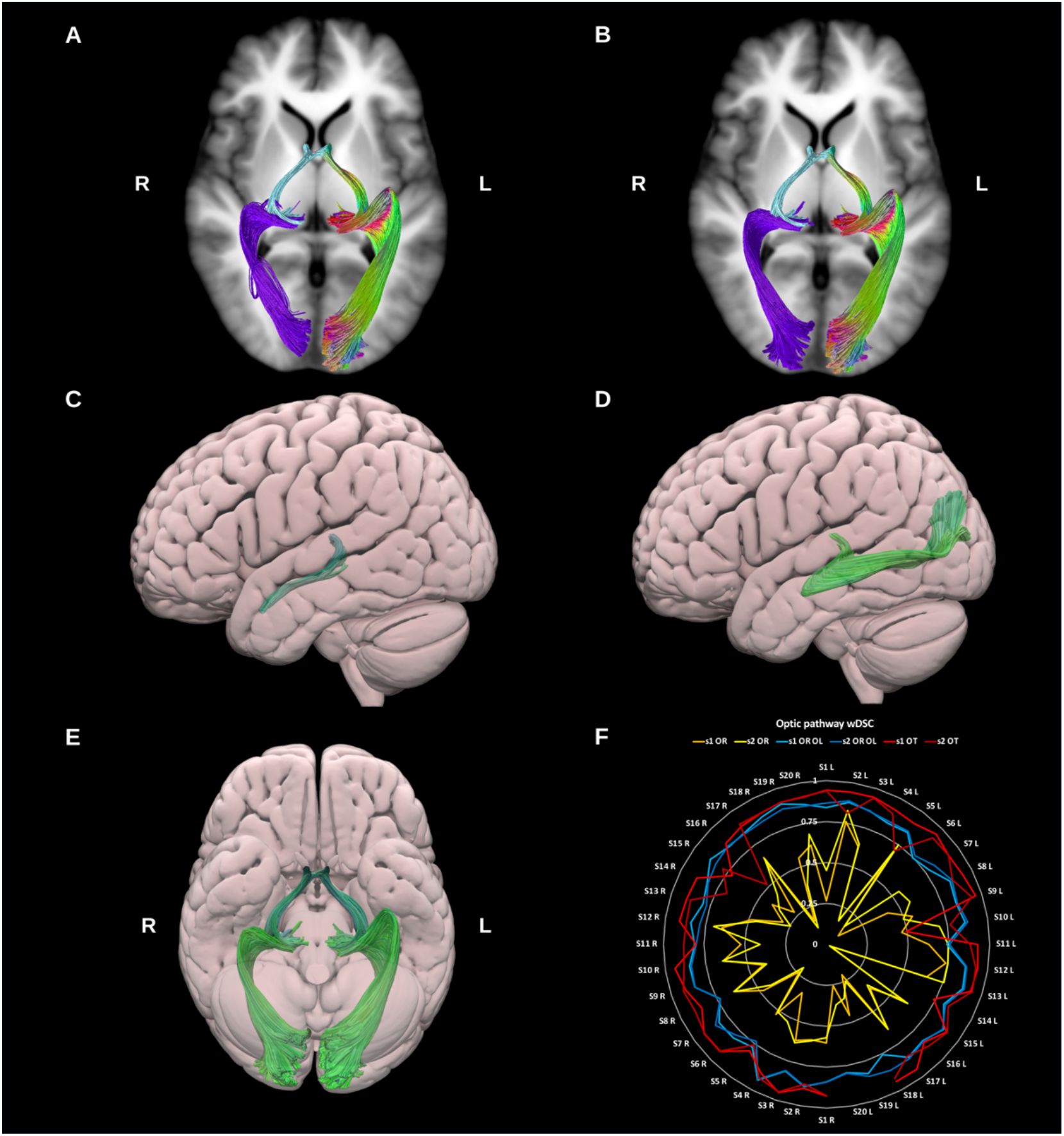
(A) and (B) show axial slices of the T1-weighted group average image with the overlaid optic tracts (light blue) and (A) classic **optic radiations (ORs)** (purple), and (B) whole occipital lobe optic radiations (OR OL) (purple) on the right side and directional color coding on the left side. (C) & (D) show lateral projections of the semitransparent MNI pial surface with the **optic tract (OT)** (C) and optic radiation (D). E shows an inferior projection of the surface view for the entire optic pathway on both sides. (F) shows a radar plot of the wDSC scores (vertical range) of the optic tracts and both versions of the optic radiations. Left sided bundles (L) are on the first half of the circle and right ones (R) are on the second half. MNI = Montreal Neurological Institute, wDSC = weighted dice similarity coefficient.

##### 3.1.3.3 Optic Tract (OT)

The OT is formed of decussating axons from the contra-lateral optic nerve as well as non-decussating fibers from the ipsilateral optic nerve. It extends from the optic chiasm to the lateral geniculate nucleus of the thalamus (Hofer et al., 2010; Mehra and Moshirfar, 2021; Wu et al., 2012), see figure 16. The OTs reconstruction failed in 3 datasets on the left side only.

##### 3.1.3.4 Pyramidal Tract (PyT)

The pyramidal tract fibers carry motor impulses from the cerebral cortex to the spinal cord through the brainstem. Here we reconstruct the Pyramidal tract (PyT) as a whole including the primary motor and sensory cortices along with the premotor, supplementary motor and parietal proprioceptive cortices superiorly and the whole brain stem inferiorly. We also provide dissections of this bundle based on specific subsystems of the sensory-motor network (Chenot et al., 2019; Jeong et al., 2013; Peng et al., 2019b; Zhang et al., 2010), see figure 17. The whole PyT was successfully reconstructed for all datasets on both sides, all its subcomponents were reconstructed successfully except for the premotor component, which failed in 6 datasets and was found to be the least reproducible.

**Figure 17:**
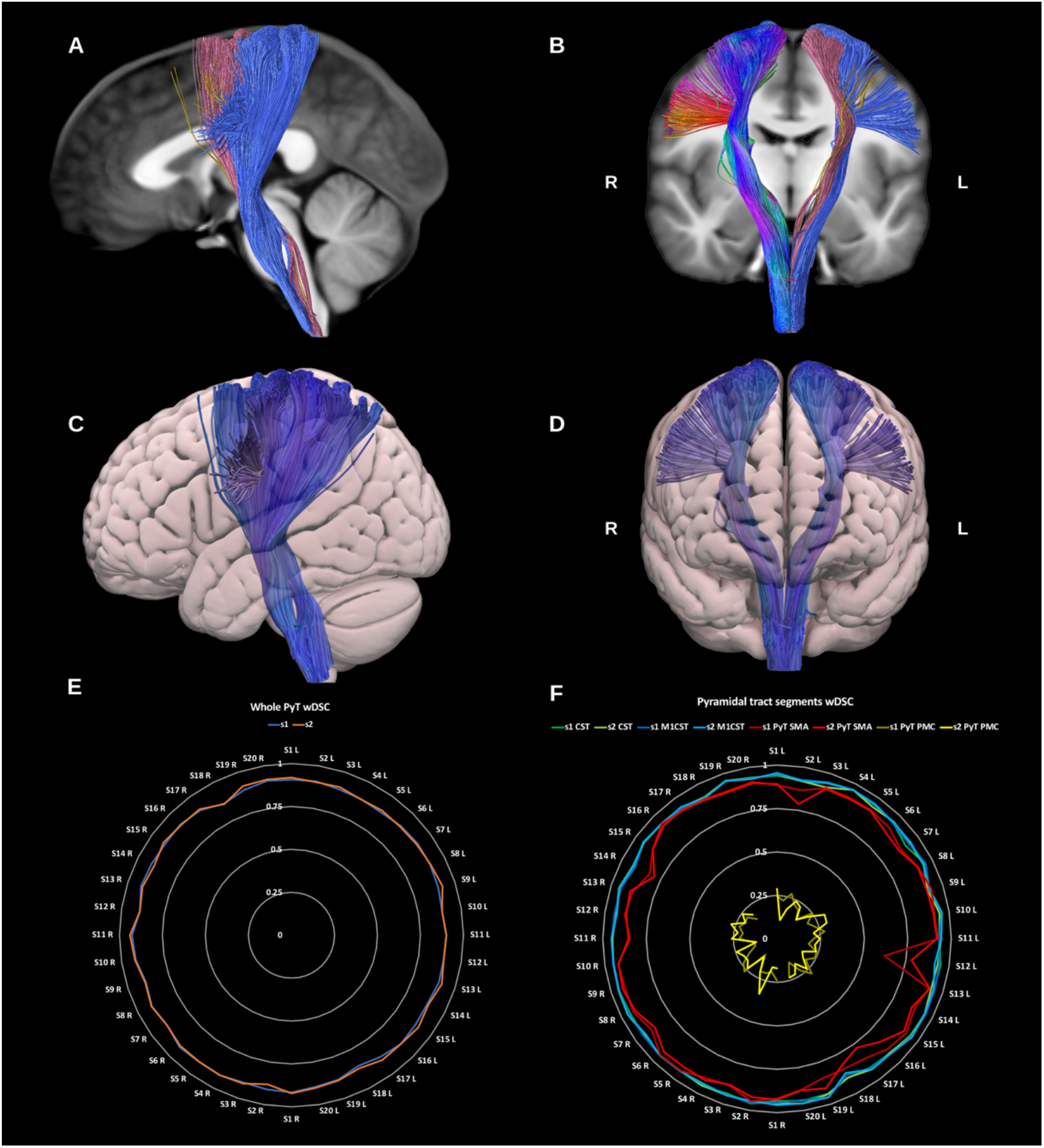
(A) and (B) show sagittal and coronal slices of the T1-weighted group average image with the **whole pyramidal tract (PyT_all)** overlaid in directional color coding on the right side and its different components on the left side in solid colors, premotor pyramidal tract (PyT_PMC) is shown in yellow, the supplementary motor pyramidal tract (PyT_SMA) in pink and the corticospinal tract (CST) in blue, (C) and (D) show lateral and anterior projections of the semitransparent MNI pial surface with the whole PyT on both sides shown in blue. Radar plots of wDSC (vertical ranges) resulting from comparison to HCP-template bundles are shown in (E) for the whole pyT, and in (F) for the different pyramidal tract segments. Left sided bundles (L) are on the first half of the circle and right ones (R) are on the second half. MNI = Montreal Neurological Institute, wDSC = weighted dice similarity coefficient, PyT = pyramidal tract, CST = corticospinal tract, M1 CST = motor only CST, SMA = supplementary motor area, PMC = premotor cortex, s = session, S = subject, L = left,

###### 3.1.3.4.1 Primary sensory-motor pyramidal Tract (Corticospinal Tract) (CST)

The CST descends predominantly from the primary motor areas of the precentral gyrus and somatosensory areas of the postcentral gyrus through the corona radiata, posterior half of the posterior limb of the internal capsule, and cerebral peduncles to the rostral brainstem where it forms the medullary pyramids before crossing the pyramidal decussation on its further descent through the spinal cord. We provide two reconstructions of this bundle, one using the primary motor and sensory cortices (CST) and another using only the primary motor cortices, excluding the postcentral gyri (M1_CST).

###### 3.1.3.4.2 Premotor pyramidal Tract (PyT PMC)

This represents the pyramidal tract fibers originating in the dorsal premotor cortex (caudal middle frontal gyrus) and descending to the brainstem via the anterior half of the posterior limb of the internal capsule. This bundle had the lowest number of streamlines and was the least reproducible of the pyramidal tract components.

###### 3.1.3.4.3 Supplementary motor pyramidal Tract (PyT SMA)

This represents the pyramidal tract fibers originating in the supplementary motor area and descending to the brainstem via the centrum semiovale and the anterior half of the posterior limb of the internal capsule.

##### 3.1.3.5 Thalamic radiations

The thalamic radiations are a group of 4-5 thalamo-cortical projection bundles comprising both afferent and efferent fibers. Collectively, the fibers of the thalamic radiations can be described as fanning out towards the cortex and banding up as they approach the thalami. As they run through the internal capsule and corona radiata they intersect with several other projection bundles, which in addition to the complex nature of thalamic microanatomy, can complicate reconstruction (Behrens et al., 2003; kakou et al., 2017; Lyness et al., 2014; Tsao et al., 2015; Younes et al., 2019), see figure 18. The posterior thalamic radiation is represented by the extended version of the optic radiation (OR OL) including the entire occipital cortex. All thalamic radiations were reconstructed using the whole thalamus as an inclusion VOI and excluding non-contributing thalamic nuclei, e.g., the VPM and VPL nuclei were excluded for the anterior thalamic radiation.

**Figure 18:**
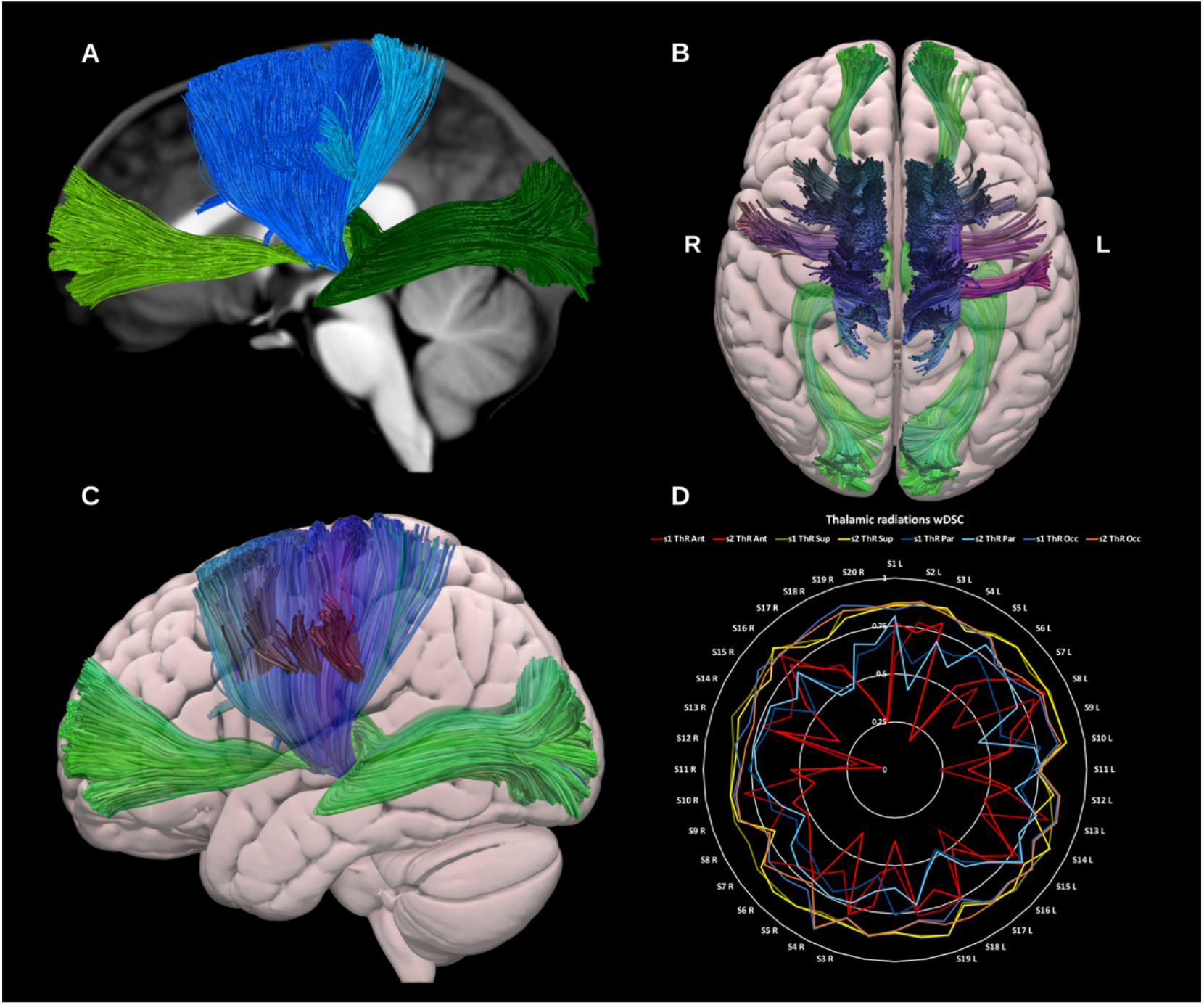
(A) shows a sagittal view of the T1-weighted group average image with the **thalamic radiations** overlaid in different colors. The anterior thalamic radiation is shown in light green, the superior thalamic radiation in blue, the parietal thalamic radiation in turquoise, and the posterior thalamic radiation in dark green. (B) and (C) show superior and lateral projections of the semitransparent MNI pial surface with the anterior and posterior thalamic radiations in green, and the superior and sensory thalamic radiations in blue. (D) shows a radar plot of the wDSC scores (vertical range) of the different thalamic radiations. Left sided bundles (L) are on the first half of the circle and right ones (R) are on the second half. MNI = Montreal Neurological Institute, wDSC = weighted dice similarity coefficient, ThR Ant = anterior thalamic radiation, ThR Sup = superior thalamic radiation, ThR Par = parietal thalamic radiation, ThR Occ = occipital thalamic radiation = Optic radiation to whole occipital lobe.

###### 3.1.3.5.1 Anterior thalamic radiation (ATR)

The ATR connects the dorso-medial (DM), dorso-lateral (DL) and anterior thalamic (ATN) nuclei to the prefrontal cortex and is thought to be involved in executive functions and complex planning (Niida et al., 2018). The ATR was reconstructed using the thalamus excluding all nuclei but the DM, DL, and ATN, and the ipsilateral medial and rostral prefrontal cortices. The ATR was reconstructed for all datasets on both sides.

###### 3.1.3.5.2 Superior thalamic radiation (STR)

The STR connects the ventral thalamic nuclei to the motor and sensory cortices via the superior thalamic peduncle, posterior limb of internal capsule, and the corona radiata, conducting cerebellar and basal ganglia input to the motor cortex (Bosch-Bouju et al., 2013). The STR was reconstructed using the thalamus, excluding the dorso-medial, dorso-lateral and pulvinar nuclei, and the ipsilateral primary motor, supplementary motor and dorsal premotor cortices (Younes et al., 2019). The STR was successfully reconstructed for all datasets on both sides.

###### 3.1.3.5.3 Parietal thalamic radiation (PaTR)

This was reconstructed using the entire parietal lobe as a cortical include and the thalamus excluding all except the VPM, and VPL nuclei. This resulted in a bundle of streamlines connecting the thalamic VPM and VPL predominantly to the primary sensory cortex. The PaTR was successfully reconstructed in all datasets on both sides.

###### 3.1.3.5.4 Posterior thalamic radiation (PoTR)

This bundle was considered to be the same as the extended optic radiation using the entire occipital lobe, and was not reconstructed separately but will be added in future versions.

#### 3.1.4 Cerebellar bundles *(in alphabetical order)*

##### 3.1.4.1 Dentato-rubro-thalamic tract (DRTT)

The DRTT is the main efferent pathway from the cerebellum to the cerebral cortex and an important subdivision of the SCP in the context of deep brain stimulation-based neurosurgery. It originates in the dentate nucleus of the cerebellum, ascends through the brain stem where the majority of its constituent axons cross the midline at the superior cerebellar decussation (SCP) to synapse with the contra-lateral red nucleus (RN) in the midbrain (Coenen et al., 2020; Kwon et al., 2011; Mollink et al., 2016; Nowacki et al., 2018), see figure 19. From here the DRTT continues to the ventrolateral (VL) and ventromedial (VM) nuclei of the thalamus. Technically the DRTT terminates in the thalamus; however, for functional completeness we extend its trajectory to terminate in the primary motor cortex.

**Figure 19:**
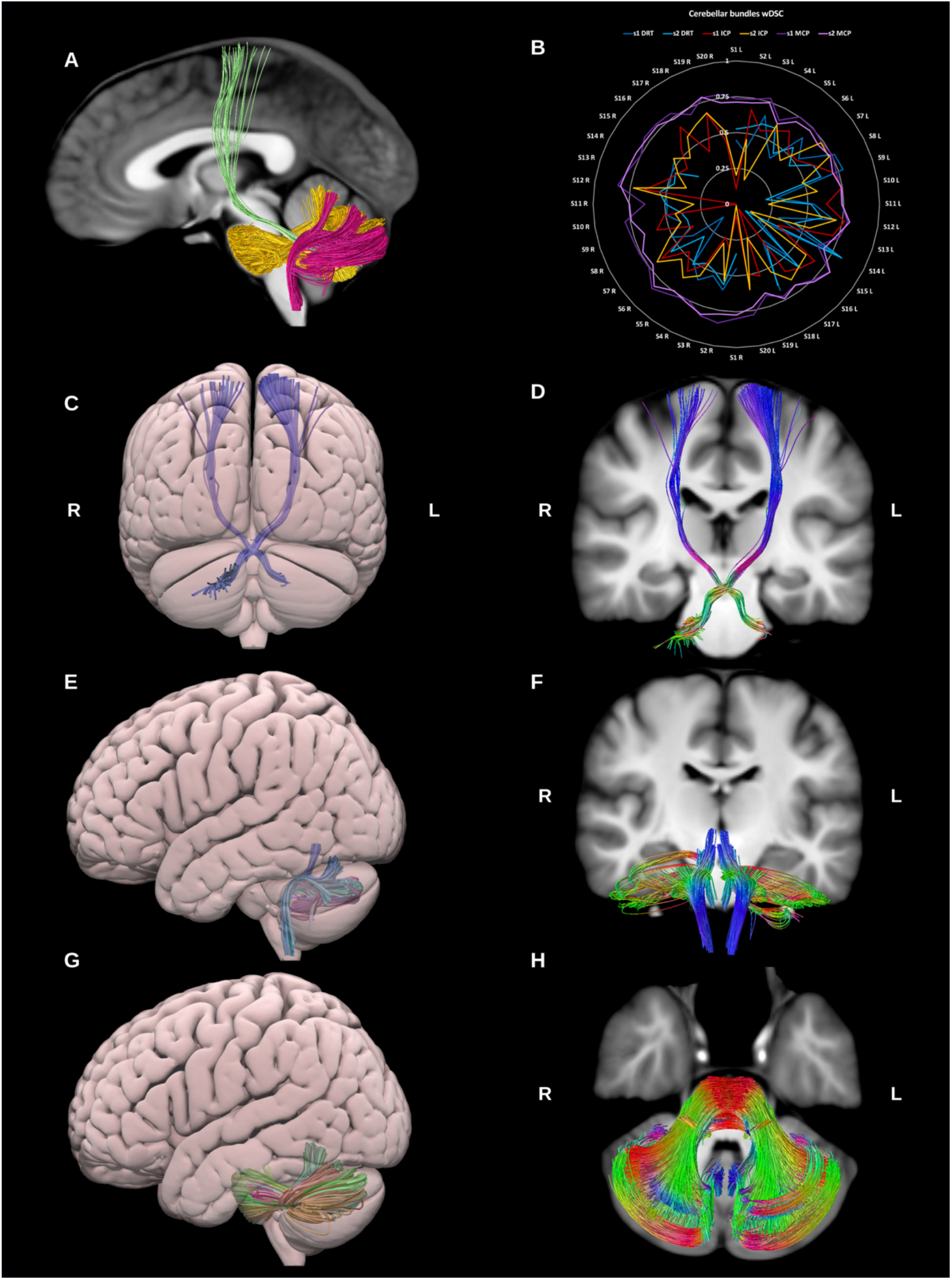
(A) shows a sagittal image of the T1-weighted group average image with the three cerebellar bundles shown in green for the **dentato-rubro-thalamic tract (DRTT)**, **inferior cerebellar peduncle (ICP)** in fuschia, and in gold the **middle cerebellar peduncle (MCP)**. (B) shows radar plots of wDSC (vertical range) for these three bundles per session. Left sided bundles (L) are on the first half of the circle and right ones (R) are on the second half. (C) and (D) shows a posterior projection surface view and a coronal slice with the DRTT overlaid. (E) and (F) show the ICPs shown on a lateral projection surface view, and a coronal slice respectively. (G) and (H) show the MCP in lateral projection surface view, and axial slice T1. wDSC = weighted dice similarity coefficient, MNI = Montreal Neurological Institute, missing results indicate failed tractography.

Whilst an ipsilateral component of the DRTT has been described in the literature, for the current purpose we have limited the reconstruction to the classical/contralateral DRTT. The left DRTT reconstruction failed in 4 datasets, while the right failed in 9 datasets. The overall DRTT wDSC showed a median of 0.498, minimum of 0.078 and Max.A.I.D. of 0.203. Additionally, we generated template DRTT bundles with 2,000 and 50,000 streamlines to illustrate the influence of the number streamline seeding attempts on tracking outcome and subsequent template-based reproducibility (supplementary figure 1).

##### 3.1.4.2 Inferior cerebellar peduncle (ICP)

The ICP consists mainly of afferent sensory fibers projecting from the spinal cord to the cerebellum. The ICPs are functionally involved in the maintenance of balance and posture via the integration of proprioceptive sensory and motor functions (Voogd, 2004). Microdissection studies reveal that it consists of four afferent bundles and 1 efferent bundle (Lingford-Hughes and Kalk, 2012), however here we reconstruct it as a single lateralized bundle using the dentate, fastigial and interposed nuclei of the cerebellum, and the ipsilateral medulla oblongata (Salamon et al., 2007; van Baarsen et al., 2016), see figure 19. The ICP was generated for all datasets on both sides.

##### 3.1.4.3 Middle cerebellar peduncle (MCP)

The MCPs are large paired bundles that connect the brainstem to the cerebellum on both sides. They are often reconstructed in neuroimaging studies as a single commissural bundle connecting both cerebellar hemispheres (Leitner et al., 2015). The MCPs are thought to play a role in the modulation of skilled manual motor functions (Lingford-Hughes and Kalk, 2012) and have been shown to consist of three sub-fascicles (superior, inferior and deep) by microdissection studies (Jhaveri et al., 2018). We reconstructed the MCP as a whole for each side using the contra-lateral pons and cerebellar cortex. The resulting streamlines cross the midline in the pons, in line with its neuroanatomical definition. (Re et al., 2017; van Baarsen et al., 2016; Voogd, 2004), see figure 19. The MCP was generated for all datasets on both sides.

### 3.2 HCP test-retest data results

In total we successfully reconstructed 2676 out of the 2720 attempted bundle reconstructions. The following bundles could not be reconstructed in all datasets: the DRTT (13 failures), premotor pyramidal tract (6 failures), optic tract (3 failures), fornix (15 failures), anterior commissure (2 failures), sensory CC (4 failures), and temporal CC (1 failure). Results of inter-session pair-wise similarity analysis per bundle for the different HCP test-retest subjects are shown as violin plots in figure 20.

**Figure 20:**
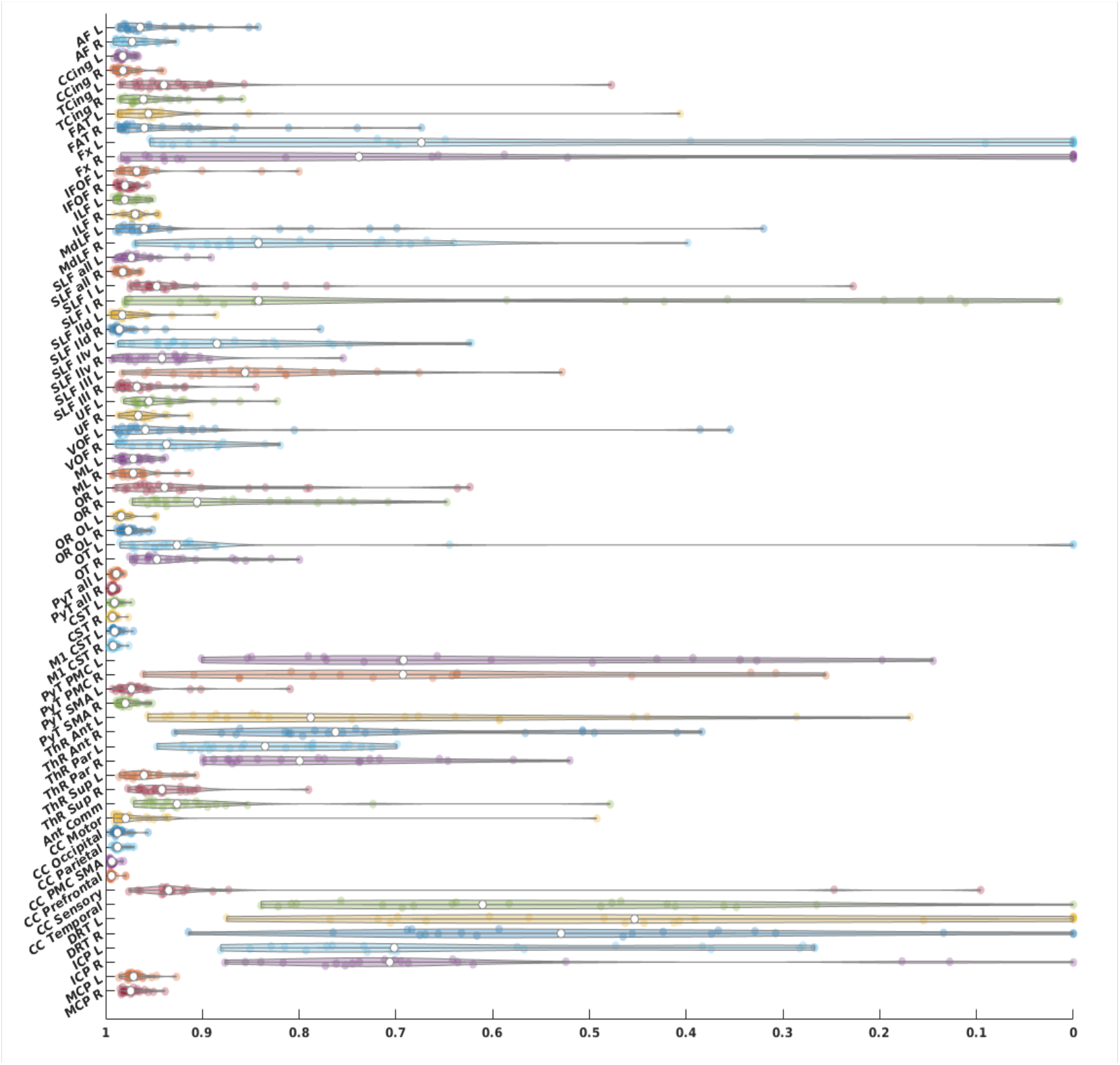
HCP tractograms wDSC scores depicted as violin plots. wDSC = weighted dice score, AF = arcuate fasciculus, CCing = cingulate cingulum, TCing = temporal cingulum, FAT = frontal aslant tract, Fx = fornix, IFOF = inferior fronto-occipital fasciculu, ILF = inferior longitudinal fasciculus, MdLF = middle longitudinal fasciculus, SLF = superior longitudinal fasciculus, SLF-IId = SLF-II dorsal division, SLF-IIv = SLF-II ventral division, UF = uncinate fasciculus, VOF = vertical occipital fasciculus, ML = medial lemniscus, OR = optic radiation, occlobe = occipital lobe, OT = optic tract, PyT = pyramidal tract, CST = corticospinal tract, M1 = primary motor cortex, PyT = pyramidal tract, PMC = premotor cortex, SMA = supplementary motor area, ThR = thalamic radiation, Ant = anterior, Par = parietal, Sup = superior, Ant Comm = anterior commissure, CC = corpus callosum, PMC and SMA = premotor cortex and supplementary motor area, DRT = dentatorubro-thalamic tract, ICP = inferior cerebellar peduncle, MCP = middle cerebellar peduncle, L = left, R = right.

Voxel-wise binary union maps per bundle show an expected pattern of maximum agreement in bundle cores and maximum variability around the periphery.

Figures 21 & 22 demonstrate these maps for the AF and CST, and fornix and DRTT, respectively as examples of some of the best and worst performing bundles in terms of similarity to HCP-template and inter-session reproducibility. Equivalent maps are provided for all other bundles in the supplementary material.

**Figure 21:**
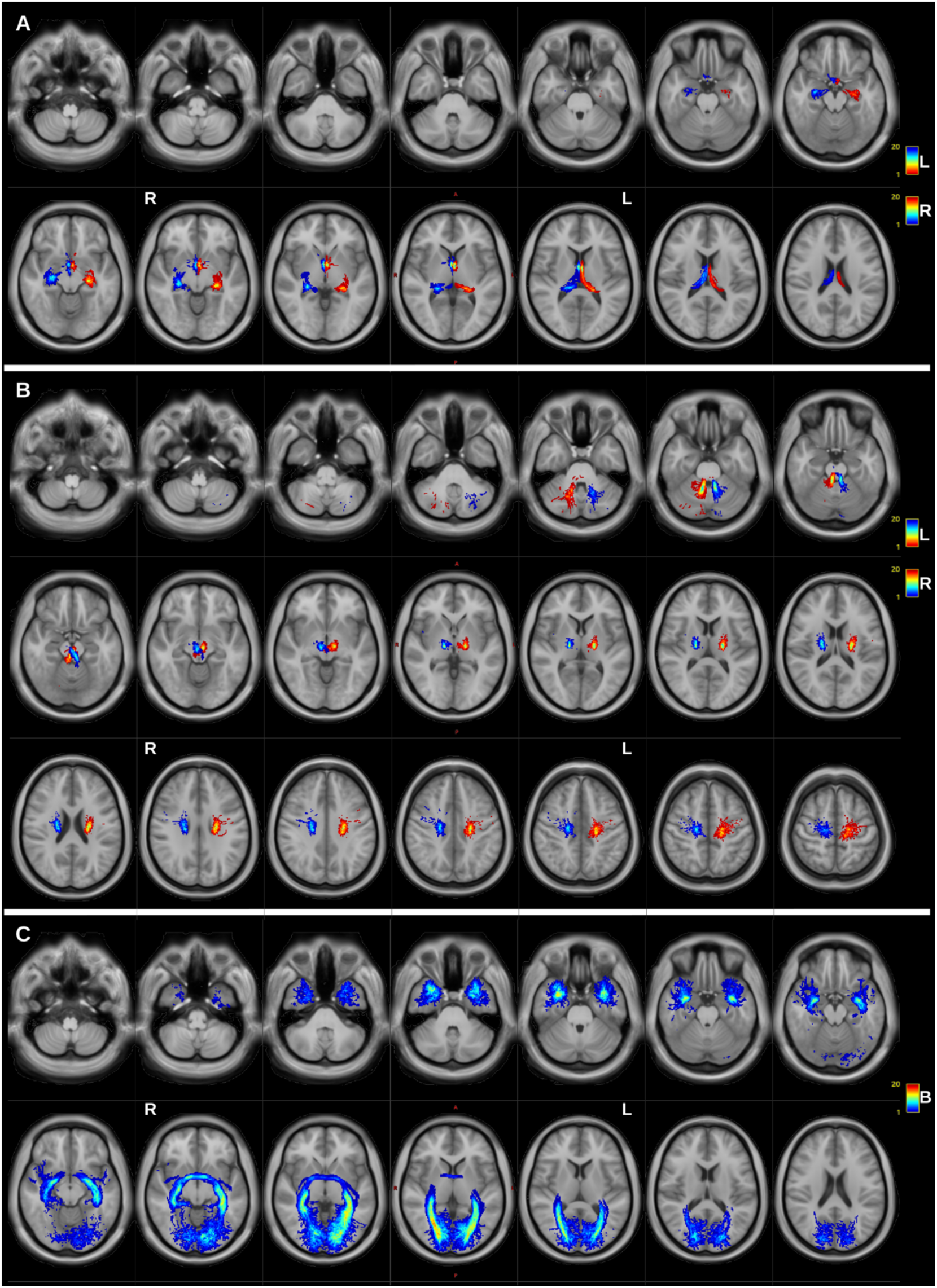
Serial axial slices of the HCP-template T1-weighted images with the cumulative voxelwise heat-maps of the fornix (A), dentato-rubro-thalamic tracts (B), and anterior commissure (C), L = left, R = right, B = bilateral. Left sided bundles are shown in red-to-blue, and right sided bundles are shown in blue-to-red.

**Figure 22:**
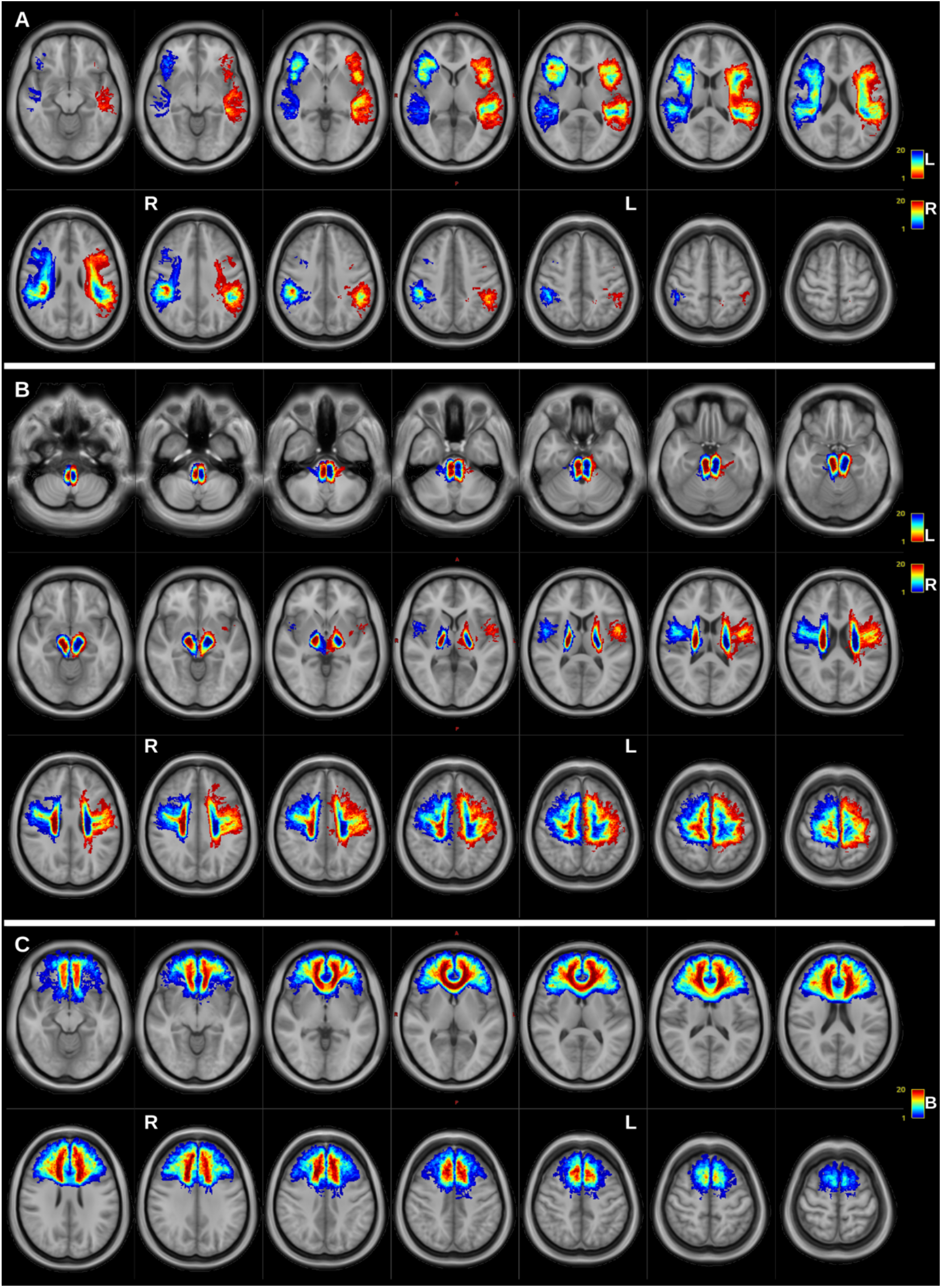
Serial axial slices of the HCP-template T1-weighted images with the cumulative voxelwise heat-maps of the arcuate fasciculi (A), whole pyramidal tracts (B), and prefrontal corpus callosum (C), L = left, R = right, B = bilateral. Left side bundles are shown in red-to-blue, and right sided bundles are shown in blue-to-red.

### 3.3 MASSIVE data results

In total we successfully reconstructed 316 out of 340 bundles (68 for each of the 5 MASSIVE datasets). 63 out of the 68 bundles were generated for every dataset, while the anterior commissure was only generated in 1 dataset, and the DRTT and fornix were not generated in any dataset. Figure 23 shows the wDSC scores derived from comparing the MASSIVE bundles to the HCP-template bundles superimposed on the results of the HCP test-retest dataset for reference. The single score agreement ICC score was 0.917 (upper-bound = 0.944, lower-bound = 0.884, p < 0.05).

**Figure 23:**
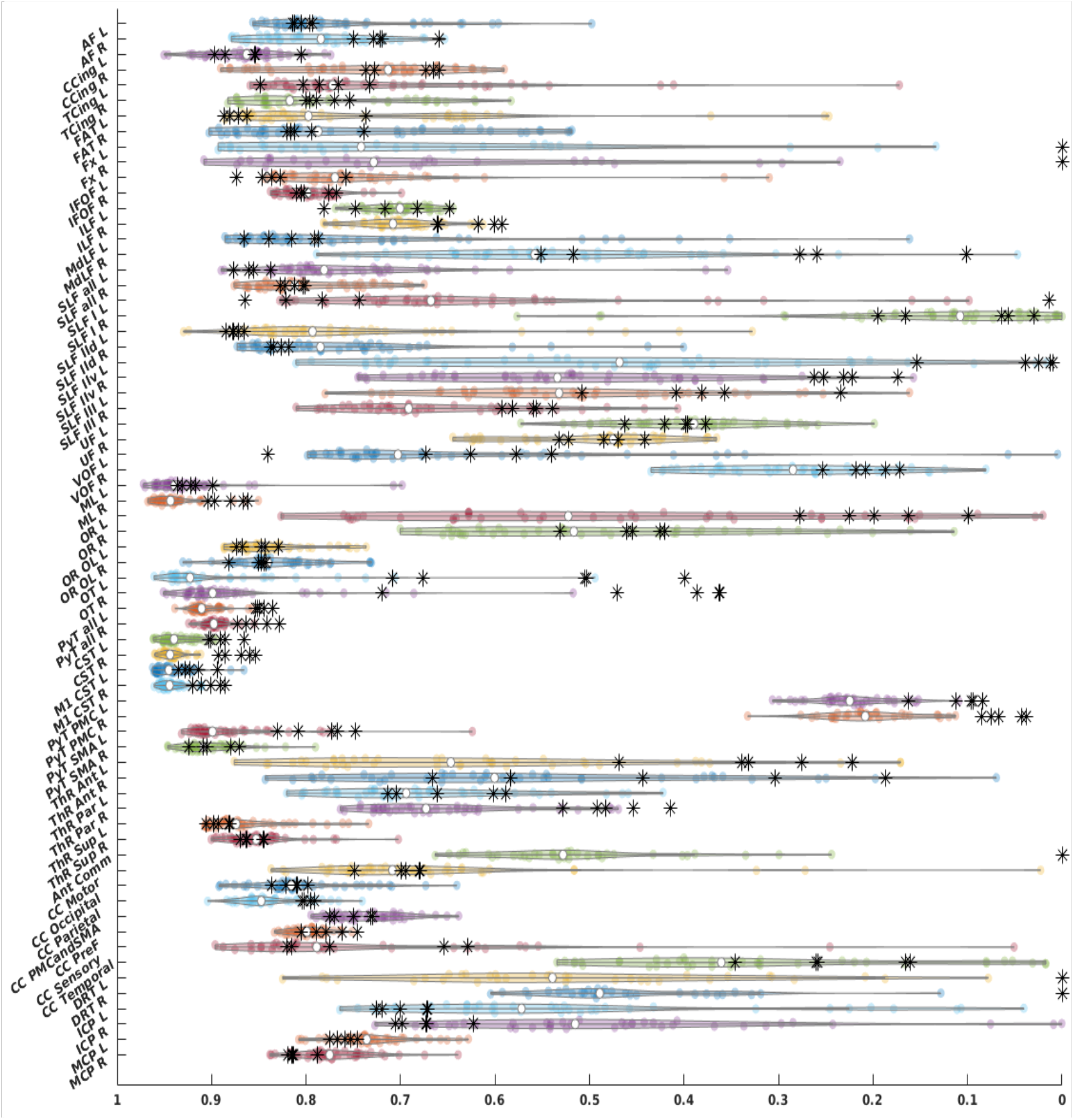
Weighted dice similarity coefficient scores for all bundles, HCP test-retest tractograms are depicted as violin plots and MASSIVE results are depicted as black asterisks., AF = arcuate fasciculus, CCing = cingulate cingulum, TCing = temporal cingulum, FAT = frontal aslant tract, Fx = fornix, IFOF = inferior fronto-occipital fasciculu, ILF = inferior longitudinal fasciculus, MdLF = middle longitudinal fasciculus, SLF = superior longitudinal fasciculus, SLF-IId = SLF-II dorsal division, SLF-IIv = SLF-II ventral division, UF = uncinate fasciculus, VOF = vertical occipital fasciculus, ML = medial lemniscus, OR = optic radiation, occlobe = occipital lobe, OT = optic tract, PyT = pyramidal tract, CST = corticospinal tract, M1 = primary motor cortex, PyT = pyramidal tract, PMC = premotor cortex, SMA = supplementary motor area, ThR = thalamic radiation, Ant = anterior, Par = parietal, Sup = superior, Ant Comm = anterior commissure, CC = corpus callosum, PMC and SMA = premotor cortex and supplementary motor area, DRT = dentato-rubro-thalamic tract, ICP = inferior cerebellar peduncle, MCP = middle cerebellar peduncle, L = left, R = right.

## 4 Discussion

Human WM segmentation based on diffusion imaging tractography has been likened to blurring the boundary between science and art (Van Hecke et al., 2016). While few would argue about the aesthetic qualities of these virtual dissections, when applied appropriately, they transcend pretty pictures to become an indispensable tool in modern neurosurgical practice and neuroscience research (Johansen-Berg and Behrens, 2006; Panesar et al., 2019). In this work we aim to facilitate the use of advanced tractography methods within the clinical research community. First, we provide an intuitive educational reference for researchers and clinicians wishing to apply CSD tractography to typical HARDI datasets. Second, we describe typical WM anatomy derived from automated subject-specific parcellation driven CSD probabilistic tractography in adults. Third, we demonstrate the variability of the reconstructed bundles within and between subjects using two test-retest datasets and offer solutions to improve tracking results for the least reproducible bundles. Finally, we provide an open-source fully-automated and customizable tractography pipeline (FWT) enabling researchers and clinicians to apply the methods presented in this work to their own data.

### 4.1 Variability of virtual dissections

Our results demonstrate that inter-bundle variability was much greater than inter-subject variability for the same bundles i.e., there was a wide range of similarity in the extent to which different bundles were reconstructed irrespective of the dataset used. The AF and PyT for example were reconstructed in largely the same way in all datasets, whereas the fornix and DRTT showed high variability in density and spatial extent and were sometimes not reconstructed at all. In line with this, inter-subject agreement was highest for bundles with high streamline densities e.g., PyT, AF, CC and lowest for bundles with low streamline densities e.g., fornix, and DRTT.

Inter-session and intra-subject agreement were high for all bundles i.e., bundles were reconstructed largely to the same extent, (or failed to be reconstructed) in both sessions of the HCP test-retest dataset. In the test-retest data from a single-subject with varying b-values and shell schemes (MASSIVE) changing the b-value(s), number of gradient directions or shell scheme did not consistently alter bundle similarity to the model.

These findings align with a large body of evidence that diffusion tractography reconstructions are highly variable and are influenced not only by data acquisition parameters such as scanner manufacturer and field strength, b-value, diffusion sampling scheme, reconstruction model (Schilling et al., 2021), but also by the virtual dissection protocol itself (Schilling et al., 2020). Notably, in the context of our study, despite using exactly the same WM dissection protocol for the HCP-template as for the individual datasets, none of the tracked bundles failed in the template data. This highlights the importance of data quality, as group-averaged data tends to have a much higher signal-to-noise ratio (SNR) despite increased blurriness in fine cortical structures due to inter-subject variability. It also demonstrates that even in a unified fully automated WM dissection protocol considerable inter-bundle and inter-subject variability remain. In contrast to previous work, we found that data acquisition parameters and sampling scheme had only a limited impact on bundle variability. This is likely due to the high number of volumes used per session, which would result in a higher SNR compared to the typical single acquisition datasets used in comparison studies.

In agreement with other studies (Bonilha et al., 2015; Cousineau et al., 2017; Gu et al., 2017; Zhang et al., 2019), we found that larger bundles, particularly if they have both a larger volume and a higher streamline count, tend to be more similar/reproducible regardless of differences in scanning parameters or even normal inter-subject anatomical variations. It follows that automated procedures are most appropriate for fasciculi with these characteristics. For smaller, less dense or more narrow bundles such as the fornix, DRTT (and relatedly, the superior cerebellar peduncle) or specific subdivisions of larger bundles (eg SLF-I, temporal corpus callosum) we recommend using either data with sub-millimeter spatial resolution, bundle-specific manual dissection or using the template bundles we provide in an automated in a RecoBundles (Garyfallidis et al., 2018) based workflow rather than the VOI-based approach. Of note, we also provide symmetrical versions of the HCP-template bundles along with aligned reference anatomical images. In line with the use of template bundles, FWT is compatible with RecoBundles (Garyfallidis et al., 2018) based pipelines that can overcome the need for structural parcellation, which can be particularly useful in clinical data to avoid false tractography results due to structural parcellation errors.

### 4.2 The benefits of using a CSD atlas in clinical studies

There are currently a number of diffusion-based tractography atlases and protocols available (Catani and Thiebaut de Schotten, 2008; Thiebaut de Schotten et al., 2011; Verhoeven et al., 2010; Wakana et al., 2004; Warrington et al., 2020; Wasserthal et al., 2018; Yendiki et al., 2011) some of which are based on HCP data (Hansen et al., 2020; Yeh et al., 2018). Here we aimed to primarily address the limitations of DTI based virtual dissection protocols, which predominate in clinical studies, either as part of comparative studies with healthy controls, or in a healthcare setting using data acquired in shorter scan times than are achievable in a research environment. We did not include formal comparison to other HARDI/higher-order modelling-based atlases which precludes us from drawing conclusions about the relative benefits of FWT over other methods for virtual dissection in this context. It is our aim to complement atlases based on detailed anatomical research and specialized acquisition schemes with a comprehensive description of CSD-based human WM anatomy which improves upon classical DTI representations in 3T data. Whilst the results presented here are based on high quality data in healthy subjects, we also tested our protocol on lower quality, neurosurgical patient data during the development phase of FWT, to ensure its translatability to clinical populations. A formal analysis relating to the application of FWT in patients with lesions is beyond the scope of this paper focusing on typical CSD anatomy and will be presented in a follow-up publication.

### 4.3 Technical considerations and limitations

This work relied on imaging data from only 20 individuals, however by using re-test scans yielding 40 datasets and the additional analysis using the MASSIVE dataset, our sample is sufficient to study test-retest reproducibility, as well as inter-subject variability. While our dissection protocol was literature-based and guided by several contemporary publications we did not include a direct comparison to any other atlas, nor did we apply formal specific criteria for assessing the validity of the definitions. Given the lack of consensus on anatomical definitions relating to WM fasciculi, it is possible that some readers will disagree with our interpretation, and that the FWT VOIs and reconstructions may need to be revised as the field evolves.

We attempted to keep rigid heuristic decisions (e.g., statistical thresholds) to a minimum especially those pertaining to streamlines filtering, where we utilized largely data-driven streamline filtering tools from Dipy (Garyfallidis et al., 2014) rather than relying on a more stringent selection of anatomical exclude VOIs. However, in order to cater to varying data quality, our choice of inclusion VOIs in some situations are influenced by considerations made for lower quality datasets than the ones utilized in this work. For example, for the AF we excluded any streamlines involving the precentral gyrus to avoid false streamlines tunneling through volume average voxels, which we have observed in lower quality data single-shell data. Similarly, the AC is generated without the olfactory component, and we do not include subdivisions to the AF, CST, full posterior thalamic radiation, or superior cerebellar peduncles proper. We also do not include more esoteric bundles such as the superior-anterior fasciculus (David et al., 2019), temporo-insular fasciculus (Nachtergaele et al., 2019; Radwan et al., 2019), and inferior (auditory) thalamic radiation (Maffei et al., 2019a, 2019b), or those with a high chance of failure due to their geometry, such as the posterior commissure, or cranial nerves.

FWT tractography utilizes spherical-deconvolution informed filtering of tractograms (SIFT) (Smith et al., 2015, 2013)) based streamlines seeding and filtering for whole brain tractography, while further improvement can be expected if an optimized whole brain tractography method is utilized, e.g., global tractography (Christiaens et al., 2015) or particle-filtering tractography (Girard et al., 2014). FWT does not use a specific required number of output streamlines during bundle segmentation from whole-brain tractograms to avoid biasing the outcome by an expected number of streamlines. However, this is unavoidable if the bundle specific approach is utilized. In this case the resulting bundle will be directly influenced by the required number of streamlines as this will influence the number of streamlines seeding attempts. This point is illustrated in supplementary figure 1, which shows that when the DRT template bundles are generated twice with 2,000 and 50,000 required streamlines initially, the outcome from the 50,000 streamlines appears more complete as can be expected. Furthermore, when applied to the individual datasets, the bundle-specific approach with 20,000 required streamlines reconstructed more bundles successfully than the whole-brain tractography and bundle segmentation approach (73 vs. 67), and resulted in higher wDSC scores particularly when compared to the 50,000 streamlines DRT template bundles.

The FWT pipeline is time-consuming, as FS recon-all alone requires at least 4 - 6 hours and dMRI preprocessing can vary widely depending on acquisition parameters, and reconstruction method. E.g., whole brain tractography can take more than 4 hours for a multi-shell multi-tissue CSD model with probabilistic tractography using second-order integration over distributions of fODFs (iFOD2) (Tournier et al., 2010) with fODF-driven dynamic seeding (Smith et al., 2015). Whilst this is offset by the flexibility of FWT allowing a more targeted approach using bundle-specific seeding and tractography in case of presurgical mapping, further development will focus on improving processing time. Additionally, future work will provide a clustering-based workflow independent of prior structural parcellation, the addition of more fiber bundles, as well as generating deterministic versions of all bundles. Finally, any justified and necessary changes to the inclusion/exclusion VOIs for any bundle can be easily implemented by changing the FWT workflows, which are openly provided, and a new atlas of bundles can be easily created by rerunning the template workflows on the HCP-group template data.

## 5 Conclusion

The FWT pipeline can reconstruct 68 WM fasciculi and shows high inter- and intra-subject reproducibility. Dense bundles such as the PyT, AF and MCP yield the most reliable and least variable reconstructions. Higher resolution data or tract specific modification may be required for thinner bundles such as the fornix, DRTT, SLF I and premotor PyT. The FWT CSD atlas may be a useful reference and virtual dissection tool for applied neuroimaging students and clinical professionals wishing to use and understand the capabilities of CSD tractography.

## Supporting information

accompanying supplementary information document

## Acknowledgments and disclosures

Kurt Schilling, and Bennet Landman are supported by the National Institutes of Health under award number 2R01EB017230. Maxime Descoteaux is the Institutional Research Chair in NeuroInformatics of Université de Sherbrooke, and is funded by the Natural Sciences and Engineering Research Council of Canada (NSERC) discovery grants. Tom Theys is a senior clinical researcher supported by the Research Foundation – Flanders (FWO) (FWO grant 1830717N) Louise Emsell and Mathieu Vandenbulcke are supported by FWO (FWO grant G0C0319N), KU Leuven research fund (C24/18/095) and the Sequoia Fund for Research on Ageing and Mental Health.

## Conflicts of interest

Maxime Descoteaux is a founder and co-owner of Imeka Solutions Inc. No conflicts related to the current work.

## Data sharing statement

All data used in this work can be recovered from their respective online sources mentioned above. The accompanying supplementary information document contains more detail on the datasets used in this work, further explanation of the FWT method, and additional similarity analysis results. The FWT workflows can be downloaded from (https://github.com/KUL-Radneuron/KUL_FWT.git). Finally, the Open Science Framework repository (https://osf.io/snq2d/) contains additional screenshots of single subject bundles, bundle heatmaps, original and symmetric versions of the HCP-template atlas bundles, and additional tractograms.

## List of abbreviations

AC: Anterior Commissure
AC/PC: Anterior Commissure/Posterior Commissure
AF: Arcuate Fasciculus
AP: Anterior-Posterior
ATR: Anterior Thalamic Radiation
BIDS: Brain Imaging Data Structure
CC: Corpus Callosum
CG: Cingulum
CSD: Constrained Spherical Deconvolution
CST: CorticoSpinal Tract
DL: DorsoLateral nucleus of the thalamus
DM: DorsoMedial nucleus of the thalamus
dMRI: Diffusion Magnetic Resonance imaging
DRTT: Dentato-Rubro-Thalamic Tract
DSC: Dice Similarity Coefficient
DSI: Diffusion Spectral Imaging
DTI: Diffusion Tensor Imaging
FA: Fractional Anisotropy
FACT: Fiber Assignment by Continuous Tracking
FAT: Frontal Aslant Tract
fODF: Fiber Orientation Distribution Function
FS: FreeSurfer
FT: Fiber Tracking/Tractography
FWT: Fun With Tracts
Fx: Fornix
HARDI: High Angular Resolution Diffusion Imaging
HCP: Human Connectome Project
ICC: Intra-Class Correlation
ICP: Inferior Cerebellar Peduncle
IFOF: Inferior Fronto-Occipital Fasciculus
ILF: Inferior Longitudinal Fasciculus
M1: Primary Motor cortex
MALP-EM: Multi-Atlas Label Propagation with Expectation Maximization
MASSIVE: Multiple Acquisitions for Standardization of Structural Imaging Validation and Evaluation
MCP: Middle Cerebellar Peduncle
MdLF: Middle Longitudinal Fasciculus
ML: Medial Lemniscus
MR: Magnetic Resonance
MRI: Magnetic Resonance Imaging
MSBP: MultiScale Brain Parcellator
NIST: Neuroimaging and Surgical Technologies
OR: Optic Radiation
OT: Optic Tract
PaTR: Parietal Thalamic Radiation
PD25: Parkinson’s Disease 25 subjects histological atlas
PoTR: Posterior Thalamic Radiation
PyT: Pyramidal Tract
ROIs: Regions Of Interest
SLF: Superior Longitudinal Fasciculus
SNR: Signal-to-Noise Ratio
STR: Superior Thalamic Radiation
SUIT: Spatially UnbIased atlas Template of the cerebellum and brainstem
UF: Uncinate Fasciculus
UKBB: United Kingdom BioBank
VOF: Vertical Occipital Fasciculus
VOIs: Volumes of interest
VPL: Ventral Postero-Lateral
VPM: Ventral Postero-Medial
wDSC: Weighted Dice Similarity Coefficient
WM: White Matter

## Notes

https://osf.io/snq2d/

https://github.com/KUL-Radneuron/KUL_FWT.git

### Competing Interest Statement

The authors have declared no competing interest.

### Summary of Updates

Fixed some minor spacing issues on page 14 and last two sections

