## Supplementary material for "An atlas of white matter anatomy, its variability, and reproducibility based on Constrained Spherical Deconvolution of diffusion MRI": accompanying supplementary information document

#### List of abbreviations:

**AC** = Anterior Commissure
**AC/PC** = Anterior Commissure/Posterior Commissure
**AF** = Arcuate Fasciculus
**AP** = Anterior-Posterior
**ATR** = Anterior Thalamic Radiation
**BIDS** = Brain Imaging Data Structure
**CC** = Corpus Callosum
**CG** = Cingulum
**CSD** = Constrained Spherical Deconvolution
**CST** = CorticoSpinal Tract
**DL** = DorsoLateral nucleus of the thalamus
**DM** = DorsoMedial nucleus of the thalamus
**dMRI** = Diffusion Magnetic Resonance imaging
**DRTT** = Dentato-Rubro-Thalamic Tract
**DSC** = Dice Similarity Coefficient
**DSI** = Diffusion Spectral Imaging
**DTI** = Diffusion Tensor Imaging
**FA** = Fractional Anisotropy
**FACT** = Fiber Assignment by Continuous Tracking
**FAT** = Frontal Aslant Tract
**FBC** = Fiber to Bundle Coherence
**fODF** = Fiber Orientation Distribution Function
**FS** = FreeSurfer
**FT** = Fiber Tracking/Tractography
**FWT** = Fun With Tracts
**Fx** = Fornix
**HARDI** = High Angular Resolution Diffusion Imaging
**HCP** = Human Connectome Project
**ICC** = Intra-Class Correlation
**ICP** = Inferior Cerebellar Peduncle
**IFOF** = Inferior Fronto-Occipital Fasciculus
**ILF** = Inferior Longitudinal Fasciculus
**M1** = Primary Motor cortex
**MALP-EM** = Multi-Atlas Label Propagation with Expectation Maximization
**MASSIVE** = Multiple Acquisitions for Standardization of Structural Imaging Validation and Evaluation
**MCP** = Middle Cerebellar Peduncle
**MdLF** = Middle Longitudinal Fasciculus
**ML** = Medial Lemniscus
**MR** = Magnetic Resonance
**MRI** = Magnetic Resonance Imaging
**MSBP** = MultiScale Brain Parcellator
**NIST** = Neuroimaging and Surgical Technologies
**OR** = Optic Radiation
**OT** = Optic Tract
**PaTR** = Parietal Thalamic Radiation
**PD25** = Parkinson’s Disease 25 subjects histological atlas
**PoTR** = Posterior Thalamic Radiation
**PyT** = Pyramidal Tract
**ROIs** = Regions Of Interest
**SLF** = Superior Longitudinal Fasciculus
**SNR** = Signal-to-Noise Ratio
**STR** = Superior Thalamic Radiation
**SUIT** = Spatially Unbiased atlas Template of the cerebellum and brainstem
**UF** = Uncinate Fasciculus
**UKBB** = United Kingdom BioBank
**VOF** = Vertical Occipital Fasciculus
**VOIs** = Volumes of interest
**VPL** = Ventral Postero-Lateral
**VPM** = Ventral Postero-Medial
**wDSC** = Weighted Dice Similarity Coefficient
**WM** = White Matter

### Methodology

#### Additional HCP subject details

Supplementary table 1 shows the HCP identification codes of each HCP test-retest subject, along with gender and age-group.

| **Supplementary table 1 HCP test-retest subject details** | | | | | | | |
|---|---|---|---|---|---|---|---|
| Code | ID | Gender | Age-group | Code | ID | Gender | Age-group |
| S1 | 103818 | F | 31-35 | S11 | 122317 | M | 31-35 |
| S2 | 111312 | F | 31-35 | S12 | 139839 | M | 26-30 |
| S3 | 115320 | F | 31-35 | S13 | 146129 | M | 22-25 |
| S4 | 125525 | F | 31-35 | S14 | 149337 | M | 31-35 |
| S5 | 135528 | F | 31-35 | S15 | 149741 | M | 26-30 |
| S6 | 137128 | F | 31-35 | S16 | 151526 | M | 26-30 |
| S7 | 158035 | F | 26-30 | S17 | 185442 | M | 22-25 |
| S8 | 172332 | F | 26-30 | S18 | 599671 | M | 26-30 |
| S9 | 177746 | F | 26-30 | S19 | 122317 | M | 31-35 |
| S10 | 660951 | F | 26-30 | S20 | 139839 | M | 26-30 |

#### FWT technical description

FWT is a collection of open-source workflows in Bash shell and python3 relying on state-of-the-art image registration and anatomical parcellation pipelines (e.g., FreeSurfer and MSBP) to perform automated diffusion tractography. We provide the user with two main approaches (A) local/targeted fiber dissection, which aims to reconstruct only the bundle of interest, and (B) whole brain tractography followed by virtual bundle dissection. Both approaches are fully automated and use exactly the same set of dissection VOIs. In addition, the user can choose between DTI and CSD models combined with different tractography algorithms. FWT also provides the option of generating automated screenshots of each bundle and computing bundle-profile measures given in quantitative (as .csv files) and qualitative forms (as bundle-profile plots).

FWT relies on MRTrix3 v3.0.2-78 (Tournier et al., 2019), FreeSurfer v6.0 (Fischl, 2012) , FSL v6.0.4 (Jenkinson et al., 2012) , ANTs v2.3.1 (Avants et al., 2011) , Dipy v1.3.0 (Garyfallidis et al., 2014) and Scilpy (Bore et al., 2021; "Scilpy documentation," 2021) v.1.1.0. The pipeline is divided into two main parts, both of which are driven by a configuration text file that specifies the bundles to be generated and, if the user chooses the bundle specific approach, the number of streamlines per bundle. The two parts of FWT are detailed below:

##### Part 1

The script for this part is named KUL_FWT_generate_VOIs.sh for individual datasets and its counterpart for group-averaged template data is KUL_FWT_generate_VOIs_4temp.sh. These are responsible for generating all dissection VOIs as well as basic DTI parametric maps (FA and MD maps) with MRTrix3 dwi2tensor and tensor2metric for registration and quality assurance purposes.

Dissection VOIs are generated by combining different parcels or VOIs from the same or different parcellation maps. FWT uses the following from FreeSurfer: the parcellation map aparc+aseg.mgz and to a lesser extent the DKT2009aparc+aseg.mgz, the segmentation maps aseg.mgz, the fornix segmentation map generated by the FreeSurfer script mri_cc, the lobes segmentation map generated using a combination of mri_annotation2label and mri_aparc2aseg, and the WM parcellation map wmparc.mgz. From MSBP FWT uses only the scale-3 parcellation maps with 234 parcels plus the thalamic nuclei and brainstem segmentations. In addition, it relies on the following volumetric maps in MNI152 space: the NIST PD25 histological atlas, the SUIT cerebellar atlas, the UKBB derived brainstem volumetric tracts, the Jülich university histological atlas, the Johns Hopkins white matter labels atlas, and manually defined midline VOIs for the AC, PC, and right and left hemispheres.

All FreeSurfer and MSBP VOIs are warped directly to diffusion space using antsIntermodalityRegistration.sh with the FA map as the target for individual datasets and the first volume of the fODF as the target for group averaged data. This is preceded by a non-linear T1-weighted image normalization to the UKBB T1 group template in MNI152 space using antsRegistrationSyN.sh, and the resulting transform and warp field are used to drive the intermodality registration step. The resulting inter-modality transform and backward and forward warp fields are applied directly to the label maps using antsApplyTransforms.sh with multi-label interpolation to avoid label aliasing. This allows an accurate transfer of the anatomical label maps to diffusion space while minimizing errors from multiple interpolations.

Next the script will combine different anatomical VOIs depending on a hard-coded recipe specified as a string array of anatomical VOI names for each bundle and an array of voxel values per labels for each source parcellation or segmentation map. Each anatomical VOI has a suffix (e.g., _FS for FreeSurfer aparc+aseg.mgz derived parcels, _MSBP for MSBP derived and _UKBB for UKBB derived, etc.). To streamline the use of white matter labels FS derived VOIs include _WM_ or _GM_ in their names. This setup is easy to read for novice computational image analysis users and easy to transfer to different formats, e.g., a json based config file to avoid hardcoding problems. Include and exclusion VOIs are generated using MRTrix3 tools, e.g., mrcalc, maskfilter, and mrfilter, as well as fsl tools e.g., fslmaths. Intermodality and normalization transforms and warps are also converted to an MRTrix compatible format using warpinit, antsApplyTransforms and warpcorrect as described on the MRTrix3 user forum (Pietsch, 2020). MSBP and JHU maps are used to subsegment some FS derived VOIs, e.g., the unsegmented WM labels, as well as other WM labels that aid in fine dissection of complex regions (e.g., the temporal stem, internal and external capsules) while minimizing reliance on manually defined VOIs. This is done using a combination of the ANTs (Avants et al., 2011) script ImageMath PropagateLabelsThroughMask functionality, which allows a multi-label map to be propagated through a different binary mask, thus allowing the subsegmentation of white matter based on grey matter parcellations for example, or the modification of one atlas based on information from another.

Inclusion and exclusion VOIs per bundle are generally kept to the lowest number possible while being anatomically valid, by adding and binarizing different VOIs, e.g., the AF has two includes, a temporo-parietal one and a frontal one, each consisting of multiple constituent anatomical parcels derived from the aparc+aseg and

MSBP maps, while the ML has three includes, the brainstem ML VOI, the thalamus and the primary sensory cortex. Multi-label maps are also generated for each include and exclude VOI, and all dissection VOIs and maps are generated in native diffusion space and in MNI152 space.

##### Part 2

The script for this part is named KUL_FWT_generate_TCKs.sh for individual datasets and its counterpart for group-averaged template data is KUL_FWT_generate_TCKs_4temp.sh. These workflows are responsible for applying the dissection protocol and for tracking and filtering all the fiber bundles using the include and exclusion VOIs for each bundle generated by part 1. This part of FWT can automatically generate screenshots of each bundle and compute bundle-profile measures given in quantitative (as .csv files) and qualitative forms (as bundle-profile plots). If the user specifies the bundle-specific tractography approach, the MRTrix3 (Tournier et al., 2019) command tckgen for local tracking will use all the inclusion VOIs simultaneously as seeding and inclusion VOIs to create the initial version of a bundle. If the user specifies the second approach (whole brain tractography followed by bundle segmentation), then tckgen will be used for whole brain tractography using the entire brain mask for seeding to generate 10 million streamlines by default for individual datasets and 20 million for group-averaged data. The initial version of a bundle is then segmented using TCKedit.

Both approaches employ an angle threshold of 45 degrees (a value established by empirical testing of 30, 45, 60 and 90 degrees), a minimum fiber length of 20 mm and a maximum of 280 mm, and default thresholds for fODF amplitude and step-size. Initial versions of bundles are then filtered using the ScilPy (Bore et al., 2021) streamline filtering tools scil_filter_tractogram.py, scil_detect_streamlines_loops.py, and scil_outlier_rejection.py to remove false-positives from the initial bundles. This sequence of bundle cleanup steps is applied to all bundles except for the optic pathway bundles, which are filtered using the Dipy (Garyfallidis et al., 2014) implementation of Fiber-to-bundle coherence (Meesters et al., 2017). Filtered bundles are resampled to 1 mm step-size and smoothed using scil_smooth_streamlines.py with a Gaussian kernel applied to the coordinate array.

Optional workflows are also included for creating multiplanar screenshots of the filtered bundles using custom python scripts based on Dipy, as well the Scilpy scripts scil_screenshot_bundle.py, and scil_visualize_bundles_mosaic.py. Finally, if specified FWT can also generate quantitative and qualitative bundle-profile measures. MRTrix3 (tensor2metric) is used to calculate additional DTI-based parametric maps (AD, and RD), CSD-based parametric maps (AFD (Raffelt et al., 2012), Dispersion, and number fODF of peaks) are calculated using MRTrix3 fod2fixel and fixel2voxel, and streamline based voxel-wise parametric maps (length (Pannek et al., 2011), curvature and track-density (Calamante et al., 2010)) are calculated using tckmap. Next, bundles are resampled to 101 points using MRTrix3 tckresample, then a centroid streamline using scil_compute_centroid.py. The bundle is then explicitly reoriented in python using the Dipy function dipy.tracking.streamline.orient_by_rois according to the order of inclusion VOIs used to generate the bundle. Scilpy's scil_compute_bundle_voxel_label_map.py is used to create a voxel-wise segmentation of the bundle of interest into 100 bands/segments (No. of points of resampled tractogram – 1) which are used to calculate the bundle profiles. Bundle profiles are generated for all available voxel-wise metrics using MRTrix3 tcksample and plotted in python as well as connectome fingerprint maps for each bundle. Supplementary table 1 lists the tracked bundles, initial required number of streamlines, and end-result number of streamlines.

**Supplementary table 1: HCP template bundles reconstructed using bundle-specific seeding and probabilistic tractography, required and end-result number of streamlines per bundle.**

| Fiber bundles | Required number of streamlines | End-result number of streamlines | |
|---|---|---|---|
| | | Left | Right |
| **Arcuate fasciculus** | 8,000 | 7,285 | 4,013 |
| **Cingulate cingulum** | 8,000 | 7,022 | 1,932 |
| **Parahippocampal cingulum** | 4,000 | 3,023 | 2,993 |
| **Fornix** | 2,000 | 1,269 | 1,470 |
| **Frontal aslant tract** | 6,000 | 5,825 | 5,668 |

| | | | |
|---|---|---|---|
| **Inferior fronto-occipital fasciculus** | 6,000 | 5,499 | 2,967 |
| **Inferior longitudinal fasciculus** | 4,000 | 3,280 | 3,844 |
| **Middle longitudinal fasciculus** | 6,000 | 5,550 | 832 |
| **Whole superior longitudinal fasciculus (SLF)** | 18,000 | 16,759 | 14,895 |
| **SLF-I** | 4,000 | 3,952 | 118 |
| **SLF-IIv** | 6,000 | 5,868 | 5,005 |
| **SLF-IId** | 6,000 | 5,890 | 5,842 |
| **SLF-III** | 4,000 | 3,321 | 2,223 |
| **Vertical occipital fasciculus** | 6,000 | 5,963 | 70 |
| **Uncinate fasciculus** | 6,000 | 613 | 843 |
| **Medial lemniscus** | 6,000 | 5,955 | 5,948 |
| **Optic radiation** | 6,000 | 5,529 | 891 |
| **Optic radiation (whole occipital lobe)** | 8,000 | 7,434 | 7,625 |
| **Optic tract** | 2,000 | 1,956 | 1,982 |
| **Whole pyramidal tract** | 20,000 | 19,337 | 19,324 |
| **Cortico-spinal tract** | 15,000 | 14,711 | 14,733 |
| **M1 cortico-spinal tract** | 12,000 | 11,736 | 11,781 |
| **Supplementary motor area (SMA) pyramidal tract** | 6,000 | 5,743 | 5,841 |
| **Premotor cortex (PMC) pyramidal tract** | 4,000 | 13 | 7 |
| **Anterior thalamic radiation** | 6,000 | 4,151 | 1,908 |
| **Superior thalamic radiation** | 8,000 | 7,042 | 6,348 |
| **Parietal thalamic radiation** | 6,000 | 431 | 258 |
| **Dentato-rubro thalamic tract** | 2,000/50,000 | 193/2223 | 29/383 |
| **Inferior cerebellar peduncle** | 2,000 | 1,337 | 899 |
| **Middle cerebellar peduncle** | 8,000 | 2,800 | 3,227 |
| **Anterior commissure** | 6,000 | 307 | |
| **Prefrontal corpus callosum (CC)** | 10,000 | 3,943 | |
| **PMC and SMA CC** | 10,000 | 1,767 | |

| | | |
|---|---|---|
| **Motor corpus CC** | 10,000 | 2,567 |
| **Sensory CC** | 10,000 | 2,828 |
| **Parietal CC** | 10,000 | 5,772 |
| **Occipital CC** | 10,000 | 3,457 |
| **Temporal CC** | 10,000 | 85 |

### Supplementary results

#### FWT dissection protocol

Supplementary table 2 includes a list of all VOIs used to generate the includes and excludes for each bundle. FS VOIs are denoted by the prefix FS, MSBP VOIs are denoted by the prefix MSBP, UKBB derived VOIs are denoted by the prefix UKBB, NIST PD25 VOIs are denoted by the prefix PD25, Jülich Histological atlas derived VOIs are denoted by JHA, and custom made VOIs are denoted by the prefix custom. Grey matter and white matter VOIs are denoted by the suffix GM and WM respectively.

**Supplementary table 2: FWT bundle inclusion and exclusion VOIs (L&R) denote left and right paired bundles, (B) denotes commissural (bilateral) bundles**

| No. | Bundles | Inclusion VOIs | Exclusion VOIs |
|---|---|---|---|
| 1 | **Arcuate fasciculus (L&R)** | 1- FS inferior frontal gyrus pars triangularis + FS inferior frontal gyrus pars opercularis<br>2- MSBP superior temporal gyrus 1 + MSBP superior temporal gyrus 2 + MSBP superior temporal gyrus 3 + FS bank of superior temporal sulcus GM + FS supramarginal GM | FS all corpus callosum + FS brainstem + FS M1 GM + custom FS derived dilated putamen + FS thalamus + MSBP superior temporal gyrus 4 + MSBP superior temporal gyrus 5 + FS parahippocampal GM + FS parahippocampal WM + custom FS derived Insula GM+WM eroded + FS cingulate lobe GM + FS cingulate lobe WM + FS superior frontal GM + FS superior frontal WM |

| | | | |
|---|---|---|---|
| 2 | **Arcuate fasciculus with precentral gyrus (L&R)** | 1- FS inferior frontal gyrus pars triangularis + FS inferior frontal gyrus pars opercularis + MSBP precentral 1 + MSBP precentral 2 + MSBP precentral 3 | FS all corpus callosum + FS brainstem + + MSBP precentral 4 + MSBP precentral 5 + MSBP precentral 6 + custom FS derived dilated putamen + FS thalamus + MSBP superior temporal gyrus 4 + MSBP superior temporal gyrus 5 + FS parahippocampal GM + FS parahippocampal WM + custom FS derived Insula GM+WM eroded + FS cingulate lobe GM + FS cingulate lobe WM + FS superior frontal GM + FS superior frontal WM |
| | | 2- MSBP superior temporal gyrus 1 + MSBP superior temporal gyrus 2 + MSBP superior temporal gyrus 3 + FS bank of superior temporal sulcus GM + FS supramarginal GM | |
| 3 | **Cingulate cingulum bundle (L&R)** | 1- FS rostral anterior cingulate GM + FS caudal anterior cingulate GM | FS all corpus callosum + FS brainstem + FS superior temporal gyrus GM + FS unsegmented WM + FS ventral diencephalon + FS inferior frontal gyrus pars opercularis GM + FS inferior frontal gyrus pars opercularis GM + FS inferior frontal gyrus pars triangularis GM + FS inferior frontal gyrus pars triangularis WM + FS parahippocampal GM + FS parahippocampal WM + FS superior frontal gyrus GM + FS Hippocampus + FS temporal lobe GM + FS temporal lobe WM + custom anterior commissure VOI + FS insula GM + FS insula WM |
| | | 2- FS posterior cingulate GM + FS isthmic posterior cingulate GM + FS precuneus GM | |
| 4 | **Temporal/ Parahippocampal cingulum bundle (L&R)** | 1- FS Hippocampus | FS all corpus callosum + FS brainstem + FS posterior cingulate GM + FS posterior cingulate WM + FS fusiform gyrus GM + FS fusiform gyrus WM + FS unsegmented WM + FS insula GM + FS insula WM + FS medial orbitofrontal GM + FS medial orbitofrontal WM + FS lateral orbitofrontal GM + FS lateral orbitofrontal WM + FS Amygdala |
| | | 2- FS precuneus GM + FS isthmic posterior cingulate GM | |
| 5 | **Fornix (L&R)** | 1- FS Fornix | FS all corpus callosum + FS brainstem + FS Putamen + FS isthmic posterior |

| | | | |
|---|---|---|---|
| | | 2- FS Hippocampus | cingulate GM + FS isthmic posterior cingulate WM + custom FS derived Thalamus eroded x2 + FS optic chiasm + FS frontal lobe GM + FS occipital lobe GM + FS occipital lobe WM + FS temporal lobe GM + FS temporal lobe WM + FS parietal lobe GM + FS parietal lobe WM + FS Amygdala + FS rostral anterior cingulate GM + FS choroid plexus + FS choroid plexus + custom FS derived anterior limb of internal capsule + FS Caudate + FS ventral diencephalon + FS Insula GM + FS Insula WM |
| **6** | **Frontal Aslant tract (L&R)** | 1- FS inferior frontal gyrus pars triangularis + FS inferior frontal gyrus pars opercularis | FS all corpus callosum + FS brainstem + FS Insula GM + FS Insula WM + FS M1 GM + FS M1 WM + FS medial orbitofrontal GM + FS medial orbitofrontal WM + FS lateral orbitofrontal GM + FS lateral orbitofrontal WM + FS inferior frontal gyrus pars orbitalis GM + FS inferior frontal gyrus pars orbitalis WM + FS caudate + MSBP superior frontal gyrus 1 + MSBP superior frontal gyrus 2 + MSBP superior frontal gyrus 3 + MSBP superior frontal gyrus 4 |
| | | 2- MSBP superior frontal gyrus 5 + MSBP superior frontal gyrus 6 | |
| **7** | **Inferior fronto-occipital fasciculus (L&R)** | 1- FS superior parietal lobule GM + FS inferior parietal lobule GM FS occipital lobe GM + FS precuneus GM + MSBP fusiform gyrus 1 + MSBP fusiform gyrus 2 | FS all corpus callosum + FS brainstem + MSBP superior temporal gyrus 5 + custom FS derived putamen eroded + FS parahippocampal gyrus GM + FS parahippocampaly gyrus WM + FS cingulate lobe GM + FS cingulate lobe WM + FS superior frontal gyrus GM + FS superior frontal gyrus WM + FS M1 GM + FS M1 WM + FS temporal pole GM + FS temporal pole WM + FS inferior temporal gyrus GM + custom anterior commissure midline VOI + FS accumbens + FS pallidum + FS amygdala + FS thalamus + FS hippocampus + FS ventral diencephalon + FS optic chiasm + custom FS derived posterior limb of internal capsule + custom FS derived anterior limb of internal capsule + custom FS derived anterior superior |
| | | 2- MSBP rostral midde frontal gyrus 5 + MSBP rostral middle frontal gyrus 6 + FS frontal pole + FS inferior frontal gyrus pars triangularis GM + FS medial orbitofrontal gyrus GM + FS lateral orbitofrontal gyrus GM | |

| | | | |
|---|---|---|---|
| | | | centrum semiovale (medial and lateral superior subsegments of FS unsegmented WM label) |
| **8** | **Inferior longitudinal fasciculus (L&R)** | 1- FS occipital lobe GM | FS all corpus callosum + FS brainstem + FS superior temporal gyrus GM + FS aparc2009 superior occipital gyrus GM + FS parahippocampal gyrus GM + FS parahippocampal gyrus WM + FS cingulate lobe GM + FS cingulate lobe WM + custom anterior commissure midline VOI + FS hippocampus + FS ventral diencephalon + FS optic chiasm + FS insula GM + FS insula WM + FS frontal lobe GM + FS frontal lobe WM + FS parietal lobe GM + FS parietal lobe WM + custom FS derived superior medial subcomponent of FS unsegmented WM label + FS M1 GM + FS M1 WM |
| | | 2- MSBP inferior temporal gyrus 1 + MSBP inferior temporal gyrus 2 + MSBP middle temporal gyrus 4 + FS temporal pole + MSBP fusiform gyrus 4 | |
| **9** | **Middle longitudinal fasciculus (L&R)** | 1- FS superior parietal lobule GM + FS inferior parietal lobule GM + FS aparc2009 superior occipital gyrus + FS cuneus GM + FS precuneus GM | FS all corpus callosum + FS brainstem + FS parahippocampal gyrus GM + FS parahippocampal gyrus WM + custom FS insular WM derived VOI (superior and anterior parts of insular WM label) + custom FS derived putamen finefinedilated + FS cingulate lobe GM + FS cingulate lobe WM + custom anterior commissure midline VOI + FS hippocampus + FS ventral diencephalon + FS optic chiasm + FS insula GM + FS insula WM + FS frontal lobe GM + FS frontal lobe WM + custom superior lateral subcomponent of FS unsegmented WM label + FS fusiform gyrus GM + FS fusiform gyrus WM + FS inferior temporal GM + FS inferior temporal WM |
| | | 2- MSBP superior temporal gyrus 5 + MSBP superior temporal gyrus 4 + MSBP superior temporal gyrus 3 + FS temporal pole GM | |
| **10** | **Whole Superior longitudinal fasciculus (L&R)** | 1- MSBP superior frontal gyrus 6 + 7 + 8 + FS caudal middle frontal gyrus + FS rostral middle frontal gyrus + FS inferior frontal gyrus pars triangularis GM + pars opercularis GM + pars orbitalis GM | MSBP superior frontal gyrus 1 + MSBPfine superior frontal gyrus 2 + MSBP superior frontal gyrus 3 + MSBP superior frontal gyrus 4 + MSBP superior frontal gyrus 5 + FS all corpus callosum + FS brainstem + custom FS derived putamen dilated + FS Thalamus + FS temporal lobe GM + FS temporal lobe WM + FS |

| | | | |
|---|---|---|---|
| | | 2- FS superior parietal lobule GM + FS inferior parietal lobule GM + FS supramarginal gyrus GM | parahippocampal GM + FS parahippocampal WM + FS insula GM + FS insula WM + FS cingulate lobe GM + FS cingulate lobe WM + FS posterior cingulate cortex GM + FS posterior cingulate cortex WM |
| **11** | **Superior longitudinal fasciculus 1 (L&R)** | 1- MSBP superior frontal gyrus 6 + MSBP superior frontal gyrus 7 + MSBP superior frontal gyrus 8 | FS all corpus callosum + FS brainstem + FS Insula GM + FS Insula WM + custom FS derived putamen dilated + FS thalamus + FS temporal lobe GM + FS temporal lobe WM + FS parahippocampal GM + FS parahippocampal WM + FS cingulate lobe GM + FS cingulate lobe WM + FS posterior cingulate GM + FS posterior cingulate WM + MSBP superior frontal gyrus 1 + MSBP superior frontal gyrus 2 + MSBP superior frontal gyrus 3 + MSBP superior frontal gyrus 4 + MSBP superior frontal gyrus 5 + FS S1 GM + FS S1 WM + FS inferior frontal gyrus pars triangularis GM + FS inferior frontal gyrus pars opercularis GM + FS inferior frontal gyrus pars orbitalis GM + FS inferior parietal lobule GM + FS supramarginal GM + FS M1 GM + MSBP superior parietal lobule 3 + MSBP superior parietal lobule 4 + MSBP superior parietal lobule 5 + MSBP superior parietal lobule 6 |
| | | 2- FS superior parietal lobule GM | |
| **12** | **Superior longitudinal fasciculus 2 dorsal (L&R)** | 1- FS caudal middle frontal GM + MSBP rostral middle frontal 1 | FS all corpus callosum + FS brainstem + FS Insula GM + FS Insula WM + custom FS derived putamen dilated + FS thalamus + FS temporal lobe GM + FS temporal lobe WM + FS rostral middle frontal GM + FS parahippocampal GM + FS parahippocampal WM + FS cingulate lobe GM + FS cingulate lobe WM + FS posterior cingulate GM + FS posterior cingulate WM + MSBP superior frontal gyrus 1 + MSBP superior frontal gyrus 2 + MSBP superior frontal gyrus 3 + MSBP superior frontal gyrus 4 + MSBP superior frontal gyrus 5 + MSBP superior frontal gyrus 6 + MSBP superior frontal gyrus 7 + MSBP |
| | | 2- FS supramarginal GM + FS inferior parietal lobule GM | |

| | | | |
|---|---|---|---|
| | | | superior frontal gyrus 8 + FS superior parietal lobule GM |
| 13 | **Superior longitudinal fasciculus 2 ventral (L&R)** | 1- FS rostral middle frontal GM | FS all corpus callosum + FS brainstem + FS Insula GM + FS Insula WM + custom FS derived putamen dilated + FS thalamus + FS temporal lobe GM + FS temporal lobe WM + FS caudal middle frontal GM + FS parahippocampal GM + FS parahippocampal WM + FS cingulate lobe GM + FS cingulate lobe WM + FS posterior cingulate GM + FS posterior cingulate WM + MSBP superior frontal gyrus 1 + MSBP superior frontal gyrus 2 + MSBP superior frontal gyrus 3 + MSBP superior frontal gyrus 4 + MSBP superior frontal gyrus 5 + MSBP superior frontal gyrus 6 + MSBP superior frontal gyrus 7 + MSBP superior frontal gyrus 8 + FS superior parietal lobule GM |
| | | 2- FS supramarginal GM + FS inferior parietal lobule GM | |
| 14 | **Superior longitudinal fasciculus 3 (L&R)** | 1- FS inferior frontal gyrus pars triangularis GM + pars opercularis GM + pars orbitalis GM | FS all corpus callosum + FS brainstem + FS Insula GM + FS Insula WM + custom FS derived putamen dilated + FS thalamus + FS temporal lobe GM + FS temporal lobe WM + FS caudal middle frontal GM + MSBP rostral middle frontal gyrus 1 + FS parahippocampal GM + FS parahippocampal WM + FS cingulate lobe GM + FS cingulate lobe WM + FS posterior cingulate GM + FS posterior cingulate WM + MSBP superior frontal gyrus 1 + MSBP superior frontal gyrus 2 + MSBP superior frontal gyrus 3 + MSBP superior frontal gyrus 4 + MSBP superior frontal gyrus 5 + MSBP superior frontal gyrus 6 + MSBP superior frontal gyrus 7 + MSBP superior frontal gyrus 8 + FS superior parietal lobule GM |
| | | 2- FS supramarginal GM + FS inferior parietal lobule GM | |
| 15 | **Uncinate fasciculus (L&R)** | 1- MSBP middle temporal gyrus 4 + MSBP inferior temporal gyrus 1 + FS temporal | FS all corpus callosum + FS brainstem + FS Accumbens + FS Thalamus + FS caudate + FS Amygdala + FS Hippocampus + FS unsegmented WM |

| | | | |
|---|---|---|---|
| 16 | | pole GM + MSBP entorhinal cortex | + FS parietal lobe GM + FS parietal lobe WM + FS occipital lobe GM + FS occipital lobe WM + FS rostral anterior cingulate cortex GM + FS rostral anterior cingulate cortex WM + FS inferior frontal gyrus pars opercularis GM + FS inferior frontal gyrus pars opercularis WM + FS M1 GM + FS M1 WM + MSBP insula 1 + MSBP insula 2 + MSBP insula 3 + custom FS derived insula WM exclude (including superior aspect, and posterior aspects of insular WM) + FS medial orbitofrontal gyrus GM |
| | | 2- FS inferior frontal gyrus pars triangularis GM + FS inferior frontal gyrus pars orbitalis GM + FS frontal pole GM + FS lateral orbitofrontal GM | |
| 16 | **Vertical occipital fasciculus (L&R)** | 1- MSBP cuneus 2 + MSBP superior parietal lobule 2 + MSBP inferior parietal lobule 3 | FS all corpus callosum + FS brainstem + FS parahippocampal gyrus GM+WM + FS frontal lobe GM + FS frontal lobe WM + FS superior temporal gyrus GM + FS superior temporal gyrus WM + FS precuneus GM + MSBP inferior temporal gyrus 1 + 2 + 3 + FS middle temporal gyrus + FS S1 GM + MSBP fusiform gyrus 3 + custom MSBP derived fusiform gyrus 4 GM+WM |
| | | 2- MSBP lateral occipital gyrus 5 + MSBP lateral occipital gyrus 5 + MSBP fusiform gyrus 1 | |
| 17 | **Medial Lemniscus (L&R)** | 1- Custom brainstem ML VOI (UKBB derived) | FS all corpus callosum + custom contralateral half brainstem + FS cingulate lobe GM + FS cingulate lobe WM + custom eroded CST VOI (UKBB derived) + custom MSBP derived bilateral hypothalamus dilated + FS temporal lobe GM + FS temporal lobe WM + FS putamen + FS caudal middle frontal GM + FS caudate + FS hippocampus |
| | | 2- PD25 VPL and VPM | |
| | | 3- FS S1 GM | |
| 18 | **Optic radiation (classic) (L&R)** | 1- PD25 Pulvinar + Jülich lateral geniculate nucleus | FS all corpus callosum + FS brainstem + FS parahippocampal gyrus GM + FS ventral diencephalon + custom fs insular WM derived VOI (superior and anterior parts of insular WM label) + PD25 ventral-anterior and ventral-lateral thalamic VOIs + FS cingulate lobe GM + FS cingulate lobe WM + custom anterior commissure midline VOI + FS hippocampus + FS ventral diencephalon + FS optic chiasm + FS insula GM + custom PLIC VOI + FS |
| | | 2- FS aparc2009 calcarine sulcus + FS pericalcarine cortex GM | |

| | | | |
|---|---|---|---|
| | | | frontal lobe GM + FS parietal lobe GM + FS isthmic posterior cingulate cortex GM + FS caudate + custom ALIC VOI + custom superior lateral subcomponent of FS unsegmented WM label + custom superior medial subcomponent of FS unsegmented WM label + custom inferior medial subcomponent of FS unsegmented WM label + custom FS derived mid-segment of insular WM + MSBP fusiform gyrus 4 + FS temporal pole GM + FS temporal pole WM + FS inferior temporal gyrus GM + FS inferior temporal WM |
| 19 | **Optic radiation (whole occipital lobe) (L&R)** | 1- FS occipital lobe GM | FS all corpus callosum + FS brainstem + FS parahippocampal gyrus GM + FS ventral diencephalon + custom fs insular WM derived VOI (superior and anterior parts of insular WM label) + PD25 ventral-anterior and ventral-lateral thalamic VOIs + FS cingulate lobe GM + FS cingulate lobe WM + custom anterior commissure midline VOI + FS hippocampus + FS optic chiasm + FS insula GM + custom PLIC VOI + FS frontal lobe GM + FS parietal lobe GM + FS isthmic posterior cingulate cortex GM + FS caudate + custom FS derived ALIC VOI + custom superior lateral subcomponent of FS unsegmented WM label + custom superior medial subcomponent of FS unsegmented WM label + custom inferior medial subcomponent of FS unsegmented WM label + custom FS derived mid-segment of insular WM + MSBP fusiform gyrus 4 + FS temporal pole GM + FS temporal pole WM + FS inferior temporal gyrus GM + FS inferior temporal WM |
| | | 2- PD25 Pulvinar + Jülich lateral geniculate nucleus 68 | |
| 20 | **Optic tract (L&R)** | 1- FS optic chiasm | FS all corpus callosum + FS brainstem + FS putamen + FS hippocampus + FS caudate + FS aseg derived cortical grey matter + FS frontal lobe WM + FS fornix + PD25 dorsomedial thalamic VOI + PD25 dorsolateral thalamic VOI |
| | | 2- PD25 Pulvinar | |

| | | | |
|---|---|---|---|
| | | | + FS parietal lobe GM + FS parietal lobe WM + FS M1 GM + FS M1 WM |
| **21** | **Whole Pyramidal tract (L&R)** | 1- FS Brainstem<br>2- FS M1 GM + FS S1 GM + FS paracingulate GM + MSBP superior frontal gyrus 5 + MSBP superior frontal gyrus 6 + MSBP superior frontal gyrus 7 + MSBP superior frontal gyrus 8 + FS caudal middle frontal gyrus | FS all corpus callosum + custom contralateral half brainstem + FS cingulate lobe GM + FS cingulate lobe WM + custom MSBP derived bilateral hypothalamus dilated + custom FS derived ipsilateral thalamus eroded + FS temporal lobe GM + FS temporal lobe WM + FS Caudate + FS putamen + FS rostral middle frontal gyrus GM + FS superior parietal lobule GM + FS occipital lobe GM + FS occipital lobe WM |
| **22** | **CorticoSpinal tract (L&R)** | 1- Custom brainstem CST VOI (UKBB derived)<br>2- FS M1 GM + FS S1 GM + FS paracingulate GM | FS all corpus callosum + custom contralateral half brainstem + FS cingulate lobe GM + FS cingulate lobe WM + custom MSBP derived bilateral hypothalamus dilated + custom FS derived ipsilateral thalamus eroded + FS temporal lobe GM + FS temporal lobe WM + FS Caudate + FS putamen + FS superior frontal gyrus GM + FS occipital lobe GM + FS occipital lobe WM |
| **23** | **M1 corticospinal tract (L&R)** | 1- Custom brainstem CST VOI (UKBB derived)<br>2- FS M1 GM + FS paracingulate GM | FS all corpus callosum + custom contralateral half brainstem + FS cingulate lobe GM + FS cingulate lobe WM + custom MSBP derived bilateral hypothalamus dilated + custom FS derived ipsilateral thalamus eroded + FS temporal lobe GM + FS temporal lobe WM + FS Caudate + FS putamen + FS superior frontal gyrus GM + FS occipital lobe GM + FS occipital lobe WM |
| **24** | **Premotor Pyramidal tract (L&R)** | 1- FS Brainstem<br>2- FS caudal middle frontal gyrus | FS all corpus callosum + custom contralateral half brainstem + FS cingulate lobe GM + FS cingulate lobe WM + custom eroded ML VOI (UKBB derived) + custom MSBP derived bilateral hypothalamus dilated + custom FS derived ipsilateral Thalamus eroded + FS temporal lobe |

| | | | |
|---|---|---|---|
| | | | GM + FS temporal lobe WM + FS putamen + FS superior frontal gyrus GM + FS superior frontal gyrus WM + FS caudate + FS M1 GM + FS M1 WM + FS S1 GM + FS S1 WM + FS rostral middle frontal gyrus + FS occipital lobe GM + FS occipital lobe WM |
| 25 | **Supplementary Motor Pyramidal tract (L&R)** | 1- FS Brainstem | FS all corpus callosum + custom contralateral half brainstem + FS cingulate lobe GM + FS cingulate lobe WM + custom eroded ML VOI (UKBB derived) + custom MSBP derived bilateral hypothalamus dilated + custom FS derived ipsilateral Thalamus eroded + FS temporal lobe GM + FS temporal lobe WM + FS putamen + FS caudal middle frontal GM + FS rostral middle frontal GM + FS caudate + FS M1 GM + FS M1 WM + FS S1 GM + FS S1 WM + FS occipital lobe GM + FS occipital lobe WM + MSBP superior frontal gyrus 1 + MSBP superior frontal gyrus 2 + MSBP superior frontal gyrus 3 + MSBP superior frontal gyrus 4 |
| | | 2- MSBP superior frontal gyrus 5 + MSBP superior frontal gyrus 6 + MSBP superior frontal gyrus 7 + MSBP superior frontal gyrus 8 | |
| 26 | **Superior thalamic radiation (L&R)** | 1- FS Thalamus | FS all corpus callosum + FS brainstem + custom anterior commissure VOI + custom MSBP derived bilateral hypothalamus dilated + FS temporal lobe GM + FS Putamen + FS Caudate + PD25 Pulvinar VOI + custom FS derived Caudate eroded + MSBP superior frontal gyrus 1 + MSBP superior frontal gyrus 2 + MSBP superior frontal gyrus 3 + MSBP superior frontal gyrus 4 + FS caudal anterior cingulate cortex GM + PD25 Dorsomedial thalamic nucleus VOI + PD25 Dorsolateral thalamic nucleus + FS Fornix + FS parietal lobe GM + FS parietal lobe WM + FS rostral middle frontal gyrus GM + FS rostral middle frontal gyrus WM |
| | | 2- FS M1 GM + FS paracentral lobule GM + MSBP superior frontal gyrus 5 + MSBP superior frontal gyrus 6 + MSBP superior frontal gyrus 7 + MSBP superior frontal gyrus 8 + FS caudal middle frontal gyrus | |
| 27 | **Anterior thalamic radiation** | 1- FS Thalamus | FS all corpus callosum + FS brainstem + custom anterior commissure VOI + |

| | | | |
|---|---|---|---|
| | **(L&R)** | 2- MSBP superior frontal gyrus 1 + MSBP superior frontal gyrus 2 + MSBP superior frontal gyrus 3 + MSBP superior frontal gyrus 4 + FS medial orbitofrontal gyrus GM + FS lateral orbitofrontal gyrus WM + FS frontal pole GM | custom MSBP derived bilateral hypothalamus dilated + FS temporal lobe GM + FS Putamen + FS Caudate + PD25 Pulvinar + FS Fornix + FS caudal anterior cingulate cortex GM + FS occipital lobe GM + FS parietal lobe GM + PD25 Ventroposterior+anterolateral+postero lateral+posteromedial thalamic nuclei + FS M1 GM + FS M1 WM + MSBP superior frontal gyrus 5 + FS caudal middle frontal gyrus GM + FS parahippocampal gyrus GM + FS parahippocampal gyrus WM + FS Insula GM + FS Insula WM + FS cingulate lobe GM + FS cingulate lobe WM + FS ventral diencephalon |
| **28** | **Parietal thalamic radiation (L&R)** | 1- FS Thalamus | FS all corpus callosum + FS brainstem + FS ventral diencephalon + custom anterior commissure VOI + custom FS derived anterior limb of internal capsule + FS temporal lobe GM + FS Putamen + FS frontal lobe GM + FS Fornix + FS occipital lobe GM + FS Caudate + PD25 Pulvinar + custom FS derived caudate eroded + MSBP superior frontal gyrus 1 + MSBP superior frontal gyrus 3 + MSBP superior frontal gyrus 4 + FS caudal anterior cingulate cortex GM + PD25 dorsomedial thalamic nucleus + PD25 dorsolateral + custom FS derived medial inferior unsegmented WM |
| | | 2- FS parietal lobe GM | |
| **29** | **Anterior commissure (B)** | 1- Custom anterior commissure VOI | FS all corpus callosum + custom posterior commissure + FS brainstem + FS optic chiasm + FS Thalamus left + FS Thalamus right" |
| | | 2- FS temporal lobe GM left + FS occipital lobe GM left | |
| | | 3- FS temporal lobe GM right + FS occipital lobe GM right | |
| **30** | **Prefrontal corpus callosum (B)** | 1- FS corpus callosum Genu (anterior) + FS corpus callosum anterior (mid-anterior) | custom anterior commissure VOI + custom posterior commissure VOI + FS CC central + FS CC isthmus (midposterior) + FS CC splenium |

| | | | |
|---|---|---|---|
| 31 | | 2- FS inferior frontal gyrus pars triangularis GM left + FS inferior frontal gyrus pars orbitalis GM left + FS frontal pole GM left + FS lateral orbitofrontal GM left + FS medial orbitofrontal GM left + FS caudal middle frontal gyrus GM left + FS rostral middle frontal gyrus GM left + FS superior frontal gyrus GM left | (posterior) + FS fornix + FS caudal middle frontal gyrus left + FS caudal middle frontal gyrus right + FS M1 GM left + FS M1 GM right + FS parietal lobe GM left + FS parietal lobe GM right + FS brainstem + FS cingulate lobe GM left + FS cingulate lobe GM right + FS insula lobe GM left + FS insula lobe WM left + FS insula lobe GM right + FS insula lobe WM right + custom FS derived anterior limb of internal capsule left & right + ventral diencephalon left & right |
| | | 3- FS inferior frontal gyrus pars triangularis GM right + FS inferior frontal gyrus pars orbitalis GM right + FS frontal pole GM right + FS lateral orbitofrontal GM right + FS medial orbitofrontal GM right + FS caudal middle frontal gyrus GM right + FS rostral middle frontal gyrus GM right + FS superior frontal gyrus GM right | |
| 31 | **Premotor and supplementary motor corpus callosum (B)** | 1- FS corpus callosum central | custom anterior commissure VOI + custom posterior commissure VOI + FS CC genu (anterior) + FS CC isthmus (midposterior) + FS CC splenium (posterior) + FS fornix + FS brainstem + FS cingulate lobe GM left + FS cingulate lobe GM right +custom FS derived anterior limb of internal capsule left & right + ventral diencephalon left & right |
| | | 2- FS caudal middle frontal gyrus GM left + MSBP superior frontal gyrus 5 left + MSBP superior frontal gyrus 6 left + MSBP superior frontal gyrus 7 left + MSBP superior frontal gyrus 8 left | |
| | | 3- FS caudal middle frontal gyrus GM right + MSBP superior frontal gyrus 5 right + MSBP | |

| | | | |
|---|---|---|---|
| | | superior frontal gyrus 6 right + MSBP superior frontal gyrus 7 right + MSBP superior frontal gyrus 8 right | |
| **32** | **Motor corpus callosum (B)** | 1- FS corpus callosum isthmus (midposterior)<br>2- FS M1 GM left<br>3- FS M1 GM right | custom anterior commissure VOI + custom posterior commissure VOI + FS CC genu (anterior) + FS CC anterior (midanterior) + FS CC central + FS CC splenium (posterior) + FS fornix + FS caudal middle frontal gyrus GM left + FS caudal middle frontal gyrus GM right + FS superior frontal gyrus GM left + FS superior frontal gyrus GM right + FS parietal lobe GM left + FS parietal lobe GM right + FS brainstem + FS cingulate lobe GM left + FS cingulate lobe GM right |
| **33** | **Sensory corpus callosum (B)** | 1- FS corpus callosum isthmus (midposterior)<br>2- FS S1 GM left<br>3- FS S1 GM right | custom anterior commissure VOI + custom posterior commissure VOI + FS CC genu (anterior) + FS CC anterior (midanterior) + FS CC central + FS CC splenium (posterior) + FS fornix + FS caudal middle frontal gyrus GM left + FS caudal middle frontal gyrus GM right + FS superior frontal gyrus GM left + FS superior frontal gyrus GM right + FS M1 GM + FS M1 WM + FS brainstem + FS cingulate lobe GM left + FS cingulate lobe GM right |
| **34** | **Parietal corpus callosum (B)** | 1- FS corpus callosum splenium (posterior)<br>2- FS parietal lobe GM left<br>3- FS parietal lobe GM right | custom anterior commissure VOI + custom posterior commissure VOI + FS CC genu (anterior) + FS CC anterior (midanterior) + FS CC central + FS CC splenium (posterior) + FS fornix + FS caudal middle frontal gyrus GM left + FS caudal middle frontal gyrus GM right + FS superior frontal gyrus GM left + FS superior frontal gyrus GM right + FS S1 GM left + FS S1 GM right + FS brainstem + FS cingulate lobe GM left + FS cingulate lobe GM right |

| | | | |
|---|---|---|---|
| 35 | **Occipital corpus callosum (B)** | 1- FS corpus callosum splenium (posterior)<br>2- FS occipital lobe GM left<br>3- FS occipital lobe GM right | custom anterior commissure VOI + custom posterior commissure VOI + FS CC genu (anterior) + FS CC anterior (midanterior) + FS CC central + FS CC isthmus (midposterior) + FS fornix + FS frontal lobe GM left + FS frontal lobe GM right + FS parietal lobe GM left + FS parietal lobe GM right + FS temporal lobe GM left + FS temporal lobe GM right + FS brainstem + FS cingulate lobe GM left + FS cingulate lobe GM right |
| 36 | **Temporal corpus callosum (B)** | 1- FS corpus callosum splenium (posterior)<br>2- FS inferior temporal gyrus GM left + FS middle temporal gyrus GM left + FS superior temporal gyrus GM left + FS temporal pole GM left<br>3- FS inferior temporal gyrus GM right + FS middle temporal gyrus GM right + FS superior temporal gyrus GM right + FS temporal pole GM right | custom anterior commissure VOI + custom posterior commissure VOI + FS CC genu (anterior) + FS CC anterior (midanterior) + FS CC central + FS CC isthmus (midposterior) + FS fornix + FS frontal lobe GM left + FS frontal lobe GM right + FS parietal lobe GM left + FS parietal lobe GM right + FS occipital lobe GM left + FS occipital lobe GM right + FS brainstem + FS cingulate lobe GM left + FS cingulate lobe GM right + FS Thalamus left + FS Thalamus right + FS parahippocampal GM left + FS parahippocampal GM right + FS hippocampus left + FS hippocampus right + FS amygdala left + FS amygdala right |
| 37 | **Inferior cerebellar peduncle (ICP) (L&R)** | 1- MSBP medulla<br>2- SUIT cerebellar Fastigial nucleus + SUIT cerebellar interposed nucleus + SUIT dentate nucleus | FS occipital lobe GM + FS occipital lobe WM + MSBP midbrain + ipsilateral FS ventral diencephalon + contralateral FS ventral diencephalon + MSBP pons + SUIT cerebellar X + SUIT cerebellar IX + SUIT cerebellar I_IV |
| 38 | **Dentato-rubro-thalamic tract (DRTT) (L&R)** | 1- SUIT cerebellar dentate nucleus<br>2- PD25 red nucleus (contralateral)<br>3- PD25 ventrolateral thalamic nucleus (contralateral) | FS corpus callosum + custom ALIC VOI + MSBP medulla + SUIT cerebellar IX |

| | | | |
|---|---|---|---|
| | | 4- FS M1 GM (contralateral) | |
| **39** | **Middle cerebellar peduncle (MCP) (L&R)** | 1- custom MSBP derived contralateral Pontine VOI | FS occipital lobe GM + FS occipital lobe WM + MSBP midbrain + MSBP medulla + FS ventral diencephalon + FS ventral diencephalon + SUIT cerebellar IX |
| | | 2- SUIT cerebellar VI + SUIT cerebellar I + SUIT cerebellar VIIb | |

#### Tractography results

The HCP-template bundles, voxel-wise heat-maps, additional screenshots are available as well as the symmetrical versions of the HCP-template bundles generated using the bundle-specific approach and with bundle segmentation from whole brain tractograms with 20 and 40 million streamlines are available via the link (https://osf.io/snq2d/). As previously mentioned, tractography results generated using the bundle-specific approach with 20,000 initial required streamlines were more reproducible. Additionally, the improvement in DRTT results were more noticeable when using the template bundles generated with 50,000 required streamlines initially. Supplementary figure 1 shows both version of the template DRTT bundles and plots for wDSC using both templates.

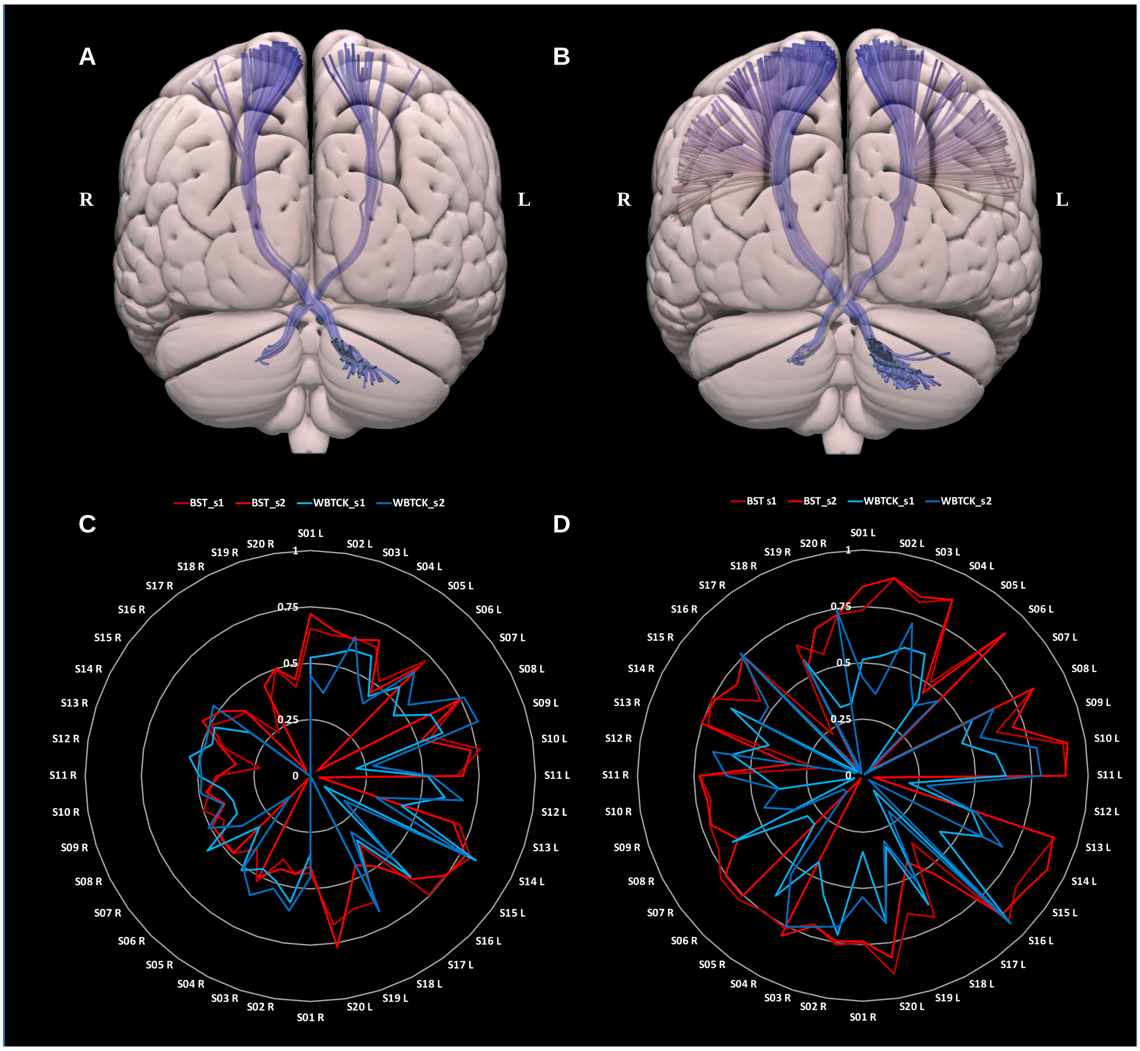


**Supplementary figure 1:** (A) and (B) show the template DRTTs generated using with 2000, and 50,000 streamlines during initial tractography (prior to streamlines filtering). (C) and (D) show the radar plots of wDSC (vertical range) for HCP test-retest output using BST with 20,000 streamlines initially, and using whole brain tractography and segmentation compared to the template bundles with 2000 streamlines (C) and to the template bundles with 50,000 streamlines (D). BST = Bundle specific tractography, WBTCK = whole brain tractography and segmentation, s1 = session 1, s2 = session 2, S = subject, L = left, R = right, wDSC = weighted dice similarity coefficient.

#### Reproducibility measures

**Bundle adjacency (BA)** is defined as the average of the coverage of bundle A by bundle B, and bundle B by bundle A (Garyfallidis et al., 2012):

$$BA(A,B) = \frac{\left(coverage(A,B) + coverage(A,B)\right)}{2}$$

The coverage of bundle A by bundle B is defined as the fraction of streamlines in A that is adjacent to B. A streamline in A is adjacent to B if there is at least one streamline in B for which MDF is lower than a set threshold. The minimum average direct-flip distance (MDF) between two streamlines is defined as the mean Euclidean distance between all corresponding points on the streamline (or the flipped version of a streamline whatever is the minimum).

**Dice similarity coefficient (DSC)** ranges between 0 (lowest) and 1 (highest), and can be defined as the ratio of voxel-wise intersection and union between 2 bundles (A and B) (Cousineau et al., 2017):

$$D(A,B) = \frac{|A \cap B|}{|A| + |B|}$$

in which | | is the number of voxels in the bundle for which at least 1 streamline is present.

**Density correlation (DC)** is defined as the Pearson correlation score between the streamline density maps of two bundles (Rheault et al., 2020). It ranges between 0 (lowest) and 1 (highest). The streamline density in a voxel is defined as the fraction of streamlines passing through that voxel.

**Weighted dice similarity coefficient** is a version of the DSC that takes streamline density into account. It ranges between 0 (lowest) and 1 (highest), and can be defined as the ratio between the sum of streamline density in all overlapping voxels and the sum of all voxels streamline density between two bundles (A and B) (Cousineau et al., 2017).

$$D(A,B) = \frac{\Sigma_{\mathrm{i \epsilon intersection}}\ (\mathrm{A + B})}{\Sigma_j\ \mathrm{A}_i}$$

where $A_i$ and $B_i$ are the streamline density in voxel i for bundle A respectively bundle B.

**Volume overlap** can be defined as the proportion of voxels in bundle A traversed by at least 1 streamline belonging to bundle B. It ranges between 0 (lowest) and 100% (highest) (Maier-Hein et al., 2017).

**Volume overreach** can be defined as the ratio between total number of voxels not belonging to bundle A but traversed by at least 1 streamline from bundle B, and total number of voxels in bundle A.

Supplementary figures 2 - 6 show violin plots for the model-based similarity measures bundle adjacency, Dice similarity coefficient, density correlation, volume overlap, and volume overreach. HCP test-retest results are used to create the violin plots and MASSIVE results are overlaid in black asterisks.

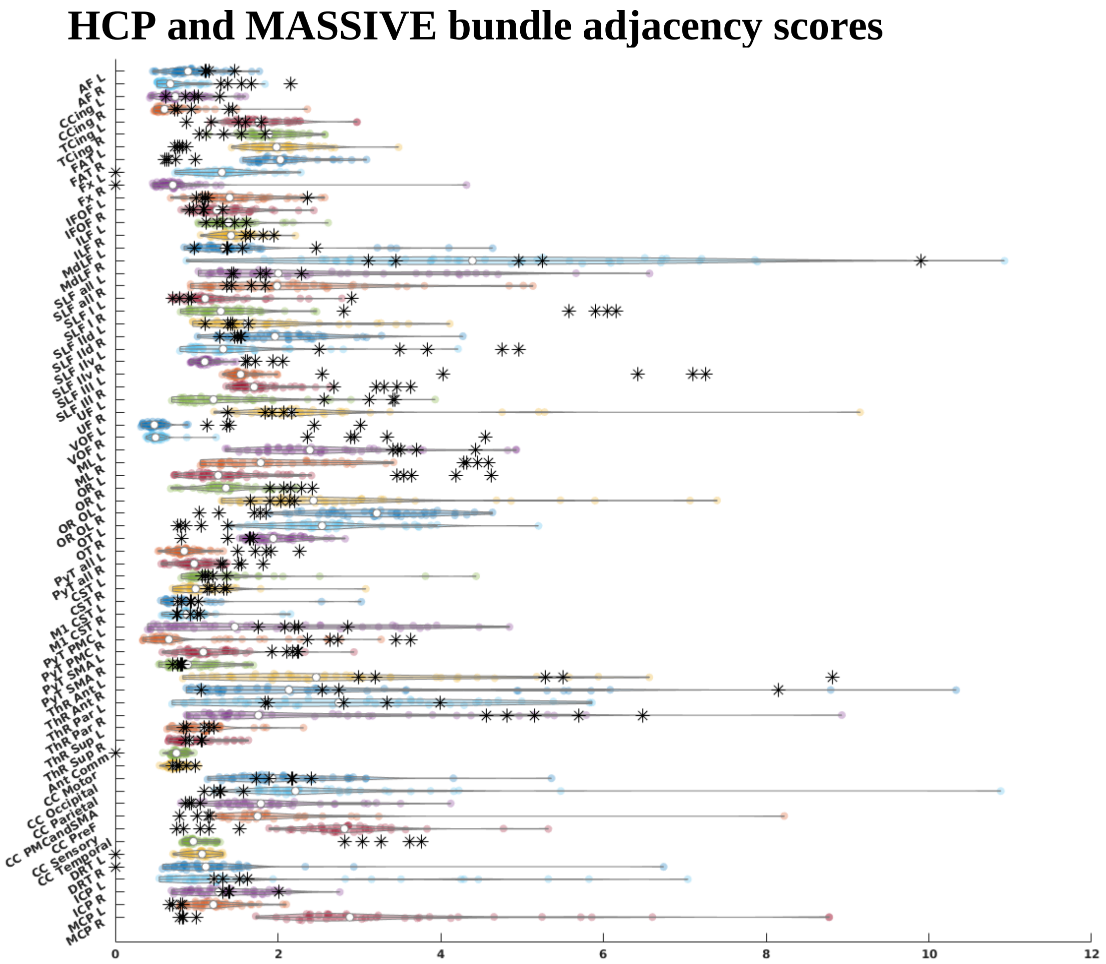


**Supplementary figure 2:** Bundle adjacency scores for all bundles, HCP test-retest tractograms are depicted as violin plots and MASSIVE results are depicted as black asterisks. AF = arcuate fasciculus, CCing = cingulate cingulum, TCing = temporal cingulum, FAT = frontal aslant tract, Fx = fornix, IFOF = inferior fronto-occipital fasciculu, ILF = inferior longitudinal fasciculus, MdLF = middle longitudinal fasciculus, SLF = superior longitudinal fasciculus, SLF-IId = SLF-II dorsal division, SLF-IIv = SLF-II ventral division, UF = uncinate fasciculus, VOF = vertical occipital fasciculus, ML = medial lemniscus, OR = optic radiation, occlobe = occipital lobe, OT = optic tract, PyT = pyramidal tract, CST = corticospinal tract, M1 = primary motor cortex, PyT = pyramidal tract, PMC = premotor cortex, SMA = supplementary motor area, ThR = thalamic radiation, Ant = anterior, Par = parietal, Sup = superior, Ant Comm = anterior commissure, CC = corpus callosum, PMC and SMA = premotor cortex and supplementary motor area, DRT = dentato-rubro-thalamic tract, ICP = inferior cerebellar peduncle, MCP = middle cerebellar peduncle, L = left, R = right.

### HCP and MASSIVE dice similarity coefficient scores

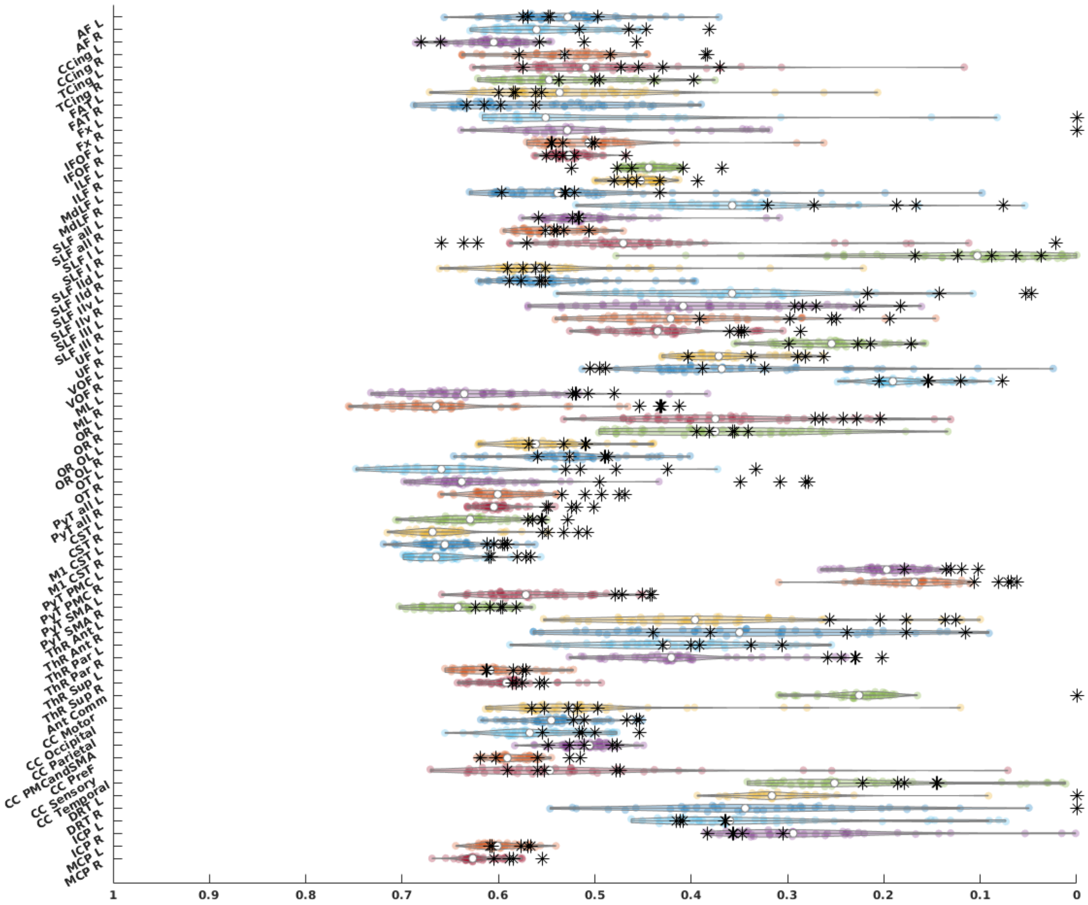


**Supplementary figure 3:** Dice similarity coefficient scores for all bundles, HCP test-retest tractograms are depicted as violin plots and MASSIVE results are depicted as black asterisks. AF = arcuate fasciculus, CCing = cingulate cingulum, TCing = temporal cingulum, FAT = frontal aslant tract, Fx = fornix, IFOF = inferior fronto-occipital fasciculu, ILF = inferior longitudinal fasciculus, MdLF = middle longitudinal fasciculus, SLF = superior longitudinal fasciculus, SLF-IId = SLF-II dorsal division, SLF-IIv = SLF-II ventral division, UF = uncinate fasciculus, VOF = vertical occipital fasciculus, ML = medial lemniscus, OR = optic radiation, occlobe = occipital lobe, OT = optic tract, PyT = pyramidal tract, CST = corticospinal tract, M1 = primary motor cortex, PyT = pyramidal tract, PMC = premotor cortex, SMA = supplementary motor area, ThR = thalamic radiation, Ant = anterior, Par = parietal, Sup = superior, Ant Comm = anterior commissure, CC = corpus callosum, PMC and SMA = premotor cortex and supplementary motor area, DRT = dentato-rubro-thalamic tract, ICP = inferior cerebellar peduncle, MCP = middle cerebellar peduncle, L = left, R = right.

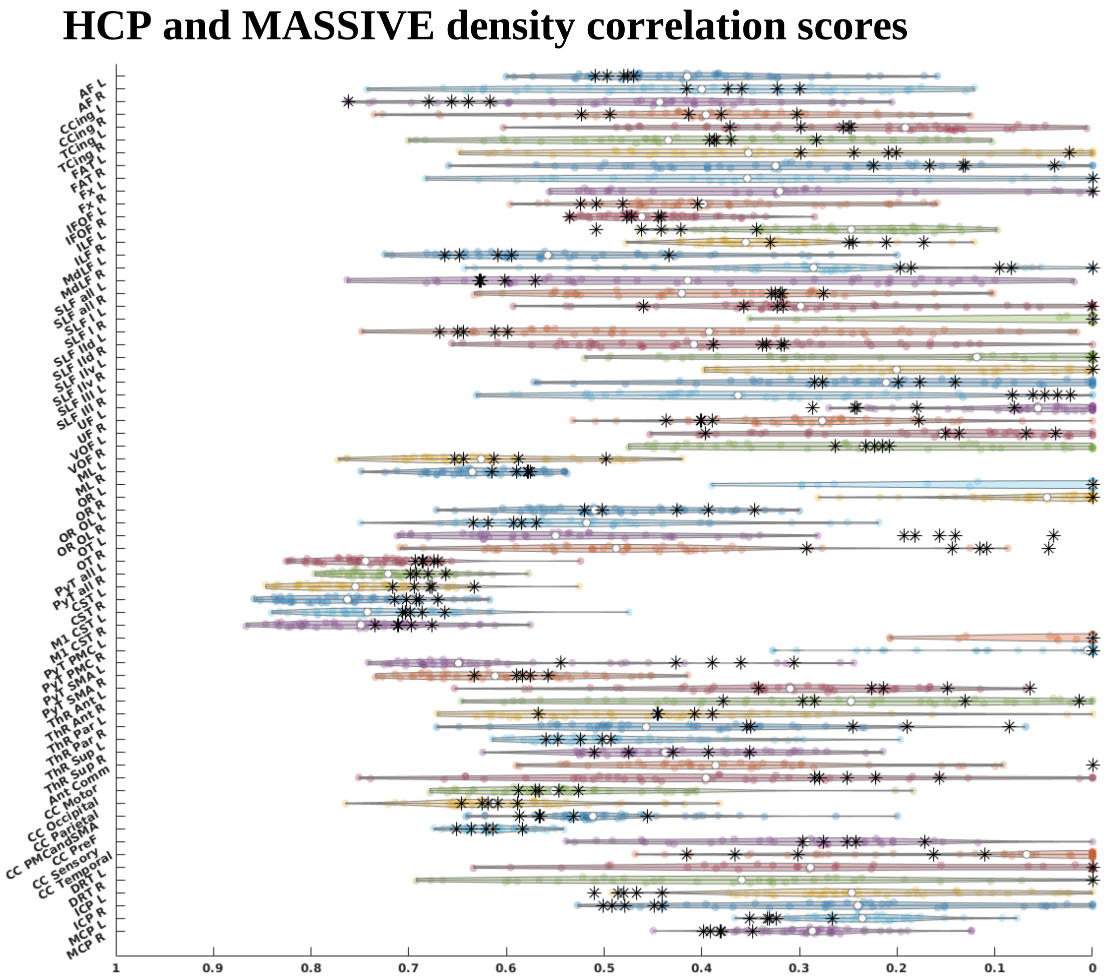


**Supplementary figure 4:** Density correlation scores for all bundles, HCP test-retest tractograms are depicted as violin plots and MASSIVE results are depicted as black asterisks. AF = arcuate fasciculus, CCing = cingulate cingulum, TCing = temporal cingulum, FAT = frontal aslant tract, Fx = fornix, IFOF = inferior fronto-occipital fasciculu, ILF = inferior longitudinal fasciculus, MdLF = middle longitudinal fasciculus, SLF = superior longitudinal fasciculus, SLF-IId = SLF-II dorsal division, SLF-IIv = SLF-II ventral division, UF = uncinate fasciculus, VOF = vertical occipital fasciculus, ML = medial lemniscus, OR = optic radiation, occlobe = occipital lobe, OT = optic tract, PyT = pyramidal tract, CST = corticospinal tract, M1 = primary motor cortex, PyT = pyramidal tract, PMC = premotor cortex, SMA = supplementary motor area, ThR = thalamic radiation, Ant = anterior, Par = parietal, Sup = superior, Ant Comm = anterior commissure, CC = corpus callosum, PMC and SMA = premotor cortex and supplementary motor area, DRT = dentato-rubro-thalamic tract, ICP = inferior cerebellar peduncle, MCP = middle cerebellar peduncle, L = left, R = right.

### HCP and MASSIVE volume overlap scores

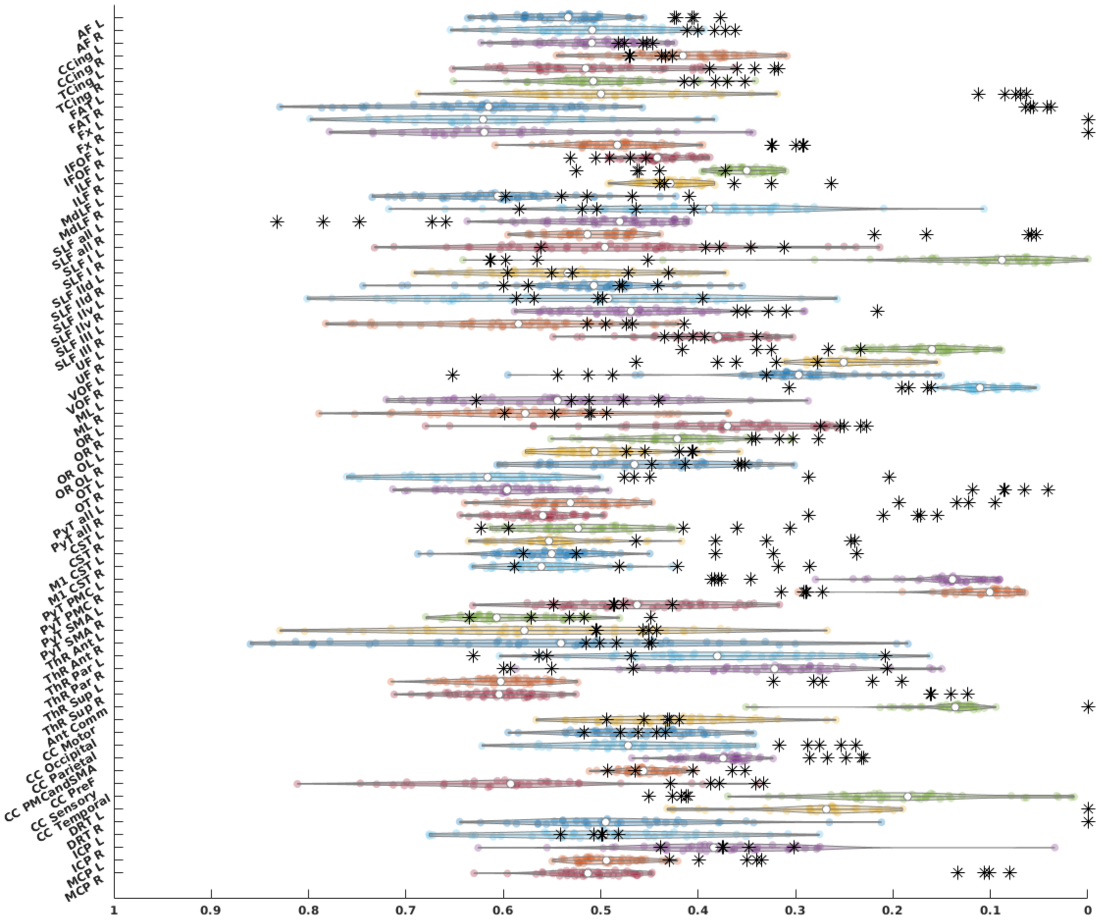


**Supplementary figure 5:** Volume overlap scores for all bundles, HCP test-retest tractograms are depicted as violin plots and MASSIVE results are depicted as black asterisks. AF = arcuate fasciculus, CCing = cingulate cingulum, TCing = temporal cingulum, FAT = frontal aslant tract, Fx = fornix, IFOF = inferior fronto-occipital fasciculu, ILF = inferior longitudinal fasciculus, MdLF = middle longitudinal fasciculus, SLF = superior longitudinal fasciculus, SLF-IId = SLF-II dorsal division, SLF-IIv = SLF-II ventral division, UF = uncinate fasciculus, VOF = vertical occipital fasciculus, ML = medial lemniscus, OR = optic radiation, occlobe = occipital lobe, OT = optic tract, PyT = pyramidal tract, CST = corticospinal tract, M1 = primary motor cortex, PyT = pyramidal tract, PMC = premotor cortex, SMA = supplementary motor area, ThR = thalamic radiation, Ant = anterior, Par = parietal, Sup = superior, Ant Comm = anterior commissure, CC = corpus callosum, PMC and SMA = premotor cortex and supplementary motor area, DRT = dentato-rubro-thalamic tract, ICP = inferior cerebellar peduncle, MCP = middle cerebellar peduncle, L = left, R = right.

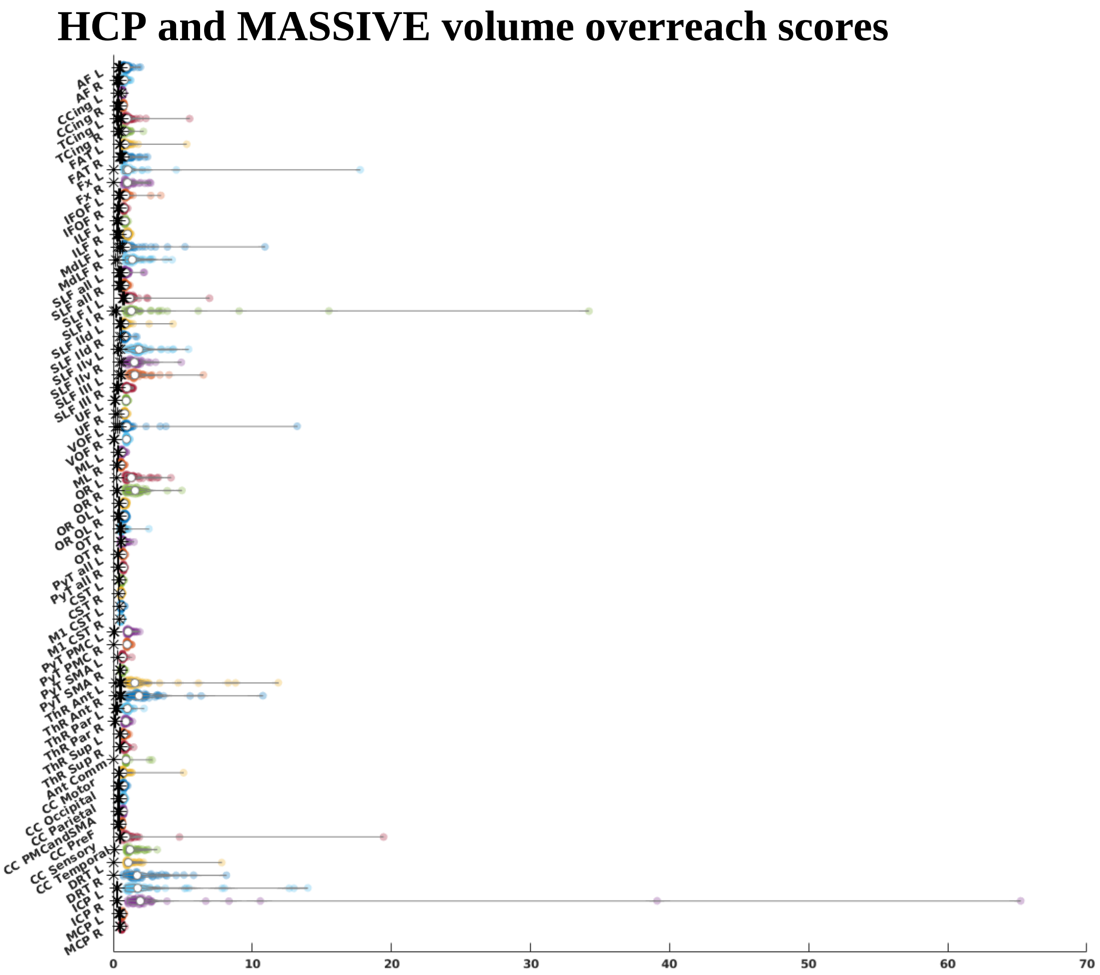


**Supplementary figure 6:** Volume overreach scores for all bundles, HCP test-retest tractograms are depicted as violin plots and MASSIVE results are depicted as black asterisks. AF = arcuate fasciculus, CCing = cingulate cingulum, TCing = temporal cingulum, FAT = frontal aslant tract, Fx = fornix, IFOF = inferior fronto-occipital fasciculu, ILF = inferior longitudinal fasciculus, MdLF = middle longitudinal fasciculus, SLF = superior longitudinal fasciculus, SLF-IId = SLF-II dorsal division, SLF-IIv = SLF-II ventral division, UF = uncinate fasciculus, VOF = vertical occipital fasciculus, ML = medial lemniscus, OR = optic radiation, occlobe = occipital lobe, OT = optic tract, PyT = pyramidal tract, CST = corticospinal tract, M1 = primary motor cortex, PyT = pyramidal tract, PMC = premotor cortex, SMA = supplementary motor area, ThR = thalamic radiation, Ant = anterior, Par = parietal, Sup = superior, Ant Comm = anterior commissure, CC = corpus callosum, PMC and SMA = premotor cortex and supplementary motor area, DRT = dentato-rubro-thalamic tract, ICP = inferior cerebellar peduncle, MCP = middle cerebellar peduncle, L = left, R = right.

9- Maier-Hein, K.H., Neher, P.F., Houde, J.-C., Côté, M.-A., Garyfallidis, E., Zhong, J., Chamberland, M., Yeh, F.-C., Lin, Y.-C., Ji, Q., Reddick, W.E., Glass, J.O., Chen, D.Q., Feng, Y., Gao, C., Wu, Y., Ma, J., Renjie, H., Li, Q., Westin, C.-F., Deslauriers-Gauthier, S., González, J.O.O., Paquette, M., St-Jean, S., Girard, G., Rheault, F., Sidhu, J., Tax, C.M.W., Guo, F., Mesri, H.Y., Dávid, S., Froeling, M., Heemskerk, A.M., Leemans, A., Boré, A., Pinsard, B., Bedetti, C., Desrosiers, M.,

Brambati, S., Doyon, J., Sarica, A., Vasta, R., Cerasa, A., Quattrone, A., Yeatman, J., Khan, A.R., Hodges, W., Alexander, S., Romascano, D., Barakovic, M., Auría, A., Esteban, O., Lemkaddem, A., Thiran, J.-P., Cetingul, H.E., Odry, B.L., Mailhe, B., Nadar, M.S., Pizzagalli, F., Prasad, G., Villalon-Reina, J.E., Galvis, J., Thompson, P.M., Requejo, F.D.S., Laguna, P.L., Lacerda, L.M., Barrett, R., Dell'Acqua, F., Catani, M., Petit, L., Caruyer, E., Daducci, A., Dyrby, T.B., Holland-Letz, T., Hilgetag, C.C., Stieltjes, B., Descoteaux, M., 2017. The challenge of mapping the human connectome based on diffusion tractography. Nature Communications 8, 1349–1349. https://doi.org/10.1038/s41467-017-01285-x

10- Meesters, S., Ossenblok, P., Wagner, L., Schijns, O., Boon, P., Florack, L., Vilanova, A., Duits, R., 2017. Stability metrics for optic radiation tractography: Towards damage prediction after resective surgery. J Neurosci Methods 288, 34–44. https://doi.org/10.1016/j.jneumeth.2017.05.029

11- Pannek, K., Mathias, J.L., Bigler, E.D., Brown, G., Taylor, J.D., Rose, S.E., 2011. The average pathlength map: a diffusion MRI tractography-derived index for studying brain pathology. Neuroimage 55, 133–141. https://doi.org/10.1016/j.neuroimage.2010.12.010

12- Raffelt, D., Tournier, J.-D., Rose, S., Ridgway, G.R., Henderson, R., Crozier, S., Salvado, O., Connelly, A., 2012. Apparent Fibre Density: A novel measure for the analysis of diffusion-weighted magnetic resonance images. NeuroImage 59, 3976–3994. https://doi.org/10.1016/J.NEUROIMAGE.2011.10.045

13- Rheault, F., De Benedictis, A., Daducci, A., Maffei, C., Tax, C.M.W., Romascano, D., Caverzasi, E., Morency, F.C., Corrivetti, F., Pestilli, F., Girard, G., Theaud, G., Zemmoura, I., Hau, J., Glavin, K., Jordan, K.M., Pomiecko, K., Chamberland, M., Barakovic, M., Goyette, N., Poulin, P., Chenot, Q., Panesar, S.S., Sarubbo, S., Petit, L., Descoteaux, M., 2020. Tractostorm: The what, why, and how of tractography dissection reproducibility. Human Brain Mapping 41, 1859–1874. https://doi.org/10.1002/hbm.24917

14- Scilpy documentation [WWW Document], 2021. URL https://scilpy.readthedocs.io/en/latest/ (accessed 2.15.21).

15- Tournier, J.-D., Smith, R.E., Raffelt, D.A., Tabbara, R., Dhollander, T., Pietsch, M., Christiaens, D., Jeurissen, B., Yeh, C.-H., Connelly, A., 2019. MRtrix3: A fast, flexible and open software framework for medical image processing and visualisation. NeuroImage 202, 116137–116137. https://doi.org/10.1016/j.neuroimage.2019.116137
